# A Split-GAL4 screen identifies novel sleep-promoting neurons in the Ventral Nerve Cord of *Drosophila*

**DOI:** 10.1101/2022.02.02.478882

**Authors:** Joseph D. Jones, Brandon L. Holder, Kiran R. Eiken, Alex Vogt, Adriana I. Velarde, Alexandra J. Elder, Jennifer A. McEllin, Stephane Dissel

## Abstract

As in the mammalian system, sleep in *Drosophila* is regulated by multiple brain regions. Among them, neurons projecting to the dorsal Fan-Shaped Body (dFB) have been intensively studied and the data suggest they play a critical role in sleep regulation. The 23E10-GAL4 driver is the most widely used tool to label and manipulate dFB neurons. Multiple studies have reported that activation of 23E10-GAL4 neurons promotes sleep. However, anatomical analyses revealed that 23E10-GAL4 labels 23-30 dFB neurons in the *Drosophila* brain and many non-dFB neurons in the brain and in the Ventral Nerve Cord (VNC), the fly equivalent of the spinal cord. To better understand the role of individual dFB neurons in sleep regulation, we undertook a Split-GAL4 screen to gain access to subsets of 23E10-GAL4 expressing cells. In this study, we report the discovery of two VNC cholinergic sleep-promoting neurons labeled by the 23E10-GAL4 driver.

## Introduction

Understanding the neural basis of behavior is a major aspect of neurobiology. However, unequivocally assigning a behavior to a specific neuron or group of neurons is not a trivial task. The extreme complexity, diversity and connectivity of the mammalian nervous system renders this task even more daunting. To study the neural basis of a behavior, an investigator must be able to specifically manipulate a distinct group of cells and monitor the behavior of interest. Such approaches require precise genetic tools that allow for the manipulation of discrete neurons or groups of neurons. While such tools exist in mammalian systems (1), they may not be available for all the diverse types of neurons that underlie a specific behavior. It is therefore not surprising that animal models have been extensively used to untangle the neural basis of many different behaviors (2). One such model, the fruit fly *Drosophila melanogaster*, with its simpler nervous system and unparalleled genetic toolbox, has become a workhorse in the study of complex behaviors, such as courtship (3, 4), feeding (5, 6), and sleep (7). Multiple binary systems have been developed to access and manipulate specific groups of neurons in the fly nervous system: GAL4/UAS (8); LexA/LexAop (9) and; QF/QUAS (10). Of these 3 systems, the GAL4/UAS system has been by far the most extensively used. However, the expression patterns of GAL4 drivers are often not restricted enough to clearly link a behavior to specific neurons. Importantly, refinement of GAL4 expression pattern can be achieved by employing the intersectional Split-GAL4 technology (11).

Sleep is a behavior that has been described in a variety of species ranging from jellyfish to humans (12). Sleep is regulated by two processes, the circadian clock which gates the occurrence of sleep, and the sleep homeostat which controls the intensity and duration of sleep, in response to prior wakefulness (13). Although the precise function of sleep still remains unknown, there is ample evidence supporting the notion that sleep is required for maintaining optimal physiological and behavioral performance (14). A priori, sleep could appear to be a detrimental activity as it competes with other motivated behaviors such as feeding, mating, or parenting, and renders organisms defenseless against potential predators. However, despite these negative outcomes, sleep has been maintained throughout evolution, emphasizing its essential value (15).

Over the last 20 years, multiple studies have clearly demonstrated that the mechanisms and regulation of sleep are largely conserved from flies to mammals (16). Like in the mammalian system, sleep-regulating centers are found in many areas in the *Drosophila* brain (16, 17). One such sleep-regulating center that has gathered a lot of attention comprises neurons that project to the dorsal Fan-Shaped body (dFB). The Fan-Shaped Body (FB) is part of the central complex in the *Drosophila* brain, a region organized into multiple layers that plays a role in many functions, including locomotion control (18), courtship behavior (19) and memory (20, 21), nociception (22), visual feature recognition (23) and processing (24), social behaviors (25) and, feeding decision-making (26).

Previous studies demonstrated that increasing the activity of neurons contained in the C5-GAL4, 104y-GAL4, and C205-GAL4 drivers strongly promotes sleep (27, 28). While the expression patterns of these 3 independent drivers are broad, they show prominent overlap in neurons that project to the dFB (27). Based on these observations, the authors concluded that it is likely that the dFB plays a role in regulating sleep but could not rule out a role for neurons outside the dFB (27). In addition, further studies established that reducing the excitability of 104y-GAL4 neurons reduces sleep (29, 30). Highlighting the strong interaction between sleep and memory, activation of 104y-GAL4 neurons consolidates Short-Term Memory (STM) to Long-Term Memory (LTM) (27, 31) and restores STM to the classical memory mutant *rutabaga* and *dunce* (32). Taken together, these data point at dFB neurons as important modulator of sleep.

Further work supported a role for the dFB in sleep homeostasis by demonstrating that sleep deprivation increases the excitability of 104y-GAL4 dFB neurons (29). A follow up study proposed that increasing sleep pressure switches dFB neurons from an electrically silent to an electrically active state and that this process is regulated by dopaminergic signaling to the dFB (33) and the accumulation of mitochondrial reactive oxygen species in dFB neurons (34). The physiological properties of dFB neurons have led to the suggestion that dFB neurons are functionally analogous to the ventrolateral preoptic nucleus (VLPO), a key sleep-regulating center in the mammalian brain (29, 35).

Importantly, recent studies have used 23E10-GAL4, a more restrictive driver to manipulate and monitor dFB neurons. (33, 36). Several groups have demonstrated that chronic and acute activation (using thermogenetic or optogenetic methods) of 23E10-GAL4 neurons increases sleep (31, 37–39). Because of its more restricted expression pattern and strong capacity to modulate sleep, 23E10-GAL4 (as it relates to sleep) is seen as a dFB specific driver by most in the scientific community. Remarkably, a recent study described an electron microscopy-based connectome of the central complex including the dFB and demonstrated that the 23E10-GAL4 driver contains 31 neurons in the brain that project to the dFB (40). Whether these 31 dFB neurons can be considered as a functionally homogeneous group is still unclear. Interestingly, Qian *et al.* demonstrated that only a small subset of 23E10-GAL4 neurons express the 5HT2b serotonin receptor and that these cells regulate sleep and sleep homeostasis (37). Unfortunately, this study did not investigate the role of non-5HT2b expressing 23E10- GAL4 neurons in sleep regulation. In addition, whether there are non-dFB 5HT2b expressing 23E10-GAL4 neurons that could potentially play a role in sleep modulation is not clearly addressed in this study either. Taken together, these data suggest that 23E10-GAL4 dFB neurons probably constitute a functionally heterogenous group of cells.

Surprisingly, activation of 23E10-GAL4 expressing neurons, unlike 104y-GAL4, does not consolidate STM to LTM (31). In this study, the authors demonstrated that in addition to the dFB, 104y-GAL4 expresses in neurons projecting to the ventral FB (vFB) and that these neurons also regulate sleep and are responsible for memory consolidation (31). These data emphasize the need to use genetic tools that are as specific as possible to clearly assign a role in sleep regulation and sleep function to specific neurons/groups of neurons. Often, GAL4 drivers that are extensively used to manipulate a specific neuronal group also express in regions outside the area of interest. This makes it difficult to clearly identify the neural basis of sleep behavior.

With this in mind, we wanted to know which of the 31 dFB 23E10-GAL4 neurons are important for sleep regulation. To do so, we employed a targeted, intersectional Split-GAL4 strategy (11) focused on 23E10-GAL4 dFB neurons. Surprisingly, our data demonstrates that in the 23E10-GAL4 driver, there are non-dFB sleep-promoting neurons. Importantly, these cholinergic neurons are located in the Ventral Nerve Cord (VNC), the fly equivalent to the spinal cord. Nevertheless, we confirm that 23E10-GAL4 dFB neurons regulate sleep homeostasis while 23E10-GAL4 VNC neurons do not. Thus, our work reveals that there are multiple groups (at least two) of sleep-regulating neurons in an extensively used “dFB-specific” GAL4 driver. These data clearly highlight the need to use extremely specific tools to manipulate limited number of neurons when trying to unequivocally assign a behavior to a group of cells.

## Results

### The 23E10-GAL4 driver promotes sleep and contains many non dFB neurons in the central nervous system

Previous studies have reported that increasing the activity of dFB neurons using the 23E10-GAL4 driver reliably increases sleep (31, 37–39). We first sought to confirm these observations in our laboratory. Thus, we expressed the thermogenetic TrpA1 cation channel, which is activated by transferring flies to 31°C (41), in 23E10-GAL4 expressing neurons. Individual flies were monitored for two days at 22°C (baseline days, no activation), then temperature was raised to 31°C for 24h on day 3 (activation day). A schematic of the experimental setup is provided in Figure 1A. As seen in Figure 1B,C, raising the temperature changes the sleep profile of both parental controls and results in a loss of sleep (Figure 1E). These data are in agreement with previous reports on the effect of high temperature on sleep (42, 43). On the other hand, activating 23E10-GAL4 neurons strongly increases sleep (Figure 1D,E). Importantly, activation of 23E10-GAL4 neurons does not affect the amount of locomotor activity when the flies are awake (Figure 1F). Thus, we confirmed that activating 23E10-GAL4 neurons promotes sleep and that the increase in sleep is not caused by a general motor defect, in agreement with previous reports using the same GAL4 driver (31, 37–39).

**Figure 1.**
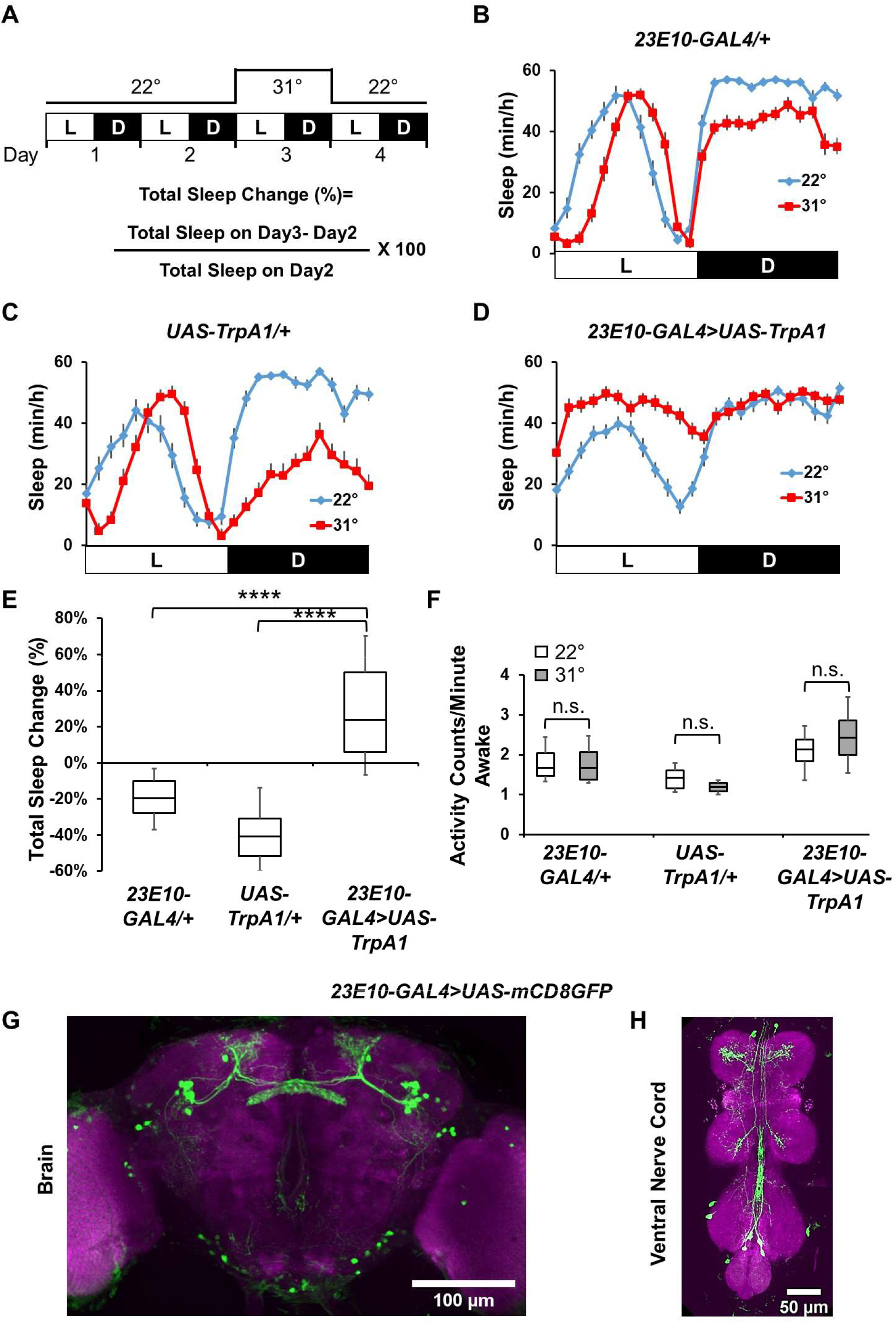
The 23E10-GAL4 driver contains many neurons outside the dFB. **A)** Diagram of the experimental assay. Sleep was measured at 22°C for 2 days to establish baseline sleep profile. Flies were then shifted to 31°C for 24h at the start of day3 to increase activity of the targeted cells by activating the TrpA1 channel, and then returned to 22°C. White bars (L) represent the 12h of light and black bars (D) represent the 12h of dark that are oscillating daily. **B-C)** Sleep profile in minutes of sleep per hour for day 2 (22°C, blue line) and day 3 (31°C, red line) for parental control female flies: *23E10-GAL4/+* (B) and *UAS-TrpA1/+* (C). **D)** Sleep profile in minutes of sleep per hour for day 2 (22°C, blue line) and day 3 (31°C, red line) for *23E10-GAL4>UAS-TrpA1* female flies. **E)** Box plots of total sleep change in % ((total sleep on day3-total sleep on day2/total sleep on day2) X 100) for data presented in B-D. The bottom and top of each box represents the first and third quartile, and the horizontal line dividing the box is the median. The whiskers represent the 10^th^ and 90^th^ percentiles. Flies expressing UAS-TrpA1 in 23E10-GAL4 significantly increase sleep when switched to 31°C compared with parental controls, Kruskal-Wallis ANOVA followed by Dunn’s multiple comparisons. ****P<0.0001, n=25-31 flies per genotype. **F)** Box plots of locomotor activity counts per minute awake for flies presented in B-D. The bottom and top of each box represents the first and third quartile, and the horizontal line dividing the box is the median. The whiskers represent the 10^th^ and 90^th^ percentiles. Two-way repeated measures ANOVA followed by Sidak’s multiple comparisons test found no differences in locomotor activity between 22°C and 31°C. n.s.= not significant. n=25-31 flies per genotype. G-H) Representative confocal stacks of a female *23E10-GAL4>UAS-mCD8GFP* brain (G) and ventral nerve cord (VNC, H). Green, anti-GFP; magenta, anti-nc82 (neuropile marker).

23E10-GAL4 has become the driver of choice to manipulate dFB neurons and many in the field see it as a “dFB-specific” tool. However, GAL4 lines often are not that specific and commonly express in neurons outside the region of interest. To clarify the expression pattern of 23E10-GAL4, we expressed GFP and carefully quantified the number of neurons that are GFP positive. As seen in Figure 1G,H, there are many neurons labelled by 23E10-GAL4 in the brain and in the VNC (See also Video 1 and Video 2). Importantly, many of these neurons are not dFB neurons. Our analysis revealed that there are more than 50 neurons in the brain labelled by 23E10-GAL4, only half of which are dFB neurons (Table1). In addition, 23E10-GAL4 labels about 18 neurons in the VNC (Table1). Interestingly, we note that while the electron microscopy study discussed above identified 31 dFB 23E10-GAL4 neurons (40), we do not systematically observe 31 neurons in every brain. This discrepancy is probably due to the different methodologies used in the two studies. We suspect that some 23E10-GAL4 dFB neurons may be only weakly expressing GAL4 and are therefore not always visible using confocal microscopy. However, based on this expression pattern, we conclude that it is impossible to unequivocally assign a role in sleep promotion to 23E10-GAL4 dFB neurons.

**Table 1:** Quantification of the number of cells labeled by the 23E10-GAL4 driver. Average number ± SEM and range of dFB neurons, all brain neurons and VNC neurons labeled in *23E10-GAL4>UAS-GFP* female flies.

|  | **Average (n)** | **Range** |
| --- | --- | --- |
| **dFB neurons** | 25.68 ± 0.50 (19) | 23-30 |
| **All brain**<br>**neurons** | 51.79 ± 1.05 (19) | 44-62 |
| **VNC neurons** | 17.56 ± 0.99 (9) | 12-22 |

### The 23E10-GAL4 driver contains sleep-promoting neurons that are not dFB neurons

To shed light on the role of individual 23E10-GAL4 expressing neurons in sleep regulation, we employed a Split-GAL4 strategy (11) to access a reduced number of 23E10-GAL4 expressing cells. In designing this screen, our primary region of interest in the brain was the dFB, a choice dictated by the number of studies supporting a role for this structure in sleep regulation (27-30, 33, 34, 37, 44, 45). Central to the Split-GAL4 technology is the fact that the functional GAL4 transcription factor can be separated into two non-functional fragments, a GAL4 DNA-binding domain (DBD) and an activation domain (AD). Different enhancers are used to express these two fragments and the functional GAL4 transcription factor is reconstituted only in the cells in which both enhancers are active (11).

We obtained 22 different AD lines based on their associated GAL4 line expressing in the dFB, as observed using the Janelia Flylight website (36, 46). Since the goal of our screen was to identify the contribution of individual 23E10 dFB neurons in sleep regulation, we designed a targeted approach by combining these individual AD lines to a 23E10-DBD line (46, 47), thus creating 22 new Split-GAL4 lines named FBS for <u>F</u>an-Shaped <u>B</u>ody <u>S</u>plits lines (See Table2 for a description of these lines). These newly created lines were screened behaviorally for their ability to modulate sleep and anatomically to identify their expression patterns. To do so, each individual Split-GAL4 FBS line was crossed to a line expressing both the thermogenetic TrpA1 cation channel and a mCD8GFP construct. Flies were maintained at 22°C for 2 days before raising the temperature to 31°C at the beginning of day 3 for a duration of 24h. Temperature was brought down to 22°C on day 4 (Figure 2A). For each individual fly, we calculated the percentage of total sleep change between activation day (day3) and baseline day (day2) (Figure 2A). As a control for this screen, we used an enhancerless AD construct (created in the vector used to make all the AD lines) combined with 23E10-DBD, since it is the common element in all the Split-GAL4 lines screened in this experiment. As seen in Figure 2B,D, control female flies show a reduction in sleep upon transferring to 31°C, an effect particularly evident during the night. These data are in agreement with previous studies documenting that changes in temperature modulate sleep architecture and that high temperature disturbs sleep at night (42, 43, 48, 49). However, acute activation of the neurons contained in 9 of the 22 FBS lines led to significant increases in percentage sleep change in female flies (FBS4, FBS5, FBS42, FBS45, FBS53, FBS68, FBS72, FBS81 and FBS84), when compared with controls. Individual sleep traces for all 22 FBS lines are shown in Figure 2E,F,G,H and Figure 2-figure supplement 1. Importantly, analysis of activity counts during awake time reveals that increases in sleep are not caused by a reduction in locomotor activity (Figure 2C). On the contrary, 3 of the strongly sleep-inducing lines even show an increase in waking activity upon neuronal activation (Figure 2C). This increase is likely explained by the need for a fly to perform tasks that are mutually exclusive to sleep in a reduced amount of waking time. While increases in total sleep are indicative of increase sleep quantity, this measurement does not provide information about sleep quality or sleep depth. To assess whether sleep quality is modulated by FBS lines, we analyzed sleep consolidation during the day and night in these flies. As seen in Figure 2-figure supplement 2A, upon neuronal activation, daytime sleep bout duration is significantly increased in 8 out of the 9 sleep-promoting lines (FBS4, FBS5, FBS42, FBS45, FBS53, FBS68, FBS72 and FBS84), while it is unaffected in FBS81. Notably, increased sleep bout duration is believed to be an indication of increased sleep depth (50). Thus, these data indicate that activating neurons contained in these 8 FBS lines not only increases sleep level but probably also increases sleep depth. Interestingly, one of the lines that did not show an increase in total sleep, has a significant increase in daytime sleep bout duration (FBS28). During the nighttime, the effect of high temperature on sleep is obvious as most lines, including the control, display a significant decrease in sleep bout duration when raising the temperature to 31°C (Figure 2-figure supplement 2B). Only 6 of the 9 sleep-promoting lines (FBS4, FBS5, FBS42, FBS45, FBS53 and FBS68) maintain similar nighttime sleep bout durations before and during neuronal activation.

**Figure 2:**
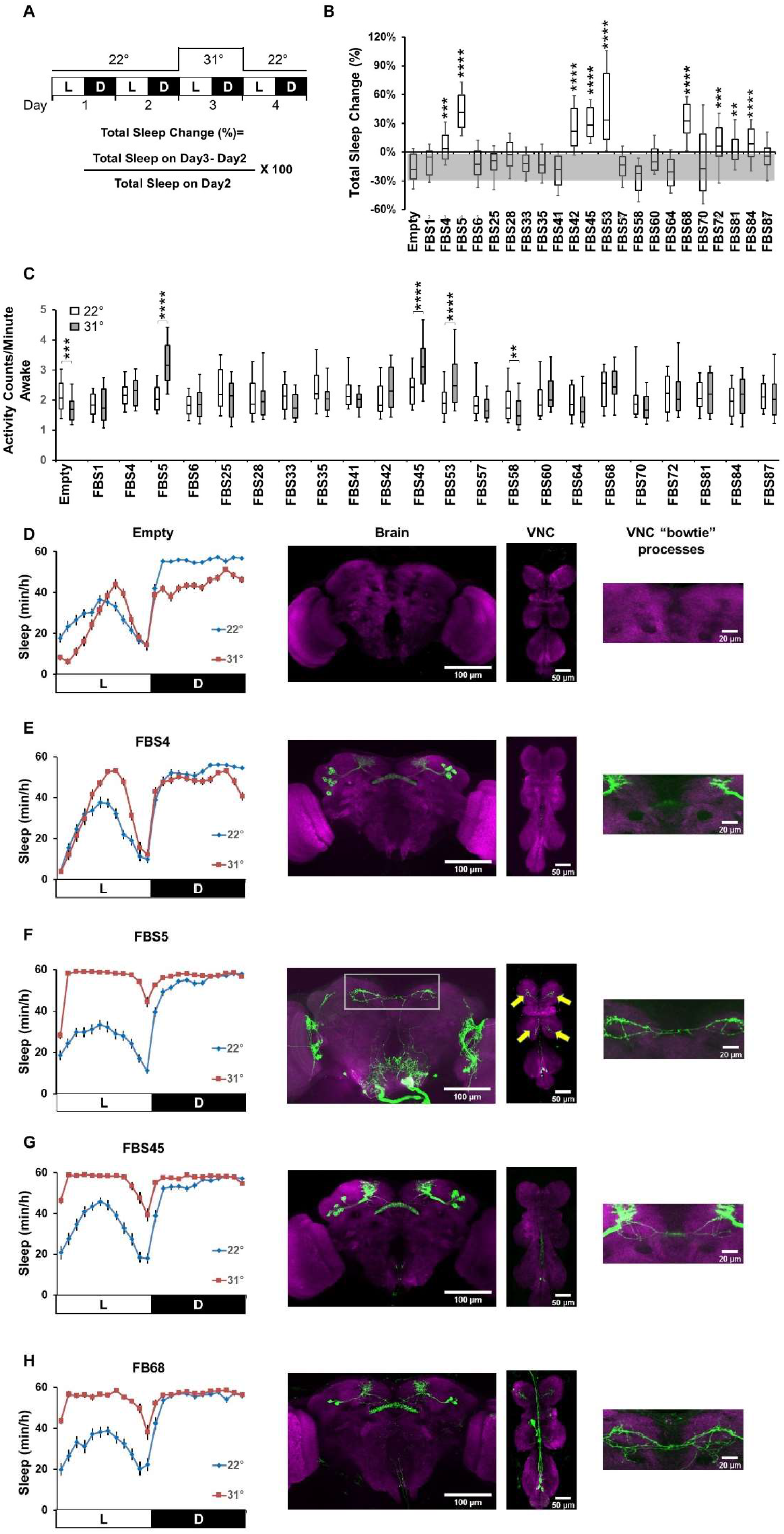
Thermogenetic screen of FBS lines. **A)** Diagram of the experimental assay. Sleep was measured at 22°C for 2 days to establish baseline sleep profile. Flies were then shifted to 31°C for 24h at the start of day3 to increase activity of the targeted cells by activating the TrpA1 channel, and then returned to 22°C. White bars (L) represent the 12h of light and black bars (D) represent the 12h of dark that are oscillating daily. **B)** Box plots of total sleep change in % ((total sleep on day3-total sleep on day2/total sleep on day2) X 100) for female control (Empty-AD; 23E10-DBD) and 22 FBS lines expressing UAS-TrpA1; UAS-GFP. The bottom and top of each box represents the first and third quartile, and the horizontal line dividing the box is the median. The whiskers represent the 10^th^ and 90^th^ percentiles. The gray rectangle spanning the horizontal axis indicates the interquartile range of the control. Kruskal-Wallis ANOVA followed by Dunn’s multiple comparisons revealed that 9 FBS lines increase sleep significantly more than control flies when transferred to 31°C. **P<0.01, ***P<0.001, ****P<0.0001, n=30-51 flies per genotype. **C)** Box plots of locomotor activity counts per minute awake for flies presented in B. The bottom and top of each box represents the first and third quartile, and the horizontal line dividing the box is the median. The whiskers represent the 10^th^ and 90^th^ percentiles. Two-way repeated measures ANOVA followed by Sidak’s multiple comparisons test found that for 3 sleep-promoting FBS lines (FBS5, FBS45 and FBS53) locomotor activity per awake time is increased while no differences are seen for the other 6 sleep-promoting lines between 22°C and 31°C. **P<0.01, ***P<0.001, ****P<0.0001, n=30-51 flies per genotype. D-H) Left, sleep profile in minutes of sleep per hour for day 2 (22°C, blue line) and day 3 (31°C, red line) for empty control (Empty-AD; 23E10-DBD, D), FBS4 (E), FBS5 (F), FBS45 (G) and FBS68 (H) female flies expressing TrpA1 and GFP. Right, Representative confocal stacks for each *FBS>UAS-TrpA1; UAS-mCD8GFP* line of a female brain (left panel), VNC (middle panel) as well as a magnified view of VNC “bowtie” processes as indicated by the gray rectangle in F. Yellow arrows in F indicate TPN1-like processes in FBS5. Green, anti-GFP; magenta, anti-nc82 (neuropile marker).

Importantly, in these 6 lines nighttime sleep bout duration is significantly increased at 31°C when compared to control flies (Figure 2-figure supplement 2B). These data indicate that most sleep-promoting Split-GAL4 lines increase daytime sleep consolidation and suggest that they could also increase nighttime sleep consolidation. However, because of the clear effect of high temperature on nighttime sleep, this claim requires further validation.

In addition, since sleep in *Drosophila* is sexually dimorphic (51–53), we systematically assessed male flies in our experiments. As seen in Figure 2-figure supplement 3, we obtained almost identical behavioral data when analyzing male flies using our thermogenetic activation approach. Thus, we conclude that we have identified 9 FBS lines that promote sleep when thermogenetically activated in males and females.

Having identified 9 different Split-GAL4 lines that significantly increased sleep when activated, we sought to identify the neurons that are contained within these lines. Because the goal of our screen was to identify sleep-regulating dFB neurons in the 23E10-GAL4 pattern of expression, we expected that most of our FBS lines, sleep-promoting or not, would express in dFB cells. This is indeed the case as 20 out of the 22 FBS lines express in some dFB neurons (Figure 2, Figure 2-figure supplement 1, Table 2). The two exceptions to this are FBS25 (expresses in no neurons at all) and FBS5 as discussed below. Regarding sleep-promoting FBS lines, 8 out of 9 have some expression in the dFB. However, and surprisingly, one of the strongly sleep-promoting lines, FBS5, does not express in any dFB neurons at all (Figure 2F, Video 3, Video 4). Rather, FBS5 strongly expresses in two clusters of 4-5 neurons located in the anterior ventrolateral protocerebrum and in 4 VNC cells located in the metathoracic ganglion. A close examination of the anatomy of these 4 VNC neurons indicate that they have processes in the brain. Two of these neurons show a very specific pattern of expression with brain processes located in the superior medial protocerebrum. Because of their characteristic look, we named these processes “bowtie”. These processes are highlighted in the right panel in Figure 2F (see also Video 3). FBS5 additionally labels two neurons in the VNC whose cell bodies are located very close to the “bowtie” neurons. These neurons have characteristic processes in each leg ganglion (yellow arrows in Figure 2F-VNC image). Based on their anatomy, we hypothesize that these cells could be Taste Projection Neurons 1 (TPN1) that receive inputs from sweet gustatory receptor leg neurons (GRNs) and convey this information to the brain in the subesophageal zone (SEZ) (54). We observed these putative TPN1 neurons in 14 out of the 22 FBS lines. Importantly, since all the neurons in FBS5 are part of the 23E10-GAL4 expression pattern, these data indicate that this driver contains non-dFB sleep-promoting cells.

**Table 2:** Description of FBS lines. Identification of the AD construct combined with 23E10-DBD in each FBS line. Average number ± SEM and range of dFB neurons labeled by each line. Bowtie (Y/N) indicates whether “bowtie” processes are seen in the brain, Y= yes, N= no. Range of additional (non-dFB) neurons labeled in the brain of each line. Average number ± SEM of VNC metathoracic cells labeled by each FBS line. TPN1 (Y/N) indicates whether TPN1 neurons are present in the expression pattern, Y= yes, N= no. Range of additional cells in the VNC include any VNC neurons that is not TPN1 or “bowtie” like (VNC-SP).

| **FBS line** | **AD component** | **Average whole brain dFB neurons ± SEM (n)** | **Range dFB neurons** | **Bowtie (Y/N)** | **Range of additional cells in brain** | **Average VNC cells in metathoracic ganglion ± SEM (n)** | **TPN1 (Y/N)** | **Range of additional cells in VNC** |
| --- | --- | --- | --- | --- | --- | --- | --- | --- |
| FBS 1 | 12D12-AD | 4.64 ± 0.40 (n=14) | 2-7 | Y | 0 | 2.00 ± 0.32 (n=11) | N | 0 |
| FBS 4 | 84C10-AD | 20.80 ± 0.35 (n=15) | 18-23 | N | 0 | 0.00 ± 0.50 (n=15) | N | 0 |
| FBS 5 | 23E12-AD | 0 ± 0 (n=26) | 0 | Y | 5-10 | 3.89 ± 0.11 (n=26) | Y | 0 |
| FBS 6 | 42H01-AD | 3.85 ± 0.52 (n=13) | 1-4 | N | 0 | 0.00 ± 0.00 (n=8) | N | 0 |
| FBS 25 | 22A12-AD | 0 ± 0 (n=5) | 0 | N | 0 | 0.00 ± 0.00 (n=5) | N | 0 |
| FBS 28 | 54G07-AD | 13.08 ± 0.63 (n=12) | 10-18 | Y | 0-1 | 3.09 ± 0.17 (n=22) | N | 0 |
| FBS 33 | 49C04-AD | 26.50 ± 1.15 (n=10) | 23-31 | Y | 0 | 3.17 ± 0.32 (n=12) | Y | 0 |
| FBS 35 | 65C03-AD | 5.00 ± 1.06 (n=6) | 2-8 | Y | 1-2 | 3.60 ± 0.24 (n=5) | Y | 0 |
| FBS 41 | VT058545-AD | 11.50 ± 0.43 (n=10) | 10-14 | N | 0-2 | 3.00 ± 0.41 (n=4) | Y | 0 |
| FBS 42 | 12G09-AD | 21.56 ± 0.80 (n=10) | 18-25 | Y | 2-4 | 3.00 ± 0.37 (n=6) | Y | 0-2 |
| FBS 45 | VT050238-AD | 25.11 ± 1.41 (n=9) | 21-32 | Y | 0 | 2.40 ± 0.16 (n=15) | N | 0 |
| FBS 53 | VT043400-AD | 23.86 ± 0.51 (n=7) | 22-26 | Y | 0 | 2.91 ± 0.31 (n=11) | N | 0-1 |
| FBS 57 | VT025720-AD | 7.40 ± 0.68 (n=5) | 5-9 | Y | 0 | 3.00 ± 0.31 (n=7) | Y | 0 |
| FBS 58 | VT064566-AD | 21.50 ± 1.28 (n=6) | 18-27 | Y | 0-2 | 3.10 ± 0.28 (n=10) | Y | 0-1 |
| FBS 60 | VT023818-AD | 24.56 ± 0.50 (n=9) | 22-27 | Y | 0 | 3.27 ± 0.38 (n=11) | Y | 0-2 |
| FBS 64 | VT027956-AD | 15.00 ± 2.12 (n=7) | 11-19 | Y | 0 | 3.40 ± 0.49 (n=10) | Y | 0 |
| FBS 68 | VT023823-AD | 4.45 ± 0.34 (n=11) | 3-6 | Y | 0 | 4.08 ± 0.49 (n=13) | Y | 0-4 |
| FBS 70 | VT034811-AD | 4.86 ± 0.40 (n=7) | 3-6 | Y | 0-7 | 3.90 ± 0.38 (n=10) | Y | 0-4 |
| FBS 72 | VT026953-AD | 24.40 ± 1.50 (n=7) | 20-28 | Y | 0-1 | 3.27 ± 0.25 (n=10) | Y | 0 |
| FBS 81 | 74G10-AD | 22.17 ± 0.70 (n=6) | 19-24 | Y | 0 | 2.80 ± 0.58 (n=5) | N | 0-2 |
| FBS 84 | VT007754-AD | 23.80 ± 0.58 (n=8) | 22-25 | Y | 0-2 | 3.36 ± 0.31 (n=12) | Y | 0-4 |
| FBS 87 | VT045793-AD | 18.55 ± 0.53 (n=13) | 17-22 | Y | 0-2 | 3.62 ± 0.29 (n=13) | Y | 0-6 |

Most other sleep-promoting line expression patterns are very discrete with only two major areas of expression; dFB neurons, and the metathoracic ganglion region containing the two “bowtie” neurons and TPN1-like cells (FBS42, FBS45, FBS53, FBS68, FBS72, FBS81 and FBS84) (Figure 2G,H; Figure 2-figure supplement 1; Table2). Some of these FBS lines show minimal amount of expression elsewhere in the brain, but there is no consistency in these additionally labeled cells between different lines. Of note, FBS4 expresses in 18-23 dFB neurons (Figure 2E, Table 2) and no other cells in the brain or VNC. Considering that 23E10-GAL4 reliably labels 23-30 dFB neurons in our confocal microscopy experiments (Table1), we conclude that FBS4 expresses in 75-80% of 23E10 dFB neurons. Because of its dFB specific expression pattern and its ability to increase sleep when thermogenetically activated, we suspect that FBS4 probably contains neuron(s) that are regulating sleep. However, we also note that the sleep increase triggered by FBS4 thermogenetic activation is not as striking as when activating other strongly sleep-promoting lines like FBS5, FBS53 or FBS68 (Figure 2B for females and Figure 2-figure supplement 3A for males).

One particularly interesting line is FBS45. FBS45 is strongly sleep-promoting and is expressed in 21-32 dFB neurons, indicating that most if not all 23E10 dFB neurons are present in FBS45 (Figure 2G, Table2). However, FBS 45 also expresses in the two “bowtie” neurons. Importantly, FBS45 does not label any other neurons in the brain and VNC.

When combining behavioral and anatomical data, it appears that the numbers of dFB neurons contained within a specific FBS line cannot be predictive of whether the line can promote sleep upon neuronal activation. For example, FBS68 strongly promotes sleep and expresses in only 3-6 dFB neurons (Figure 2H, Table 2), while FBS60 contains 22-27 dFB neurons and does not promote sleep (Figure 2-figure supplement 1L, Table 2).

Remarkably, a careful examination of expression patterns reveals that many FBS lines appear to express in the “bowtie” VNC neurons. Note that the brain processes of these cells are located extremely close to the axonal projections of dFB neurons, making the visualization of these VNC projections sometimes difficult, depending on the strength of dFB projection’s staining. Nevertheless, these VNC “bowtie” neurons are present in 17 out of 22 FBS lines (See Figure 2 and Figure 2-figure supplement 1 VNC “bowtie” processes). Importantly, we can see these neurons and their processes in all sleep-promoting lines with the notable exception of FBS4 (Figure 2, Figure 2-figure supplement 1, Table 2), as discussed above. Upon examination of the literature, we found that neurons with similar projections are frequently seen in many Janelia GAL4 lines (55). This study reports that these neurons (referred to as “sparse T” in this work) and their processes are seen in up to 60% of all Janelia GAL4 lines. The fact that these “bowtie” neurons are frequently present in many GAL4 lines probably explains why we see them in the vast majority of our FBS lines (18 out of 22 or 82% of them).

In conclusion, our thermogenetic screen and anatomical analysis identified 9 sleep-promoting lines with diverse patterns of expression. While the identity of the neurons responsible for sleep promotion in these different lines is not clear at this stage, our analysis indicates that there are multiple groups of sleep-promoting neurons in the 23E10-GAL4 pattern of expression; one of which appears to be part of the dFB (FBS4) while the other is non-dFB based (FBS5).

### Optogenetic confirmation of sleep promoting FBS lines

Because of the strong effect that temperature has on sleep, we sought to confirm our thermogenetic findings using an alternative way to manipulate FBS lines. The logic behind our thinking is that it may be difficult to fully describe and characterize the sleep behaviors of our FBS lines when temperature itself has such a profound effect on sleep. Furthermore, a recent study demonstrated that some of the effects of high temperature on sleep are mediated by GABAergic transmission on dFB neurons (49). Taken together, these data suggest that thermogenetic manipulations may not be optimal to conduct our screen. We thus undertook an optogenetic screen using CsChrimson (56). Our optogenetic experimental setup is described in Figure 3A. Notably, for neurons to be activated when expressing CsChrimson, flies need to be fed all trans-retinal and the neurons must be stimulated with a 627nm LED light source (56). Thus, optogenetic approaches provide a much better control on the timing and parameters of activation.

**Figure 3:**
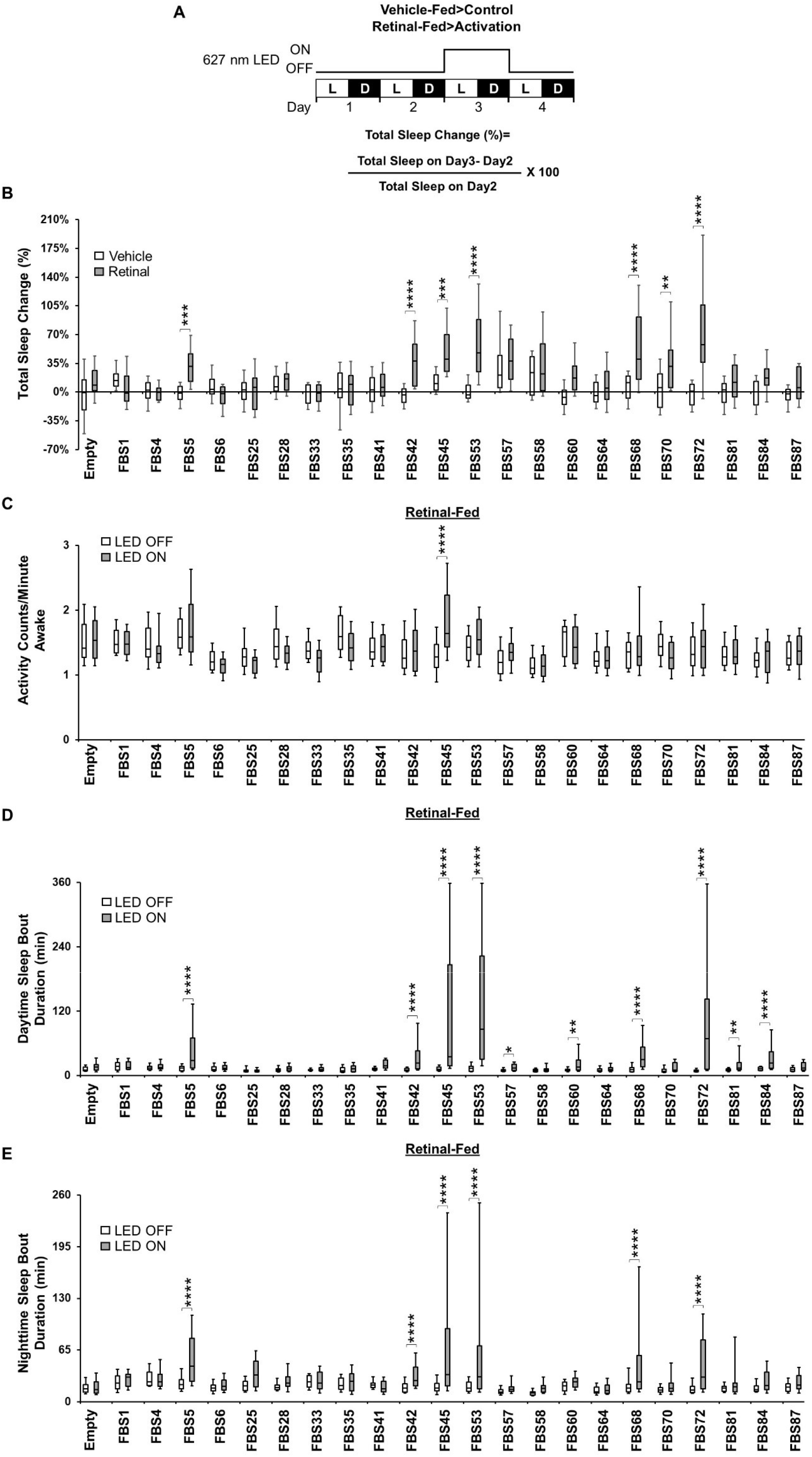
Optogenetic screen of FBS lines. **A)** Diagram of the experimental assay. Sleep was measured in retinal-fed and vehicle-fed flies for 2 days without 627nm LED stimulation to establish baseline sleep profile. LEDs were were then turned on for 24h at the start of day3 to increase activity of the targeted cells by activating the CsChrimson channel, and then turned off on day4. White bars (L) represent the 12h of light and black bars (D) represent the 12h of dark that are oscillating daily. **B)** Box plots of total sleep change in % ((total sleep on day3-total sleep on day2/total sleep on day2) X 100) for control (Empty) and 22 experimental vehicle-fed and retinal-fed female flies expressing CsChrimson upon 627nm LED stimulation. The bottom and top of each box represents the first and third quartile, and the horizontal line dividing the box is the median. The whiskers represent the 10^th^ and 90^th^ percentiles. Two-way ANOVA followed by Sidak’s multiple comparisons revealed that 7 retinal-fed FBS lines increase sleep significantly when stimulated with 627nm LEDs when compared with vehicle-fed flies. **P<0.01, ***P<0.001, ****P<0.0001, n=21-39 flies per genotype and condition. **C)** Box plots of locomotor activity counts per minute awake for retinal-fed flies presented in B. The bottom and top of each box represents the first and third quartile, and the horizontal line dividing the box is the median. The whiskers represent the 10^th^ and 90^th^ percentiles. Two-way repeated measures ANOVA followed by Sidak’s multiple comparisons test found that for most sleep-promoting FBS lines, locomotor activity per awake time is not affected when the flies are stimulated with 627nm LEDs while it is increased in *FBS45>UAS-CsChrimson* flies. ****P<0.0001, n=21-38 flies per genotype. **D)** Box plots of daytime sleep bout duration (in minutes) for retinal-fed flies presented in B. The bottom and top of each box represents the first and third quartile, and the horizontal line dividing the box is the median. The whiskers represent the 10^th^ and 90^th^ percentiles. Two-way repeated measures ANOVA followed by Sidak’s multiple comparisons indicate that daytime sleep bout duration is increased in 10 FBS lines expressing CsChrimson when stimulated with 627nm LEDs. *P<0.05, **P<0.01, ****P<0.0001, n=21-38 flies per genotype. **E)** Box plots of nighttime sleep bout duration (in minutes) for retinal-fed flies presented in B. The bottom and top of each box represents the first and third quartile, and the horizontal line dividing the box is the median. The whiskers represent the 10^th^ and 90^th^ percentiles. Two-way repeated measures ANOVA followed by Sidak’s multiple comparisons indicate that nighttime sleep bout duration is increased in 6 FBS lines expressing CsChrimson when stimulated with 627nm LEDs. ****P<0.0001, n=21-38 flies per genotype.

Nonetheless, optogenetic activation gave results that are mostly identical to the thermogenetic approach. That is, most sleep-promoting lines identified using TrpA1 also induce sleep when activated with CsChrimson in retinal-fed flies (FBS5, FBS42, FBS45, FBS53, FBS68 and FBS72) (Figure 3B for female data). Notable differences are FBS4, FBS81 and FBS84 which do not promote sleep when optogenetically activated in female flies. However, activating FBS81 and FBS84 significantly increases sleep in retinal-fed male flies (Figure 3-figure supplement 2A). Importantly, unlike FBS81 and FBS84, activation of FBS4 neurons does not increase sleep in retinal-fed males (Figure 3-figure supplement 2A). Thus, there is a clear discrepancy for FBS4 comparing thermogenetic and optogenetic activations. Another difference between the two activation approaches is FBS70 that is sleep-promoting using CsChrimson in retinal-fed female flies while it was not using TrpA1 (Figure 3B). Surprisingly, retinal-fed FBS70>UAS-CsChrimson males do not increase sleep upon LED stimulation, potentially revealing a sexually dimorphic phenomenon (Figure 3-figure supplement 2A). Importantly, assessment of locomotor activity during waking period reveals that optogenetic activation does not decrease waking activity in long sleeping retinal-fed females (Figure 3C) and males (Figure 3-figure supplement 2B), ruling out the possibility that locomotor defects are the underlying cause of the sleep phenotypes observed. Examination of sleep bout duration in retinal-fed flies indicates that most sleep-promoting FBS lines increase sleep consolidation during the day (Figure 3D for females and Figure 3-figure supplement 2C for males) and during the night (Figure 3E for females and Figure 3-figure supplement 2D for males). These data indicate that optogenetic activation increases sleep quantity and probably sleep quality (sleep depth) in many of the FBS lines first identified using our thermogenetic approaches.

Interestingly, some FBS lines that did not increase total sleep upon optogenetic activation increased daytime sleep bout duration: FBS57 and FBS60 females (Figure 3D) and FBS33, FBS35, FBS64 and FBS87 males (Figure 3-figure supplement 2C). These data suggest that these lines may also express in sleep-regulating neurons.

Importantly, analysis of sleep parameters in vehicle-fed flies demonstrate that the sleep phenotypes we observed are specific to retinal-fed and LED stimulated flies (Figure 3-figure supplement 1 for females and Figure 3-figure supplement 3 for males).

In conclusion, our thermogenetic and optogenetic neuronal activation revealed several strongly sleep-promoting lines. When looking at their expression patterns, our data clearly demonstrate that 23E10 GAL4 contains at least two different sleep-promoting regions, one that is part of the dFB and one that is not.

### The 23E10-GAL4 driver contains sleep-promoting neurons that are not located in the brain

To reveal the identity of the non-dFB 23E10 sleep-promoting neurons, we undertook a second screen creating additional Split-GAL4 lines focusing on neurons that are present in FBS5. To facilitate this task, we used Color Depth MIP mask search, a software plug-in for Fiji (57). This approach identified several AD lines (containing neurons of interest) that we combined with our 23E10-DBD stock. An example of such a line is shown in Figure 4A-D. This Split-GAL4 line (VT020742-AD;23E10-DBD) expresses only in 4 VNC neurons (Figure 4C). Projections of 2 of these are clearly “bowtie” like (gray arrows in Figure 4A-B). Projections of the other two appear to be identical to TPN1 neurons that have been previously described (yellow arrows in Figure 4A-D). Remarkably, VT020742-AD;23E10-DBD does not express in any other neurons in the brain and VNC. To confirm the presence of TPN1 neurons in this Split-GAL4 line we expressed GFP using a TPN1 specific driver, 30A08-LexA (54) while simultaneously expressing RFP using VT020742-AD;23E10-DBD. As seen in Figure 4-figure supplement 4A, there is clear GFP and RFP co-expression in two out of the four VT020742-AD;23E10-DBD neurons (yellow arrows). In addition, we also see co-expression of GFP and RFP in the processes that go into each leg ganglion, but only RFP expression in the “bowtie” processes. From these data we conclude that VT020742-AD;23E10-DBD labels 4 neurons in the metathoracic ganglion of the VNC: 2 “bowtie” neurons and 2 TPN1 cells. Optogenetic activation of these 4 neurons strongly increases sleep in retinal-fed females (Figure 4K) and males (Figure 4-figure supplement 2A). In addition, daytime and nighttime sleep bout duration are significantly increased in retinal-fed VT020742-AD;23E10-DBD>UAS-CsChrimson flies (Figure 4M-N for females and Figure 4-figure supplement 2C-D for males). Importantly, analysis of waking activity reveals that the sleep increases are not caused by a deficit in locomotor activity (Figure 4L for females and Figure 4-figure supplement 2B for males).

**Figure 4:**
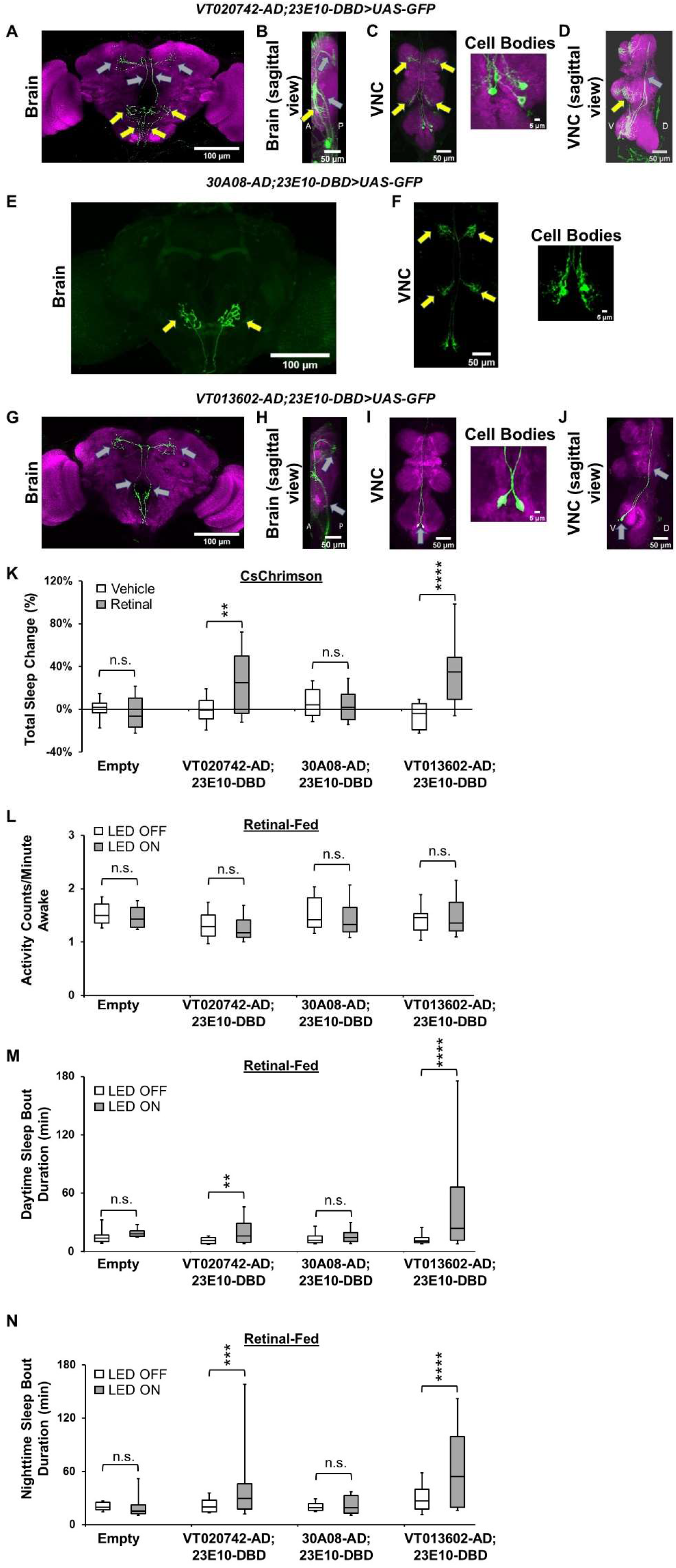
The 23E10-GAL4 driver contains sleep-promoting neurons that are located in the VNC. **A-D)** Representative confocal stacks of a female *VT020742-AD; 23E10-DBD>UAS-GFP* brain (A), brain sagittal view (B), VNC (C) and VNC sagittal view (D). The cell bodies panel in C is a magnified view of the metathoracic area of the VNC. Yellow arrows indicate TPN1 like processes and gray arrows “bowtie” processes. Green, anti-GFP; magenta, anti-nc82 (neuropile marker). A= anterior, P= posterior, V= ventral and D= dorsal. **E-F)** Representative confocal stacks of a female *30A08-AD; 23E10-DBD>UAS-GFP* brain (E) and VNC (F). The cell bodies panel in F is a magnified view of the metathoracic area of the VNC. Yellow arrows indicate TPN1 processes in the brain and VNC. Green, anti-GFP. **G-J)** Representative confocal stacks of a female *VT013602-AD; 23E10-DBD>UAS-GFP* brain (G), brain sagittal view (H), VNC (I) and VNC sagittal view (J). The cell bodies panel in I is a magnified view of the metathoracic area of the VNC. Gray arrows indicate “bowtie” processes and cell bodies in I and J. Green, anti-GFP; magenta, anti-nc82 (neuropile marker). A= anterior, P= posterior, V= ventral and D= dorsal. **K)** Box plots of total sleep change in % for control (*Empty-AD; 23E10-DBD>UAS-CsChrimson), VT020742-AD; 23E10-DBD>UAS-CsChrimson, 30A08-AD; 23E10-DBD>UAS-CsChrimson and VT013602-AD; 23E10-DBD>UAS-CsChrimson* vehicle-fed and retinal-fed female flies upon 627nm LED stimulation. The bottom and top of each box represents the first and third quartile, and the horizontal line dividing the box is the median. The whiskers represent the 10^th^ and 90^th^ percentiles. Two-way ANOVA followed by Sidak’s multiple comparisons revealed that retinal-fed *VT020742-AD; 23E10-DBD>UAS-CsChrimson* and *VT013602-AD; 23E10-DBD>UAS-CsChrimson* flies increase sleep significantly when stimulated with 627nm LEDs when compared with vehicle-fed flies. **P<0.01, ****P<0.0001, n.s.= not significant. n=13-34 flies per genotype and condition. **L)** Box plots of locomotor activity counts per minute awake for retinal-fed flies presented in K. The bottom and top of each box represents the first and third quartile, and the horizontal line dividing the box is the median. The whiskers represent the 10^th^ and 90^th^ percentiles. Two-way repeated measures ANOVA followed by Sidak’s multiple comparisons test shows that locomotor activity per awake time is not affected when the flies are stimulated with 627nm LEDs. n.s.= not significant. n=15-34 flies per genotype. **M)** Box plots of daytime sleep bout duration (in minutes) for retinal-fed flies presented in K. The bottom and top of each box represents the first and third quartile, and the horizontal line dividing the box is the median. The whiskers represent the 10^th^ and 90^th^ percentiles. Two-way repeated measures ANOVA followed by Sidak’s multiple comparisons indicates that daytime sleep bout duration is increased in retinal-fed *VT020742-AD; 23E10-DBD>UAS-CsChrimson* and *VT013602-AD; 23E10-DBD>UAS-CsChrimson* flies when stimulated with 627nm LEDs. **P<0.01, ****P<0.0001, n.s.= not significant. n=15-34 flies per genotype. **N)** Box plots of nighttime sleep bout duration (in minutes) for retinal-fed flies presented in K. The bottom and top of each box represents the first and third quartile, and the horizontal line dividing the box is the median. The whiskers represent the 10^th^ and 90^th^ percentiles. Two-way repeated measures ANOVA followed by Sidak’s multiple comparisons indicates that nighttime sleep bout duration is increased in retinal-fed *VT020742-AD; 23E10-DBD>UAS-CsChrimson* and *VT013602-AD; 23E10-DBD>UAS-CsChrimson* flies when stimulated with 627nm LEDs. ***P<0.001, ****P<0.0001, n.s.= not significant. n=15-34 flies per genotype.

We then wanted to find out whether TPN1 neurons are the cells involved in sleep promotion in VT020742-AD;23E10-DBD. We thus created a 30A08-AD;23E10-DBD Split-GAL4 line. This line labels only the two TPN1 neurons and no other cells in the brain and VNC (Figure 4E-F). Importantly, optogenetic activation of 30A08-AD;23E10-DBD neurons has no effect on sleep (Figure 4K-N and Figure 4-figure supplement 2A-D), indicating that the sleep-promoting capacity of VT020742-AD;23E10-DBD most likely lies within the two “bowtie” neurons. To further rule out an involvement of TPN1 neurons in sleep promotion, we adopted the Split-GAL4 repressor Killer Zipper (KZip+) technology which inhibits an AD and DBD from forming a functional GAL4 (58).

Expressing KZip+ with 30A08-LexA while expressing GFP with VT020742-AD;23E10-DBD, reduces the number of GFP positive cells from 4 to 2 (Figure 4-figure supplement 4B). Only the two “bowtie” neurons can be seen in this combination. Thus, we can effectively subtract the TPN1 neurons from the VT020742-AD;23E10-DBD pattern of expression. Optogenetic activation of the remaining “bowtie” neurons (in retinal-fed VT020742-AD;23E10-DBD>UAS-CsChrimson, 30A08-LexA>LexAop2-KZip+ flies) still promotes sleep (Figure 4-figure supplement 4C). Taken together these data indicate that the two “bowtie” neurons in VT020742-AD;23E10-DBD (and 23E10-GAL4) increase sleep when activated and that TPN1 neurons are not involved in sleep promotion.

To further refine our analysis, we constructed an additional Split-GAL4 line (VT013602-AD;23E10-DBD) that only expresses in the two “bowtie” like VNC neurons (gray arrows in Figure 4G-J, Video 5, Video 6). Optogenetic activation of retinal-fed VT013602-AD;23E10-DBD neurons strongly promotes sleep (Figure 4K and Figure 4-figure supplement 2A for male data) and increases daytime (Figure 4M and Figure 4-figure supplement 2C) and nighttime bout duration (Figure 4N and Figure 4-figure supplement 2D). Again, increases in sleep are not caused by abnormal locomotor activity (Figure 4L and Figure 4-figure supplement 2B). Importantly, analysis of sleep parameters in vehicle-fed flies reveal sleep phenotypes observed are specific to activation of VT013602-AD; 23E10-DBD neurons (Figure 4-figure supplement 1 and Figure 4-figure supplement 3). Since activating the two ‘bowtie” neurons increases daytime and nighttime sleep bout duration, we hypothesize that this manipulation increases sleep quality, namely sleep depth. To investigate this possibility, we measured arousal thresholds in VT013602-AD;23E10-DBD>UAS-CsChrimson flies. As seen in Figure 4-figure supplement 5, the proportion of retinal-fed flies awakened by mild mechanical perturbations (1 and 2) is significantly reduced in LED stimulated (LED ON, activated) individuals compared with non-activated flies (LED OFF). Thus, these data demonstrate that when activated, the two “bowtie” neurons increase sleep quantity and sleep depth as indicated by an increase in arousal threshold. Importantly, at the highest perturbation level (4), retinal-fed LED activated VT013602-AD;23E10-DBD>UAS-CsChrimson flies respond to the stimulus similarly to controls, indicating that the sleep that is induced by activation of these neurons is reversible.

To assess whether the “bowtie” neurons seen in VT020742-AD;23E10-DBD and VT013602-AD;23E10-DBD are the same, we constructed a line that carries both VT020742-AD and VT013602-AD as well as 23E10-DBD. Expressing GFP with this line labels only 4 VNC neurons, 2 “bowtie” cells and 2 TPN1 (Figure 4-figure supplement 4D). If the “bowtie” neurons were different in VT020742-AD;23E10-DBD and VT013602-AD;23E10-DBD we would expect to see 6 cells. Since this is not the case, we can conclude that the 2 ‘bowtie” neurons observed in VT020742-AD;23E10-DBD and VT013602-AD;23E10-DBD are the same. To provide functional support to this finding, we created an additional Split-GAL4 line, VT013602-AD;VT020742-DBD. This Split-GAL4 line labels 2 “bowtie” cells in the VNC with typical processes in the brain (gray arrows in Figure 4-figure supplement 4E) and promotes sleep when activated thermogenetically (Figure 4-figure supplement 4F-H). Taken together, these data demonstrate that we have unequivocally identified two novel sleep-promoting neurons located in the VNC. Importantly, these cells are part of the 23E10-GAL4 expression pattern. We will from now on refer to these neurons as VNC-SP (for VNC Sleep-Promoting).

### Silencing VNC-SP neurons reduces sleep

Having demonstrated that activation of VNC-SP neurons increases sleep duration and sleep depth, we wondered whether these cells could modulate sleep bi-directionally. Thus we performed experiments to silence the activity of VNC-SP neurons either chronically, by expressing the inward rectifying potassium channel Kir2.1 (59) or acutely by using the temperature sensitive Shi^ts1^ construct (60). As seen in Figure 5A-B, total sleep is significantly reduced when expressing Kir2.1 in VNC-SP neurons in female flies. This reduction in sleep is accompanied by a reduction of both daytime (Figure 5C) and nighttime sleep bout duration (Figure 5D), suggesting a defect in sleep consolidation. Importantly, analysis of activity counts during waking time reveals that the decrease in sleep is not due to hyperactivity (Figure 5E). We obtained identical behavioral results when expressing Kir2.1 in VNC-SP neurons of male flies (Figure 5F-J). Thus, chronic silencing of VNC-SP neurons decreases sleep and disrupts sleep consolidation.

**Figure 5:**
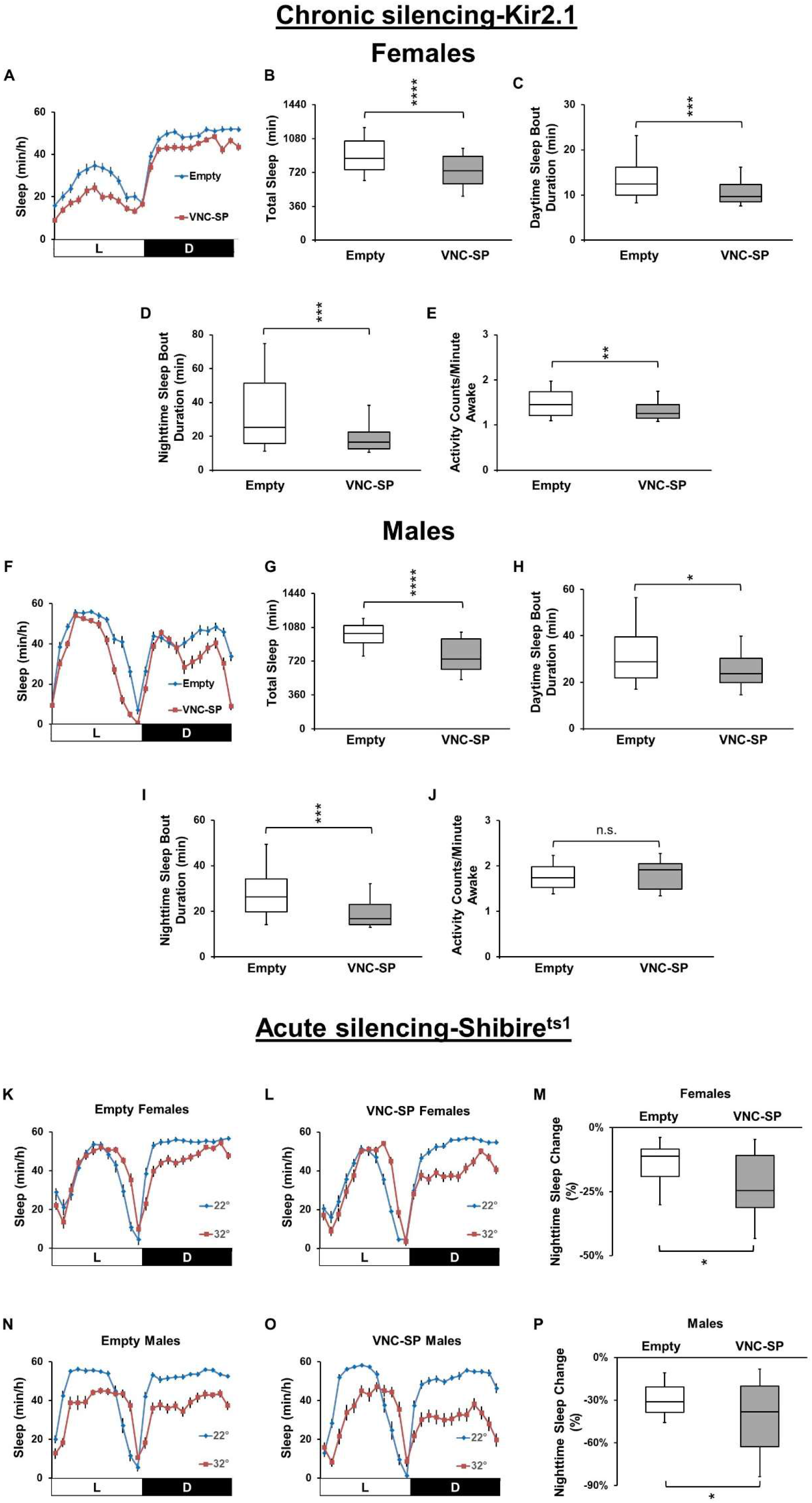
Silencing VNC-SP neurons reduces sleep. **A)** and F) Sleep profile in minutes of sleep per hour for control *(Empty-AD; 23E10-DBD>UAS-Kir2.1, blue line)* and *VNC-SP>Kir2.1 (VT013602-AD; 23E10-DBD>UAS-Kir2.1*, red line) female (A) and male (F) flies. **B)** and G) Box plots of total sleep time (in minutes) for flies presented in A and F, respectively. The bottom and top of each box represents the first and third quartile, and the horizontal line dividing the box is the median. The whiskers represent the 10^th^ and 90^th^ percentiles. A two-tailed unpaired t test revealed that total sleep is significantly reduced in *VNC-SP>Kir2.1* female flies compared to controls (B). A two-tailed Mann-Whitney U test revealed that total sleep is significantly reduced in *VNC-SP>Kir2.1* male flies compared to controls (G). ****P<0.0001. n=58-60 flies per genotype for females and n=48 flies per genotype for males. **C)** and H) Box plots of daytime sleep bout duration (in minutes) for flies presented in A and F, respectively. The bottom and top of each box represents the first and third quartile, and the horizontal line dividing the box is the median. The whiskers represent the 10^th^ and 90^th^ percentiles. Two-tailed Mann-Whitney U tests revealed that daytime sleep bout duration is significantly reduced in *VNC-SP>Kir2.1* female (C) and male (H) flies compared to controls. *P<0.05, ***P<0.0001. n=58-60 flies per genotype for females and n=48 flies per genotype for males. **D)** and I) Box plots of nighttime sleep bout duration (in minutes) for flies presented in A and F, respectively. The bottom and top of each box represents the first and third quartile, and the horizontal line dividing the box is the median. The whiskers represent the 10^th^ and 90^th^ percentiles. Two-tailed Mann-Whitney U tests revealed that nighttime sleep bout duration is significantly reduced in *VNC-SP>Kir2.1* female (D) and male (I) flies compared to controls. ***P<0.001. n=58-60 flies per genotype for females and n=48 flies per genotype for males. **E)** and J) Box plots of locomotor activity counts per minute awake for flies presented in A and F, respectively. The bottom and top of each box represents the first and third quartile, and the horizontal line dividing the box is the median. The whiskers represent the 10^th^ and 90^th^ percentiles. Two-tailed Mann-Whitney U tests revealed that activity per minute awake is significantly reduced in *VNC-SP>Kir2.1* female (E) while it is unchanged in male (J) flies compared to controls. **P<0.001, n.s.= not significant. n=58-60 flies per genotype for females and n=48 flies per genotype for males. **K)** and N) Sleep profile in minutes of sleep per hour for control (*Empty-AD; 23E10-DBD>UAS-Shi*^ts1^) female (K) and male (N) flies at 22°C (blue line) and 32°C (red line). **L)** and O) Sleep profile in minutes of sleep per hour for *VNC-SP>UAS-Shi*^ts1^ female (L) and male (O) flies at 22°C (blue line) and 32°C (red line). **M)** and P) Box plots of nighttime sleep change in % for female flies presented in K and L (M) and male flies presented in N and O. The bottom and top of each box represents the first and third quartile, and the horizontal line dividing the box is the median. The whiskers represent the 10^th^ and 90^th^ percentiles. Two-tailed unpaired t tests revealed that *VNC-SP>Shi*^ts1^ flies lose significantly more sleep when transferred to 32°C compared with controls in both males and females. *P<0.05. n=27-31 flies per genotype for females and n=31 flies per genotype for males.

To confirm the findings obtained using chronic silencing we decided to employ an acute silencing protocol by expressing Shi^ts1^ (60) in VNC-SP neurons. Notably, this approach relies on increasing temperature to silence neurons. Because of the previously reported effect of temperature on sleep, it is not surprising to see that the shift in temperature reduces sleep in control flies (Figure 5K for females and Figure 5N for males). Acute silencing of VNC-SP neurons also reduces sleep, an effect particularly striking at night (Figure 5L and Figure 5M). Importantly, nighttime sleep is significantly more reduced in VNC-SP> Shi^ts1^ flies than in controls (Figure 5M and Figure 5P).

Taken together, our chronic and acute silencing data demonstrate that VNC-SP neurons positively regulate sleep. Importantly, these results further reinforce our findings using optogenetic and thermogenetic activation experiments.

### VNC-SP neurons express and use acetylcholine to modulate sleep

Having demonstrated that VNC-SP neurons modulate sleep bi-directionally, we sought to identify the neurotransmitter(s) used by these cells. A previous study suggested that most neurons in the VNC use one of three neurotransmitters: GABA, Glutamate or Acetylcholine (61). With this in mind, we expressed GFP in VNC-SP neurons and incubated them with antibodies to GABA, choline acetyltransferase (ChAT), the enzyme necessary to produce Acetylcholine and the vesicular glutamate transporter (VGlut). As seen in Figure 6A, we observed clear co-labelling of GFP and ChAT in VNC-SP neurons. To further confirm that VNC-SP neurons express ChAT, we employed a combinatorial approach expressing RFP in VNC-SP cells and GFP driven by a ChAT-LexA (62) construct that is inserted within the endogenous ChAT locus. This construct ensures that the expression of LexA is controlled by ChAT regulatory sequences and should reflect transcriptional activity at the endogenous ChAT locus. As seen in Figure 6-figure supplement 1A, we observe clear co-labelling of GFP and RFP in VNC-SP neurons (gray arrow). In addition, we employed a similar approach using 23E10-GAL4 to drive expression of RFP. As seen in Figure 6-figure supplement 1B, we observe clear co-labelling of GFP and RFP in two neurons in the metathoracic ganglion of the VNC (gray arrows). The evidence clearly suggests that these two cells are the VNC-SP neurons. Interestingly, in the process of this experiment, we found that 2-3 dFB neurons per brain hemisphere are co-labelled by 23E10-GAL4 and ChAT-LexA construct (Figure 6-figure supplement 1B, bottom panels, red arrows). Taken together, these data indicate that VNC-SP neurons and some 23E10-GAL4 dFB neurons are cholinergic.

**Figure 6:**
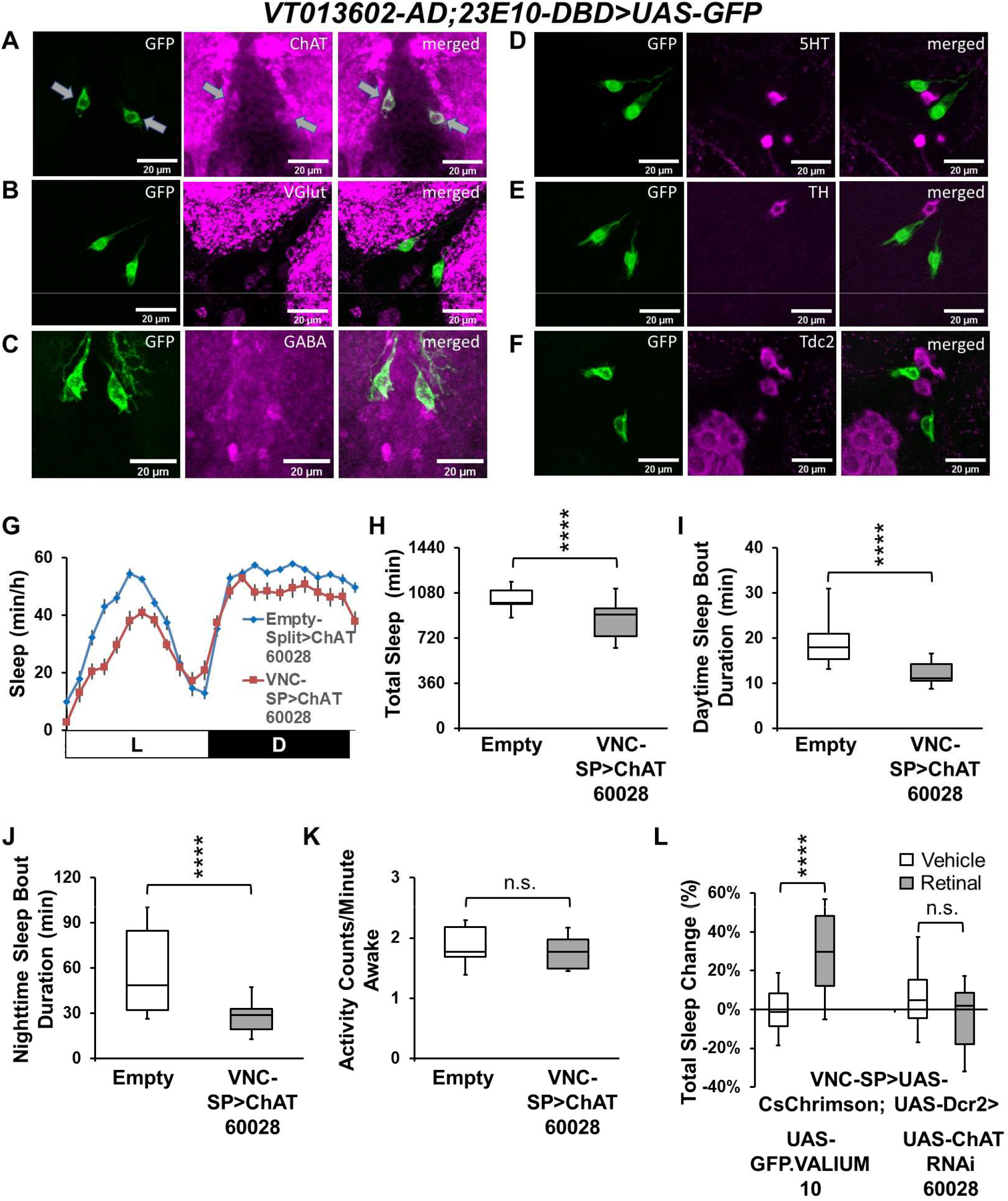
VNC-SP neurons are cholinergic. **A-F)** Representative confocal stacks focusing on the metathoracic ganglion of the VNC of female *VT013602-AD; 23E10-DBD>UAS-GFP* flies stained with antibodies to ChAT (A), VGlut (B), GABA (C), 5HT (D), TH (E) and Tdc2 (F). Gray arrows in A indicate colocalization of GFP and ChAT staining in VNC-SP neurons. Green, anti-GFP; magenta, anti-ChAT (A), anti-VGlut (B), anti-GABA (C), anti-5HT (D), anti-TH (E) and anti-Tdc2 (F). **G)** Sleep profile in minutes of sleep per hour for control (*Empty-AD; 23E10-DBD>UAS-ChAT*^RNAi 60028^, blue line) and *VNC-SP>ChAT*^RNAi^ (*VT013602-AD; 23E10-DBD> UAS-ChAT*^RNAi 60028^, red line) female flies. **H)** Box plots of total sleep time (in minutes) for flies presented in G. The bottom and top of each box represents the first and third quartile, and the horizontal line dividing the box is the median. The whiskers represent the 10^th^ and 90^th^ percentiles. A two-tailed unpaired t test revealed that *VNC-SP> ChAT*^RNAi^ flies sleep significantly less than controls. ****P<0.0001. n=22-30 flies per genotype. **I)** Box plots of daytime sleep bout duration (in minutes) for flies presented in G. The bottom and top of each box represents the first and third quartile, and the horizontal line dividing the box is the median. The whiskers represent the 10^th^ and 90^th^ percentiles. A two-tailed Mann-Whitney U test revealed that *VNC-SP> ChAT*^RNAi^ flies daytime sleep bout duration is significantly reduced compared with controls. ****P<0.0001. n=22-30 flies per genotype. **J)** Box plots of nighttime sleep bout duration (in minutes) for flies presented in G. The bottom and top of each box represents the first and third quartile, and the horizontal line dividing the box is the median. The whiskers represent the 10^th^ and 90^th^ percentiles. A two-tailed Mann-Whitney U test revealed that *VNC-SP> ChAT*^RNAi^ flies nighttime sleep bout duration is significantly reduced compared with controls. ****P<0.0001. n=22-30 flies per genotype. **K)** Box plots of locomotor activity counts per minute awake for flies presented in G. The bottom and top of each box represents the first and third quartile, and the horizontal line dividing the box is the median. The whiskers represent the 10^th^ and 90^th^ percentiles. A two-tailed Mann-Whitney U test revealed that there is no difference in waking activity between *VNC-SP> ChAT*^RNAi^ flies and controls. n.s= not significant. n=22-30 flies per genotype. **L)** Box plots of total sleep change in % for vehicle-fed and retinal-fed *VT013602-AD; 23E10-DBD>UAS-Cs Chrimson*; *UAS-GFP.VALIUM10* (control) and *VT013602-AD; 23E10-DBD>UAS-Cs Chrimson; UAS-ChAT*^RNAi 60028^ flies stimulated by 627nm LEDs. The bottom and top of each box represents the first and third quartile, and the horizontal line dividing the box is the median. The whiskers represent the 10^th^ and 90^th^ percentiles. A two-way ANOVA followed by Sidak’s multiple comparisons shows that sleep is significantly increased in control flies and that expressing ChAT RNAi in VNC-SP neurons completely abolishes the sleep-promoting effect of VNC-SP neurons activated by 627nm LEDs. ****P<0.0001, n.s= not significant. n=31-36 flies per genotype and condition.

Importantly, we did not detect any co-labelling between GFP and VGlut (Figure 6B) or GABA (Figure 6C) in these experiments. To find out whether VNC-SP neurons express any additional neurotransmitters we also performed experiments using antibodies to serotonin (5HT, Figure 6D), tyrosine hydroxylase (TH, Figure 6E), the key enzyme in the synthesis of dopamine, and tyrosine decarboxylase 2 (Tdc2, Figure 6F), an enzyme involved in the synthesis of octopamine. We did not observe any co-labelling with any of these 3 antibodies.

These data indicate that VNC-SP neurons are cholinergic but they do not provide any information as to whether cholinergic transmission is needed for the sleep modulating role of VNC-SP neurons. To address this question, we sought to disrupt cholinergic transmission in VNC-SP neurons using an RNAi approach. Thus, we obtained two independent RNAi lines for ChAT. We first sought to validate them by crossing them to the pan-neuronal elav-GAL4 driver. One of the lines (60028) led to lethality when crossed to elav-GAL4, indicative of a significative reduction in ChAT levels. The second line (25856) gave rise to progeny suggesting it is not very efficient. Note that a similar approach was used to validate ChAT RNAi constructs in other studies (63). However, to further confirm the efficiency of these ChAT RNAi constructs we performed a qPCR analysis on flies expressing ChAT RNAi (60028) specifically in the eyes using a GMR-GAL4 driver. Figure 6-figure supplement 1C demonstrates that ChAT mRNA levels are significantly reduced (about 20%) in the eyes of GMR-GAL4>ChAT RNAi (60028) flies compared with controls. Notably, ChAT mRNA levels are unaffected in the body of these flies, illustrating the efficiency and specificity of the ChAT RNAi line. Importantly, qPCR analysis of the second ChAT RNAi line (25856) showed no significant reduction in ChAT levels, further demonstrating its inefficiency (data not shown). Based on this data, we decided to only employ one ChAT RNAi (60028) in our behavioral studies.

As seen in Figure 6-figure supplement 1D-E, ChAT levels as measured by immunocytochemistry are significantly reduced when expressing ChAT RNAi in VNC-SP neurons. Behaviorally, reducing ChAT levels in VNC-SP neurons significantly reduces sleep (Figure 6G-H) and, daytime (Figure 6I) and nighttime (Figure 6J) sleep bout duration in female flies. Importantly, the reduction in sleep is not due to hyperactivity (Figure 6K). We obtained similar behavioral data when looking at males (Figure 6-figure supplement 1F-I). Taken together, our data indicate that cholinergic transmission is involved in regulating the sleep modulating effect of VNC-SP neurons. We also note that the behavioral effects of reducing ChAT levels and silencing VNC-SP neurons are consistent with each other, suggesting that in the absence of ChAT, the ability of VNC-SP neurons to positively regulate sleep is seriously diminished. To further interrogate the necessity of cholinergic transmission for VNC-SP neurons to promote sleep, we activated VNC-SP neurons optogenetically, while simultaneously expressing ChAT RNAi in these cells. As seen in Figure 6L, expression of CsChrimson and a RNAi line targeting GFP (control RNAi) does not prevent sleep increase in retinal-fed flies. However, expressing ChAT RNAi completely abolishes the sleep promotion triggered by activation of VNC-SP neurons. In conclusion, our data clearly demonstrate that VNC-SP neurons are cholinergic and that disruption of cholinergic transmission abolishes the sleep-promoting role of these cells.

To further confirm that the sleep phenotypes we observe are due to VNC-SP neurons, we optogenetically activated VNC-SP cells while expressing the Split-GAL4 repressor KZip+ (58) under the control of a ChAT-LexA driver. As seen in Figure 6-figure supplement 1J, expressing the KZip+ repressor in ChAT expressing neurons blocks the sleep induction caused by activation of VNC-SP neurons. These data strongly suggest that the KZip+ is expressed in VNC-SP neurons and actively represses the binding of the GAL4-AD and GAL4-DBD constructs. To visually confirm repression of GAL4 activity, we expressed GFP in VNC-SP neurons and the KZip+ repressor in ChAT-LexA cells. As seen in Figure 6-figure supplement 1K, expressing the KZip+ repressor in ChAT expressing neurons completely abolishes the expression of GFP seen in VNC-SP neurons. Taken together, these data confirm that the sleep phenotypes we observe are due to activation of VNC-SP neurons and further demonstrate that these cells are cholinergic.

Since VNC-SP neurons are observed in 18 of our 22 FBS lines, and in most sleep-promoting lines (except for FBS4), we wondered whether VNC-SP neurons are the cells responsible for the sleep increase in all our lines. We decided to focus on FBS45, since it expresses in most if not all 23E10-GAL4 dFB neurons (Table 2) and in the two VNC-SP neurons. Importantly, FBS45 is one of the strongest sleep-promoting lines we have studied in this work. Our question was the following: Is the sleep increase caused by activation of FBS45 due to VNC-SP neurons, dFB neurons, or both group of cells? To answer this question, we expressed UAS-CsChrimson in FBS45 neurons while simultaneously driving the expression of the KZip+ repressor in ChAT-LexA cells. We hypothesize that this manipulation should repress GAL4 formation specifically in VNC-SP neurons. However, since we have reported that ChAT-LexA also expresses in some 23E10-GAL4 dFB neurons (Figure 6-figure supplement 1B), our manipulation may repress GAL4 in a few dFB neurons. As seen in Figure 6-figure supplement 1L, expressing the KZip+ repressor in ChAT-LexA prevents retinal-fed FBS45>UAS-CsChrimson flies from increasing sleep when activated. These data indicate that the neurons responsible for the sleep increase when activating FBS45 cells are cholinergic and that expressing CsChrimson only in FBS45 neurons that are not targeted by ChAT-LexA (non-cholinergic) cannot promote sleep. We then visually assessed the extent of ChAT-LexA driven KZip+ repression in FBS45 neurons by expressing UAS-GFP. As seen in Figure 6-figure supplement 1M, expressing the KZip+ repressor in ChAT-LexA completely abolishes GFP expression in VNC-SP neurons in *FBS45>UAS-GFP; ChAT-LexA>LexAop2-KZip+* flies. However, as we expected based on our previous data, ChAT-LexA>LexAop2-KZip+ consistently removes GFP expression in 4-6 dFB neurons as well. Taken together, these data suggest that the sleep-promoting capacity of FBS45 relies on cholinergic neurons. While we strongly believe that the cells responsible are VNC-SP neurons, we cannot completely exclude a role for one of the ChAT expressing 23E10-GAL4 dFB neurons. Further work will help clarify the role of cholinergic dFB neurons in sleep regulation.

Since the VNC receives and integrates sensory inputs from the periphery and sends this information to the brain (64–66), we wondered whether VNC-SP neurons could similarly send sleep-relevant signals to the brain. The location of VNC-SP neurons in the VNC, and their processes in the brain suggests that this is a likely possibility. To find out, we expressed the dendritic marker DenMark (67) in conjunction with the presynaptic marker synaptotagmin-GFP (syt.eGFP) (68) in VNC-SP neurons. Our data indicate that VNC-SP neurons have mostly postsynaptic sites in the VNC, as expected if these cells receive sensory information from the periphery (Figure 6-figure supplement 2, middle panels). However, and surprisingly, the brain processes of these cells are labeled with both the presynaptic and postsynaptic markers (Figure 6-figure supplement 2, top panels). This suggests that VNC-SP neurons are well positioned to receive sensory inputs from the periphery in the VNC and that they are sending this information to the brain. However, since the brain processes of VNC-SP neurons are also harboring postsynaptic sites, it is highly likely that they also get inputs in the brain. Nevertheless, our data suggest that VNC-SP neurons send sleep-relevant signals to the brain.

### Multiple sleep-promoting regions in 23E10-GAL4 with functional specificity

Our data unequivocally demonstrate that VNC-SP neurons, which are part of the 23E10-GAL4 pattern of expression, are sleep-promoting. Since 23E10-GAL4 is widely seen as dFB specific driver when it comes to sleep regulation, the identification of VNC-SP neurons is a major finding that impacts our current views about the role of the dFB in sleep regulation. Our data shows that all but one FBS sleep-promoting lines contain VNC-SP neurons. The only sleep-promoting line that is dFB specific (FBS4) only mildly promotes sleep (when compared with other lines) in our thermogenetic screen but does not using optogenetic. These data raise important questions about the role of the dFB in sleep regulation and sleep-promotion.

To try to address these questions we went back to the 23E10-GAL4 driver and its 23E10-LexA counterpart (69). First, we wanted to check whether 23E10-GAL4 and 23E10-LexA express in the same number of neurons. This is an important control since it was reported that often for a given enhancer sequence, the LexA expression pattern does not fully recapitulate the GAL4 expression pattern (69). We created a line expressing GFP under the control of 23E10-LexA and RFP under the control of 23E10-GAL4. As seen in Figure 7A-B, the 23E10-LexA driver does not fully recapitulate the 23E10-GAL4 pattern of expression in the dFB. Some cells are clearly co-labelled by GFP and RFP (yellow arrows) but some other dFB neurons only express RFP (gray arrow). Importantly, we did not observe any dFB cells (or any other cell in the brain) that are only labeled by 23E10-LexA. These data indicate that, like many other LexA drivers, 23E10-LexA only expresses in a subset of the 23E10-GAL4 pattern of expression. We then turned our attention to the metathoracic ganglion of the VNC. As seen in Figure 7C, 23E10-LexA labels the two TPN1 neurons (yellow arrows) while 23E10-GAL4 labels the two TPN1 cells as well as the two VNC-SP neurons (gray arrows). The processes of TPN1 neurons are highlighted in Figure 7E and those from VNC-SP in Figure 7D. We see clear co-labelling of TPN1 processes by 23E10-LexA and 23E10-GAL4, but VNC-SP processes are only labeled by 23E10-GAL4. Taken together, we conclude that 23E10-LexA only expresses in a subset of 23E10-GAL4 neurons and that 23E10-LexA does not express in VNC-SP neurons.

**Figure 7:**
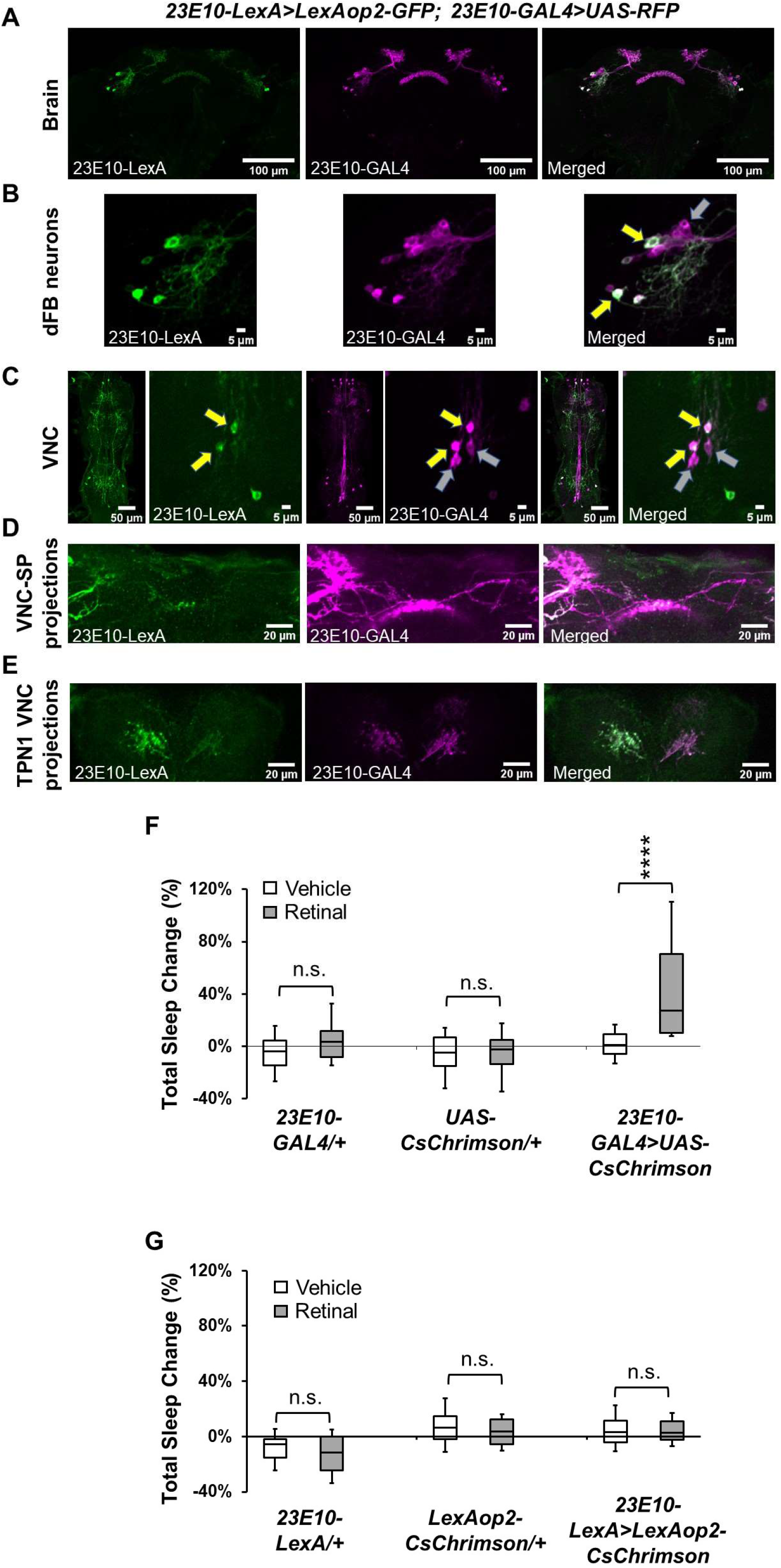
The 23E10-LexA driver does not express in VNC-SP neurons. **A-E)** Representative confocal stacks of a female *23E10-LexA>LexAop2-GFP; 23E10-GAL4>UAS-RFP* fly showing the brain (A), a magnified view of dFB neurons (B), the VNC (C), VNC-SP neurons projections in the brain (D) and TPN1 neurons processes in the VNC (E). Yellow arrows in B show dFB neurons that are labeled by both the 23E10-LexA and 23E10-GAL4 drivers, while the gray arrow shows a neuron that is only labeled by 23E10-GAL4. Our analysis revealed that in these co-labelling experiments, 23E10-LexA expresses in 13-20 dFB neurons while 23E10-GAL4 labels 19-26 dFB cells per brain. Importantly, all the neurons labeled by 23E10-LexA are part of the 23E10-GAL4 expression pattern. Yellow arrows in C show the TPN1 neurons and gray arrows the VNC-SP neurons. Green, anti-GFP; magenta, anti-RFP. **F)** Box plots of total sleep change in % for vehicle-fed and retinal-fed *23E10-GAL4/+ and UAS-CsChrimson/+* parental controls and *23E10-GAL4>UAS-CsChrimson* female flies stimulated by 627nm LEDs. The bottom and top of each box represents the first and third quartile, and the horizontal line dividing the box is the median. The whiskers represent the 10^th^ and 90^th^ percentiles. Two-way ANOVA followed by Sidak’s multiple comparisons revealed that retinal-fed *23E10-GAL4>UAS-CsChrimson* flies significantly increase sleep when stimulated with 627nm LEDs. Both parental controls show no difference. ****P<0.0001, n.s= not significant. n=24-42 flies per genotype and condition. **G)** Box plots of total sleep change in % for vehicle-fed and retinal-fed *23E10-LexA/+* and *LexAop2-CsChrimson/+* parental controls and *23E10-LexA>LexAop2-CsChrimson* female flies stimulated by 627nm LEDs. The bottom and top of each box represents the first and third quartile, and the horizontal line dividing the box is the median. The whiskers represent the 10^th^ and 90^th^ percentiles. A two-way ANOVA followed by Sidak’s multiple comparisons revealed that no differences in sleep following stimulation. n.s= not significant. n=23-31 flies per genotype and condition.

We then wondered whether 23E10-LexA could promote sleep when activated optogenetically. As seen in Figure 7F, optogenetic activation of retinal-fed 23E10-GAL4>UAS-CsChrimson strongly increases sleep as expected. Controls show no changes in sleeping levels. However, optogenetic activation of retinal-fed 23E10-LexA>LexAop2-CsChrimson has no effect on sleep (Figure 7G). The stark difference in behavior seen when activating 23E10-GAL4 and 23E10-LexA could be explained by the fact that 23E10-GAL4 expresses in more neurons than 23E10-LexA. In this scenario the absence of VNC-SP neurons in the 23E10-LexA pattern of expression must account for some if not all this difference. However, since 23E10-LexA does not fully recapitulate the expression pattern of 23E10-GAL4 in the dFB, we cannot rule out that some of these missing dFB neurons may be involved in sleep promotion. An alternative explanation is that 23E10-LexA may not be as strong a driver as 23E10-GAL4 and therefore would not express sufficient levels of the CsChrimson construct. In our experience this is very unlikely as we have optogenetically activated other pairs of Gal4/LexA drivers and obtained identical and strong sleep activation with both the LexA and GAL4 counterparts (data not shown).

Does our data mean that the dFB plays no role at all in sleep regulation? Based on the number of studies supporting a role for this structure in sleep regulation (27-30, 33, 34, 37, 44, 45) we think that this is very unlikely. Most of the previous work supports a role for dFB neurons in sleep homeostasis. In particular, sleep deprivation increases the excitability of dFB neurons (29) and augmented sleep pressure switches dFB neurons from an electrically silent to an electrically active state (33). Furthermore, reducing the excitability of dFB neurons by reducing levels of the Rho-GTPase-activating protein encoded by the crossveinless-c (cv-c) gene leads to a defect in sleep homeostasis (29).

With this in mind, we wanted to assess whether silencing all 23E10-GAL4 neurons, or just 23E10-GAL4 dFB and VNC-SP neurons separately would affect sleep homeostasis (Figure 8A). As seen in Figure 8B, chronic silencing of all 23E10-GAL4 neurons by expression of Kir2.1 effectively blocks homeostatic sleep rebound when compared with both parental controls. Interestingly, chronic silencing of VNC-SP cells does not disrupt homeostatic sleep rebound (Figure 8C) while silencing of most 23E10-GAL4 dFB neurons (contained in the FBS4 line) does (Figure 8D). These data support a role for 23E10-GAL4 dFB neurons in sleep homeostasis, in agreement with previous studies.

**Figure 8:**
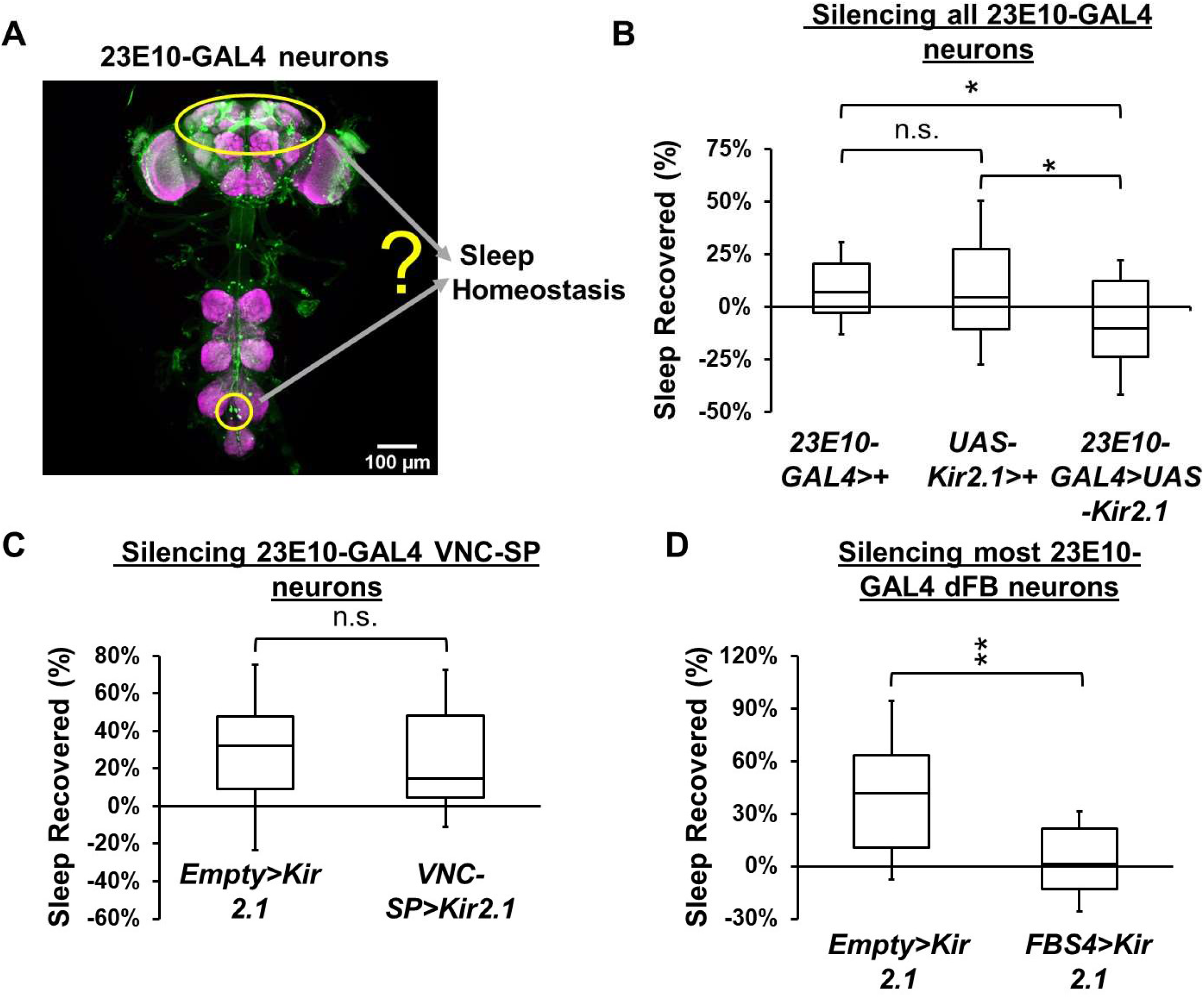
The 23E10-GAL4 contains two groups of sleep regulating neurons. **A)** Representative confocal stacks of a female *23E10-GAL4>UAS-GFP* fly showing dFB neurons and VNC-SP neurons. **B)** Box plots of total sleep recovered in % during the first 24h following 12h of sleep deprivation at night for *23E10-GAL4/+* and *UAS-Kir2.1/+* parental controls and *23E10-GAL4>UAS-Kir2.1* female flies. A one-way ANOVA followed by Tukey’s multiple comparisons demonstrate that both parental controls have a bigger homeostatic sleep rebound than *23E10-GAL4>UAS-Kir2.1* female flies. *P<0.05, n.s= not significant. n=47-62 flies per genotype. **C)** Box plots of total sleep recovered in % during the first 24h following 12h of sleep deprivation at night for *Empty-AD; 23E10-DBD>UAS-Kir2.1* and *VT013602-AD; 23E10-DBD>UAS-Kir2.1* female flies. A two-tailed unpaired t test revealed that there is no difference in homeostatic sleep rebound. n.s= not significant. n=24-29 flies per genotype. **D)** Box plots of total sleep recovered in % during the first 24h following 12h of sleep deprivation at night for *Empty-AD; 23E10-DBD>UAS-Kir2.1* and *FBS4>UAS-Kir2.1* female flies. A two-tailed unpaired t test revealed that *FBS4>UAS-Kir2.1* flies have a significantly reduced homeostatic sleep rebound compared to controls. **P<0.01. n=15-28 flies per genotype.

Taken together, our data indicate that the 23E10-GAL4 driver contains at least two groups of sleep-regulating neurons, one located in the dFB that is mostly involved in sleep homeostasis and VNC-SP neurons that are dispensable for sleep homeostasis.

## Discussion

The strength of the fly model lies in its strong genetic techniques, allowing researchers to selectively manipulate discrete populations of neurons and monitor how specific behaviors are affected. To do so, an impressive collection of binary expression systems have been developed. In particular, the GAL4/UAS system (8) has been the keystone of *Drosophila* neurobiological studies. However, GAL4 drivers are often not specific enough and express in cells outside the region of interest. This can make the task of unequivocally assigning a given behavior to a specific neuron or group of neurons particularly difficult. Importantly, refinement of GAL4 expression pattern can be achieved by employing the intersectional Split-GAL4 technology (11).

Like mammalian sleep, sleep in *Drosophila* is regulated by multiple areas in the brain (16, 17). Particularly, an expanding amount of evidence supports a role for dFB neurons in sleep regulation (27-30, 33, 34, 37, 44, 45). The most widely used tool to manipulate dFB neurons is the 23E10-GAL4 driver (33, 36), and thermogenetic or optogenetic activation of 23E10-GAL4 neurons results in increased sleep (31, 37–39). However, whether 23E10-GAL4 dFB neurons can be considered as a functionally homogeneous group is still unclear. Also unclear is whether the sleep-promotion caused by activation of 23E10-GAL4 neurons is due to dFB neurons. This concern is reinforced by the fact that 23E10-GAL4 expresses in many neurons outside the dFB in the brain and in the VNC (Figure 1, Table 1). In this study, we sought to identify the role of individual 23E10-GAL4 dFB neurons in sleep regulation.

To identify sleep-promoting neurons within the 23E10-GAL4 pattern of expression we adopted a targeted Split-GAL4 strategy combining 22 individual AD lines (selected for the strong dFB expression pattern of their GAL4 counterparts) with a 23E10-DBD line. We screened these 22 novel Split-GAL4 FBS lines behaviorally and anatomically. First, we employed a thermogenetic approach to activate the neurons contained in these 22 lines. These experiments identified 9 FBS lines that promote sleep when activated in females and males (Figure 2). An anatomical assessment of each line revealed that most sleep-promoting FBS lines express in some dFB neurons, as expected based on our targeted approach. However, they also express in other areas in the brain and especially in the metathoracic ganglion of the VNC. In fact, we identified two very clear and characteristic processes associated with 4 different VNC neurons in many FBS lines, sleep-promoting or not. The first set of two VNC neurons have typical processes in each leg ganglion and send processes in the brain that terminate in the SEZ. Our data demonstrated that these cells are TPN1 neurons (Figure 4) (54). We observed these neurons in 14 out of the 22 FBS lines. However, we demonstrated that TPN1 neurons are not promoting sleep when activated (Figure 4). The second set of two VNC neurons send processes to the superior medial protocerebrum. Due to their characteristic anatomical features, we named these processes “bowtie”. Importantly, these processes are seen in 18 out of 22 FBS lines (and in 8 out of the 9 sleep-promoting lines). The high frequency of observation of these processes in our FBS lines suggest that “bowtie” neurons are commonly expressed in many GAL4 lines. In fact, a MultiColor FlpOut (MCFO) study reported that these neurons are observed in more than 60% of all Janelia GAL4 lines (55). Only one line, FBS4, promotes sleep when thermogenetically activated and expresses only in dFB cells, containing about 80% of 23E10-GAL4 dFB neurons. FBS4 can thus be considered as a dFB specific driver. We note that the amount of sleep induction triggered by thermogenetic activation of FBS4 is lower than in most other sleep-promoting FBS lines. However, the most surprising finding from our anatomical analysis is FBS5, a line that has no expression in the dFB, and still promotes sleep when activated. This clearly demonstrates that 23E10-GAL4 contains sleep-promoting neurons that are not dFB cells.

Because temperature has a strong impact on sleep (42, 43, 48, 49), we confirmed our findings using an optogenetic activation method (Figure 3). Most sleep-promoting FBS lines increased sleep when activated using either methods. The notable exception is the dFB specific FBS4 line, which did not affect sleep at all upon optogenetic activation. What is the source of this discrepancy? We can propose several scenarios. The first possible explanation is that the threshold of activation needed for the neurons contained in FBS4 to fire and promote sleep is high. This would imply that thermogenetic activation, which is constant during the 24h activation day, provides stronger activation than the fragmented optogenetic protocol. Considering that dFB neurons are subject to dopaminergic regulation (33), we hypothesize that under baseline conditions, dFB neurons are under severe dopaminergic inhibition and that our optogenetic activation protocol is not sufficient to bring these cells above firing threshold. A second possibility is that FBS4 expresses in neurons underlying the effect of high temperature on sleep. This hypothesis is supported by a recent study demonstrating that some of the effects of high temperature on sleep are mediated by GABAergic transmission on dFB neurons (49). It is thus conceivable, that activating FBS4 neurons thermogenetically may counteract the effect of increasing temperature on sleep. This would explain why we only see a sleep-promoting effect when activating FBS4 using thermogenetic and not optogenetic.

A further screen demonstrated that the VNC ‘bowtie” neurons are sleep-promoting (Figure 4). Activating these cells (renamed VNC-SP neurons) increases sleep, sleep consolidation and sleep depth by increasing arousal thresholds. Moreover, chronic, and acute silencing of VNC-SP neurons reduce sleep and sleep consolidation (Figure 5) confirming that VNC-SP neurons positively regulate sleep. We then sought to discover the neurochemical identity of VNC-SP neurons and identified acetylcholine as a key modulator of sleep in VNC-SP neurons (Figure 6). Reducing cholinergic transmission in VNC-SP neurons reduces sleep and sleep consolidation, an effect mirroring the silencing experiments described earlier. Moreover, we demonstrate that VNC-SP neurons use cholinergic transmission to promote sleep when optogenetically activated.

Since we observed ‘bowtie” processes in most FBS lines, we assume that most of these lines express in VNC-SP neurons. One could therefore wonder why we do not observe total sleep changes when activating these lines? Importantly, our analysis of sleep parameters including day and night sleep bout duration reveals that sleep consolidation is significantly increased using at least one activation protocol with 7 of these “non sleep-promoting” “bowtie” containing lines (FBS28, FBS33, FBS35, FBS57, FBS60, FBS64 and FBS87). Thus, out of the 18 FBS lines that express in VNC-SP neurons, 16 of them increase at least one sleep parameter when activated. The only lines that express in VNC-SP neurons and do not affect any sleep parameters when activated are FBS1 and FBS58. Interestingly, we observed that the GFP signal in the VNC-SP “bowtie” processes of these two lines is weak (Figure 2-figure supplement 1), indicating reduced GAL4 expression in these cells. This observation may provide an explanation for the lack of sleep changes when activating these lines. We also suspect that levels of GAL4 expression in VNC-SP neurons could explain why some FBS lines are promoting sleep more strikingly than others.

Where in the nervous system are these VNC-SP cholinergic signals received? We show that VNC-SP neurons receive synaptic inputs in the VNC and have presynaptic sites in the brain (Figure 6-figure supplement 2). This suggests that VNC-SP neurons are well positioned to receive and integrate sleep-relevant sensory inputs from the periphery in the VNC and that they are sending this information to the brain. The nature of these inputs will require additional investigations that will further uncover the function and role of VNC-SP neurons in sleep regulation. Known modulators of sleep include the immune system and metabolic functions. Based on their location, it is possible that VNC-SP neurons integrate signals relevant to these two sleep-modulating systems.

Our data raised important questions about the role of dFB neurons in sleep regulation. Importantly, we show that 23E10-GAL4 dFB neurons are involved in sleep homeostasis as silencing them completely abrogates sleep rebound following a night of sleep deprivation (Figure 8). Our data is in agreement with previous work supporting a role for dFB neurons in sleep homeostasis (29, 33). Interestingly, we also demonstrate that VNC-SP neurons are not part of the homeostatic sleep circuit. These findings suggest that dFB neurons and VNC-SP neurons regulate different aspects and functions of sleep. Here again, further work will determine the role of VNC-SP neurons.

Finally, our data clearly highlight the need to obtain tools that are as specific as possible when attempting to link a given behavior with a specific neuron or group of neurons. In particular, the frequency at which VNC-SP neurons are observed in existing GAL4 lines (in up to 60% of all Janelia GAL4 lines) (55) raises some important methodological questions. For example, are VNC-SP neurons present in the expression pattern of a GAL4 line that is known to modulate sleep and is seen as a tool specific for a given structure in the brain? Future studies using GAL4 lines in sleep studies will need to take account of the potential presence of VNC-SP neurons in their expression pattern. Perhaps more importantly, existing data may need to be reinterpreted in the light of our finding.

## Conclusions

Our work identified a novel group of cholinergic sleep-promoting neurons located in the VNC in *Drosophila*. We show that these cells are perfectly positioned to receive and integrate sensory inputs from the periphery and send this information to the brain. We demonstrate that these cells regulate sleep quantity and sleep quality. We further demonstrate that they are not involved in sleep homeostasis whereas 23E10-GAL4 dFB neurons are. Further work will need to determine the nature of the sleep-relevant information that is relayed by VNC-SP neurons.

## Materials and Methods

### Key Resources Table

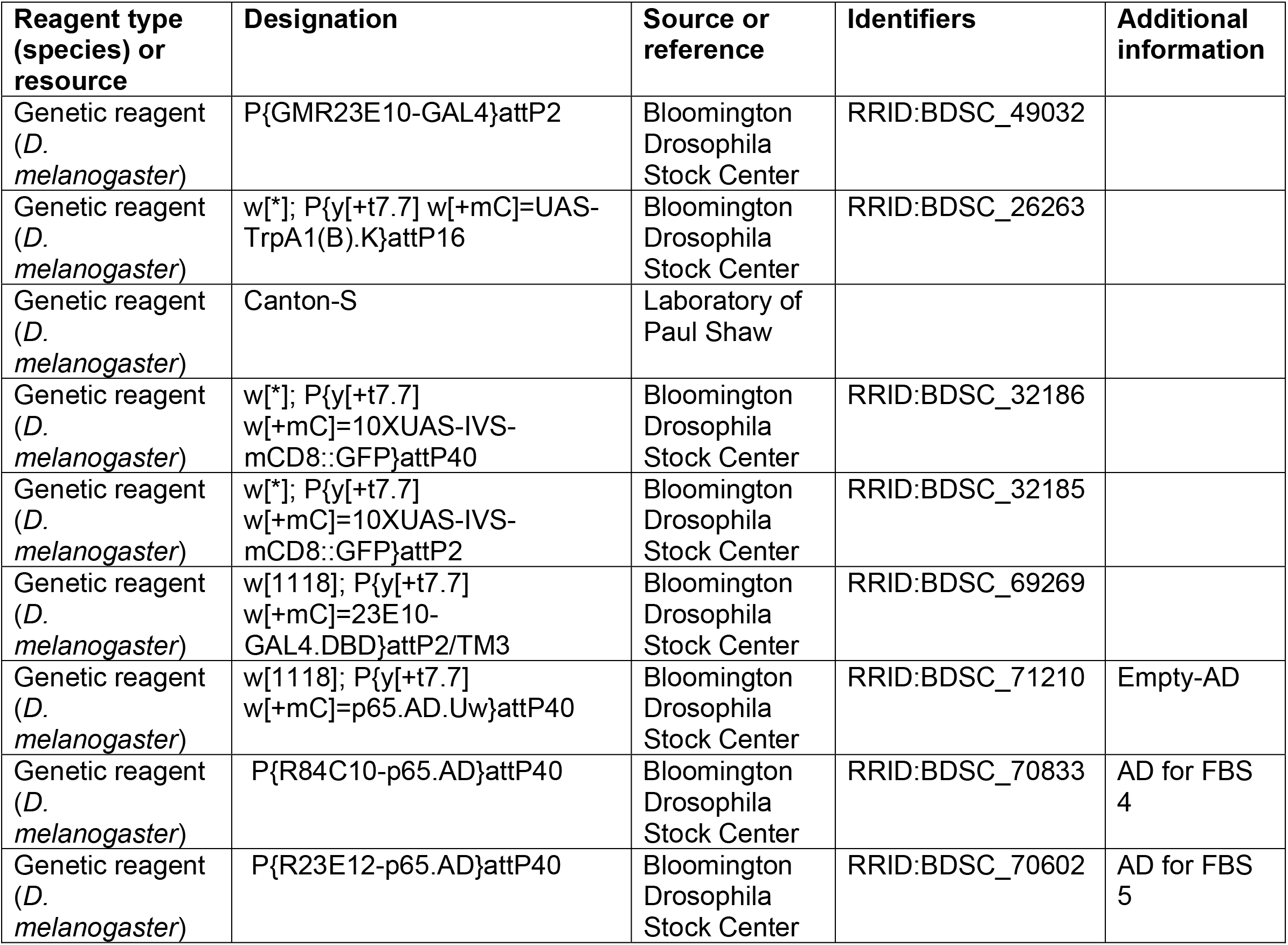
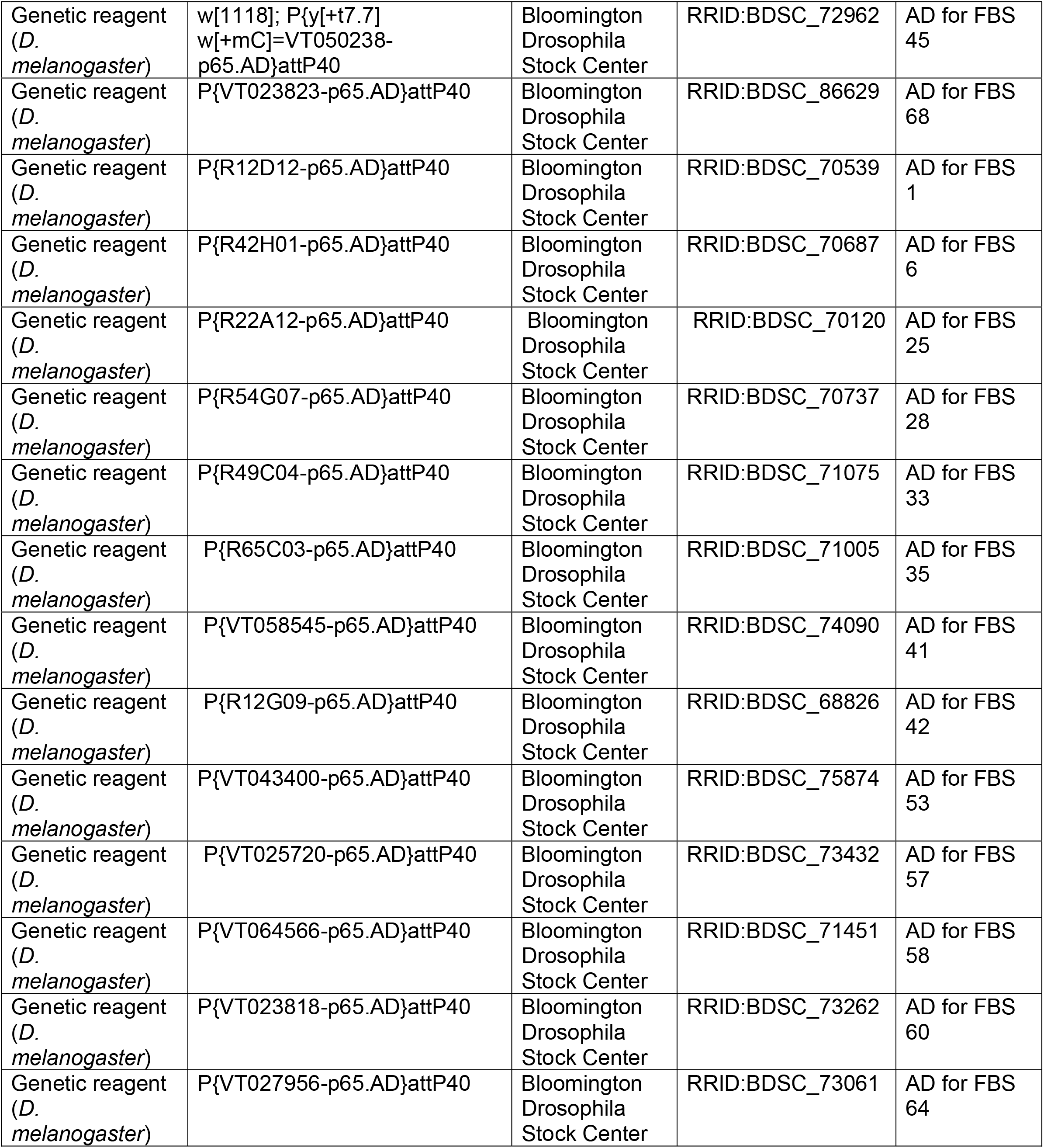
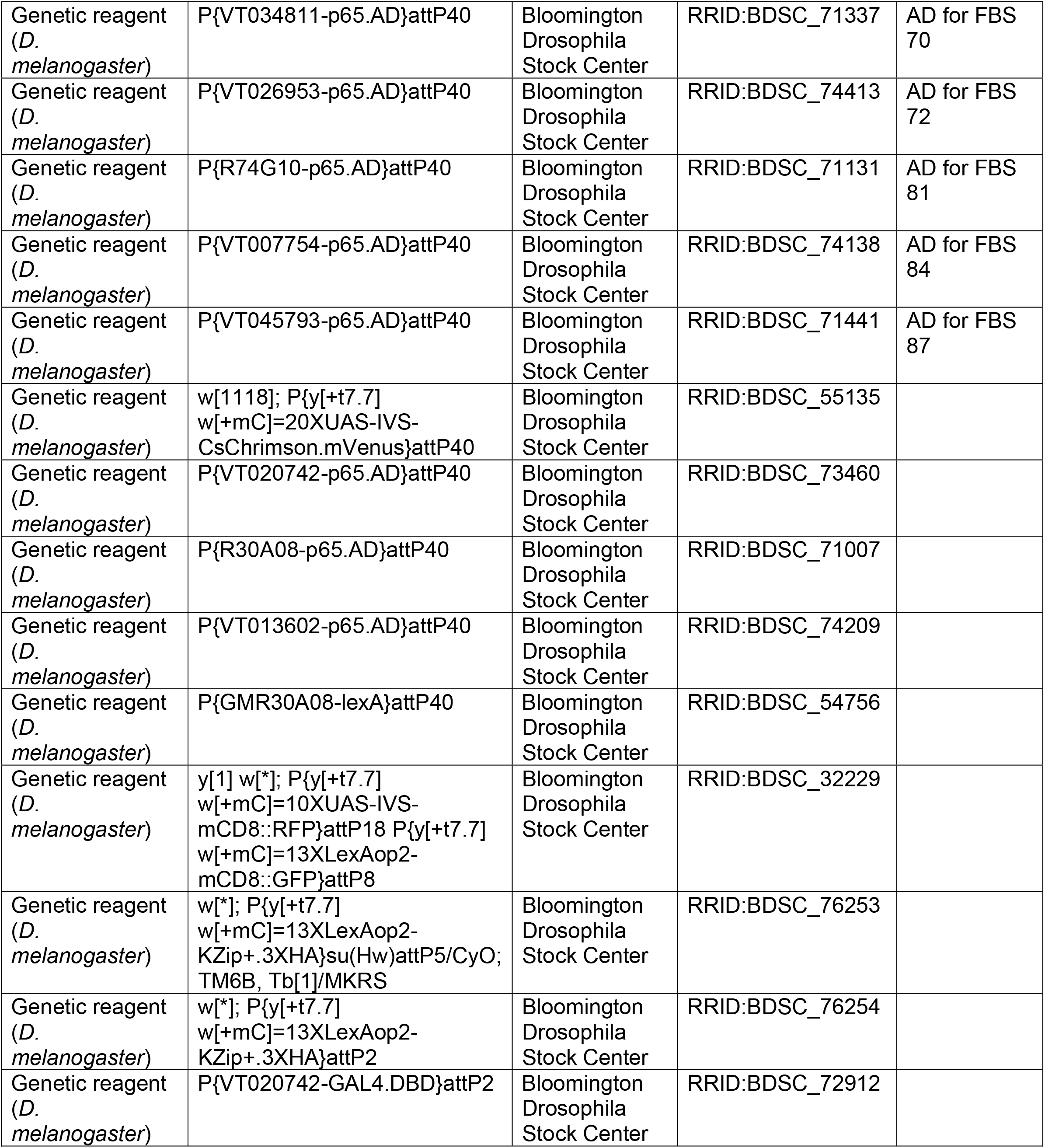
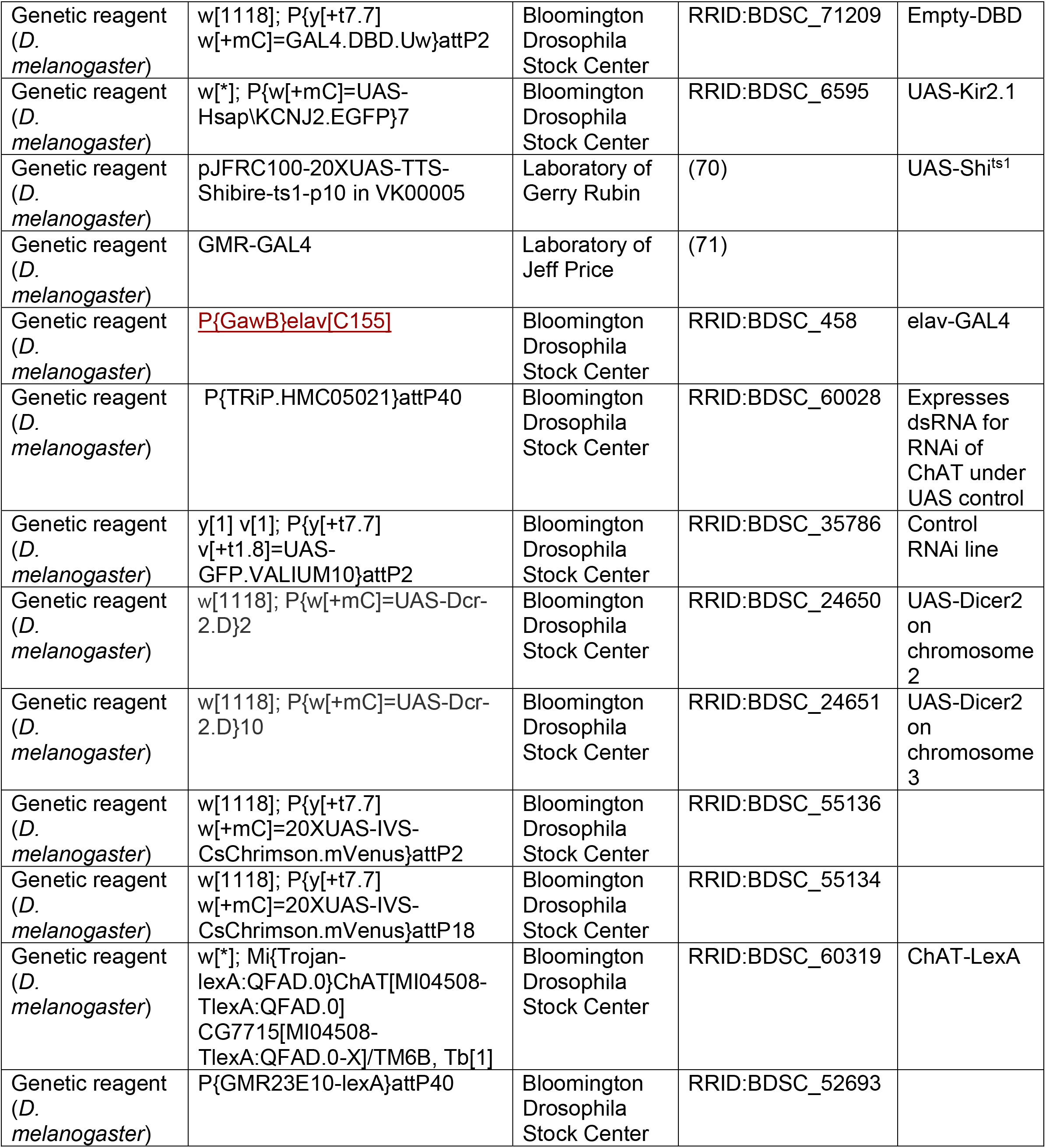
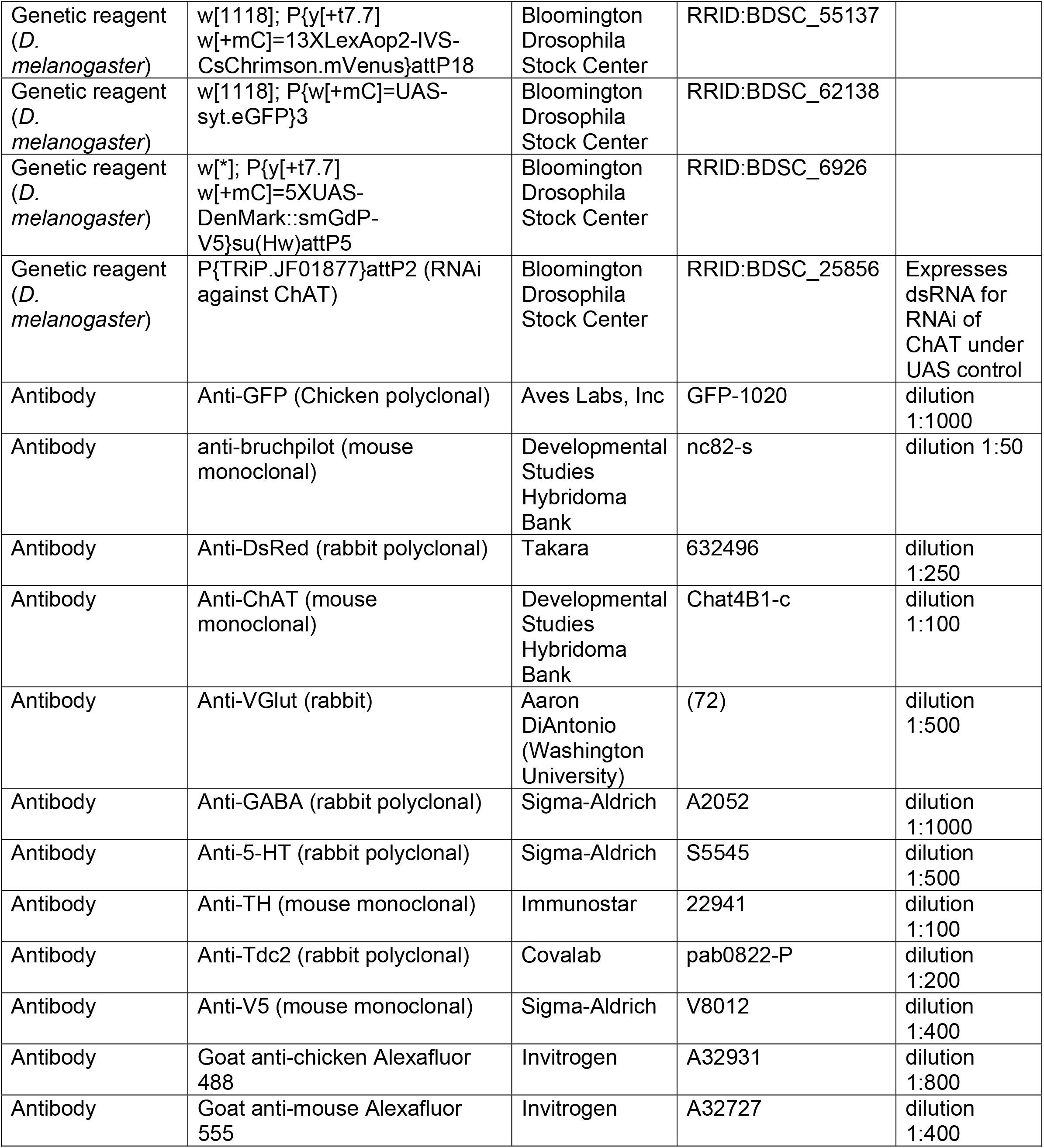
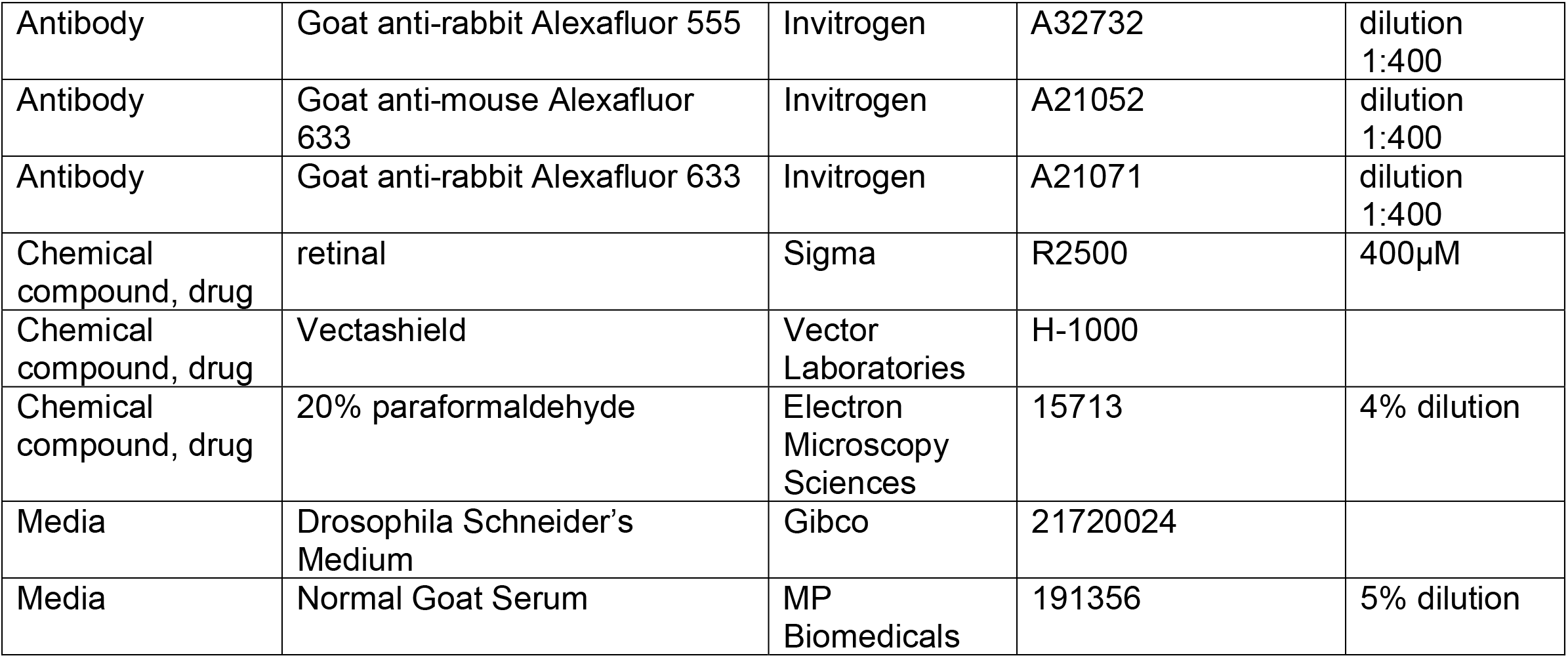

| Reagent type (species) or resource | Designation | Source or reference | Identifiers | Additional information |
| --- | --- | --- | --- | --- |
| Genetic reagent (*D. melanogaster*) | P{GMR23E10-GAL4}attP2 | Bloomington Drosophila Stock Center | RRID:BDSC_49032 |  |
| Genetic reagent (*D. melanogaster*) | w[*]; P{y[+t7.7] w[+mC]=UAS-TrpA1(B).K}attP16 | Bloomington Drosophila Stock Center | RRID:BDSC_26263 |  |
| Genetic reagent (*D. melanogaster*) | Canton-S | Laboratory of Paul Shaw |  |  |
| Genetic reagent (*D. melanogaster*) | w[*]; P{y[+t7.7] w[+mC]=10XUAS-IVS-mCD8::GFP}attP40 | Bloomington Drosophila Stock Center | RRID:BDSC_32186 |  |
| Genetic reagent (*D. melanogaster*) | w[*]; P{y[+t7.7] w[+mC]=10XUAS-IVS-mCD8::GFP}attP2 | Bloomington Drosophila Stock Center | RRID:BDSC_32185 |  |
| Genetic reagent (*D. melanogaster*) | w[1118]; P{y[+t7.7] w[+mC]=23E10-GAL4.DBD}attP2/TM3 | Bloomington Drosophila Stock Center | RRID:BDSC_69269 |  |
| Genetic reagent (*D. melanogaster*) | w[1118]; P{y[+t7.7] w[+mC]=p65.AD.Uw}attP40 | Bloomington Drosophila Stock Center | RRID:BDSC_71210 | Empty-AD |
| Genetic reagent (*D. melanogaster*) | P{R84C10-p65.AD}attP40 | Bloomington Drosophila Stock Center | RRID:BDSC_70833 | AD for FBS 4 |
| Genetic reagent (*D. melanogaster*) | P{R23E12-p65.AD}attP40 | Bloomington Drosophila Stock Center | RRID:BDSC_70602 | AD for FBS 5 |
| Genetic reagent<br>(*D. melanogaster*) | w[1118]; P{y[+t7.7]<br>w[+mC]=VT050238-<br>p65.AD}attP40 | Bloomington<br>Drosophila<br>Stock Center | RRID:BDSC_72962 | AD for FBS<br>45 |
| Genetic reagent<br>(*D. melanogaster*) | P{VT023823-p65.AD}attP40 | Bloomington<br>Drosophila<br>Stock Center | RRID:BDSC_86629 | AD for FBS<br>68 |
| Genetic reagent<br>(*D. melanogaster*) | P{R12D12-p65.AD}attP40 | Bloomington<br>Drosophila<br>Stock Center | RRID:BDSC_70539 | AD for FBS<br>1 |
| Genetic reagent<br>(*D. melanogaster*) | P{R42H01-p65.AD}attP40 | Bloomington<br>Drosophila<br>Stock Center | RRID:BDSC_70687 | AD for FBS<br>6 |
| Genetic reagent<br>(*D. melanogaster*) | P{R22A12-p65.AD}attP40 | Bloomington<br>Drosophila<br>Stock Center | RRID:BDSC_70120 | AD for FBS<br>25 |
| Genetic reagent<br>(*D. melanogaster*) | P{R54G07-p65.AD}attP40 | Bloomington<br>Drosophila<br>Stock Center | RRID:BDSC_70737 | AD for FBS<br>28 |
| Genetic reagent<br>(*D. melanogaster*) | P{R49C04-p65.AD}attP40 | Bloomington<br>Drosophila<br>Stock Center | RRID:BDSC_71075 | AD for FBS<br>33 |
| Genetic reagent<br>(*D. melanogaster*) | P{R65C03-p65.AD}attP40 | Bloomington<br>Drosophila<br>Stock Center | RRID:BDSC_71005 | AD for FBS<br>35 |
| Genetic reagent<br>(*D. melanogaster*) | P{VT058545-p65.AD}attP40 | Bloomington<br>Drosophila<br>Stock Center | RRID:BDSC_74090 | AD for FBS<br>41 |
| Genetic reagent<br>(*D. melanogaster*) | P{R12G09-p65.AD}attP40 | Bloomington<br>Drosophila<br>Stock Center | RRID:BDSC_68826 | AD for FBS<br>42 |
| Genetic reagent<br>(*D. melanogaster*) | P{VT043400-p65.AD}attP40 | Bloomington<br>Drosophila<br>Stock Center | RRID:BDSC_75874 | AD for FBS<br>53 |
| Genetic reagent<br>(*D. melanogaster*) | P{VT025720-p65.AD}attP40 | Bloomington<br>Drosophila<br>Stock Center | RRID:BDSC_73432 | AD for FBS<br>57 |
| Genetic reagent<br>(*D. melanogaster*) | P{VT064566-p65.AD}attP40 | Bloomington<br>Drosophila<br>Stock Center | RRID:BDSC_71451 | AD for FBS<br>58 |
| Genetic reagent<br>(*D. melanogaster*) | P{VT023818-p65.AD}attP40 | Bloomington<br>Drosophila<br>Stock Center | RRID:BDSC_73262 | AD for FBS<br>60 |
| Genetic reagent<br>(*D. melanogaster*) | P{VT027956-p65.AD}attP40 | Bloomington<br>Drosophila<br>Stock Center | RRID:BDSC_73061 | AD for FBS<br>64 |
| Genetic reagent<br>(*D. melanogaster*) | P{VT034811-p65.AD}attP40 | Bloomington<br>Drosophila<br>Stock Center | RRID:BDSC_71337 | AD for FBS<br>70 |
| Genetic reagent<br>(*D. melanogaster*) | P{VT026953-p65.AD}attP40 | Bloomington<br>Drosophila<br>Stock Center | RRID:BDSC_74413 | AD for FBS<br>72 |
| Genetic reagent<br>(*D. melanogaster*) | P{R74G10-p65.AD}attP40 | Bloomington<br>Drosophila<br>Stock Center | RRID:BDSC_71131 | AD for FBS<br>81 |
| Genetic reagent<br>(*D. melanogaster*) | P{VT007754-p65.AD}attP40 | Bloomington<br>Drosophila<br>Stock Center | RRID:BDSC_74138 | AD for FBS<br>84 |
| Genetic reagent<br>(*D. melanogaster*) | P{VT045793-p65.AD}attP40 | Bloomington<br>Drosophila<br>Stock Center | RRID:BDSC_71441 | AD for FBS<br>87 |
| Genetic reagent<br>(*D. melanogaster*) | w[1118]; P{y[+t7.7]<br>w[+mC]=20XUAS-IVS-<br>CsChrimson.mVenus}attP40 | Bloomington<br>Drosophila<br>Stock Center | RRID:BDSC_55135 |  |
| Genetic reagent<br>(*D. melanogaster*) | P{VT020742-p65.AD}attP40 | Bloomington<br>Drosophila<br>Stock Center | RRID:BDSC_73460 |  |
| Genetic reagent<br>(*D. melanogaster*) | P{R30A08-p65.AD}attP40 | Bloomington<br>Drosophila<br>Stock Center | RRID:BDSC_71007 |  |
| Genetic reagent<br>(*D. melanogaster*) | P{VT013602-p65.AD}attP40 | Bloomington<br>Drosophila<br>Stock Center | RRID:BDSC_74209 |  |
| Genetic reagent<br>(*D. melanogaster*) | P{GMR30A08-lexA}attP40 | Bloomington<br>Drosophila<br>Stock Center | RRID:BDSC_54756 |  |
| Genetic reagent<br>(*D. melanogaster*) | y[1] w[*]; P{y[+t7.7]<br>w[+mC]=10XUAS-IVS-<br>mCD8::RFP}attP18 P{y[+t7.7]<br>w[+mC]=13XLexAop2-<br>mCD8::GFP}attP8 | Bloomington<br>Drosophila<br>Stock Center | RRID:BDSC_32229 |  |
| Genetic reagent<br>(*D. melanogaster*) | w[*]; P{y[+t7.7]<br>w[+mC]=13XLexAop2-<br>KZip+.3XHA}su(Hw)attP5/CyO;<br>TM6B, Tb[1]/MKRS | Bloomington<br>Drosophila<br>Stock Center | RRID:BDSC_76253 |  |
| Genetic reagent<br>(*D. melanogaster*) | w[*]; P{y[+t7.7]<br>w[+mC]=13XLexAop2-<br>KZip+.3XHA}attP2 | Bloomington<br>Drosophila<br>Stock Center | RRID:BDSC_76254 |  |
| Genetic reagent<br>(*D. melanogaster*) | P{VT020742-GAL4.DBD}attP2 | Bloomington<br>Drosophila<br>Stock Center | RRID:BDSC_72912 |  |
| Genetic reagent<br>(*D. melanogaster*) | w[1118]; P{y[+t7.7]<br>w[+mC]=GAL4.DBD.Uw}attP2 | Bloomington<br>Drosophila<br>Stock Center | RRID:BDSC_71209 | Empty-DBD |
| Genetic reagent<br>(*D. melanogaster*) | w[*]; P{w[+mC]=UAS-<br>Hsap\KCNJ2.EGFP}7 | Bloomington<br>Drosophila<br>Stock Center | RRID:BDSC_6595 | UAS-Kir2.1 |
| Genetic reagent<br>(*D. melanogaster*) | pJFRC100-20XUAS-TTS-Shibire-ts1-p10 in VK00005 | Laboratory of<br>Gerry Rubin | (70) | UAS-Shi<sup>ts1</sup> |
| Genetic reagent<br>(*D. melanogaster*) | GMR-GAL4 | Laboratory of<br>Jeff Price | (71) |  |
| Genetic reagent<br>(*D. melanogaster*) | P{GawB}elav[C155] | Bloomington<br>Drosophila<br>Stock Center | RRID:BDSC_458 | elav-GAL4 |
| Genetic reagent<br>(*D. melanogaster*) | P{TRiP.HMC05021}attP40 | Bloomington<br>Drosophila<br>Stock Center | RRID:BDSC_60028 | Expresses<br>dsRNA for<br>RNAi of<br>ChAT under<br>UAS control |
| Genetic reagent<br>(*D. melanogaster*) | y[1] v[1]; P{y[+t7.7]<br>v[+t1.8]=UAS-<br>GFP.VALIUM10}attP2 | Bloomington<br>Drosophila<br>Stock Center | RRID:BDSC_35786 | Control<br>RNAi line |
| Genetic reagent<br>(*D. melanogaster*) | w[1118]; P{w[+mC]=UAS-Dcr-<br>2.D}2 | Bloomington<br>Drosophila<br>Stock Center | RRID:BDSC_24650 | UAS-Dicer2<br>on<br>chromosome<br>2 |
| Genetic reagent<br>(*D. melanogaster*) | w[1118]; P{w[+mC]=UAS-Dcr-<br>2.D}10 | Bloomington<br>Drosophila<br>Stock Center | RRID:BDSC_24651 | UAS-Dicer2<br>on<br>chromosome<br>3 |
| Genetic reagent<br>(*D. melanogaster*) | w[1118]; P{y[+t7.7]<br>w[+mC]=20XUAS-IVS-<br>CsChrimson.mVenus}attP2 | Bloomington<br>Drosophila<br>Stock Center | RRID:BDSC_55136 |  |
| Genetic reagent<br>(*D. melanogaster*) | w[1118]; P{y[+t7.7]<br>w[+mC]=20XUAS-IVS-<br>CsChrimson.mVenus}attP18 | Bloomington<br>Drosophila<br>Stock Center | RRID:BDSC_55134 |  |
| Genetic reagent<br>(*D. melanogaster*) | w[*]; Mi{Trojan-<br>lexA:QFAD.0}ChAT[MI04508-TlexA:QFAD.0]<br>CG7715[MI04508-TlexA:QFAD.0-X]/TM6B, Tb[1] | Bloomington<br>Drosophila<br>Stock Center | RRID:BDSC_60319 | ChAT-LexA |
| Genetic reagent<br>(*D. melanogaster*) | P{GMR23E10-lexA}attP40 | Bloomington<br>Drosophila<br>Stock Center | RRID:BDSC_52693 |  |
| Genetic reagent<br>(*D. melanogaster*) | w[1118]; P{y[+t7.7]<br>w[+mC]=13XLexAop2-IVS-<br>CsChrimson.mVenus}attP18 | Bloomington<br>Drosophila<br>Stock Center | RRID:BDSC_55137 |  |
| Genetic reagent<br>(*D. melanogaster*) | w[1118]; P{w[+mC]=UAS-<br>syt.eGFP}3 | Bloomington<br>Drosophila<br>Stock Center | RRID:BDSC_62138 |  |
| Genetic reagent<br>(*D. melanogaster*) | w[*]; P{y[+t7.7]<br>w[+mC]=5XUAS-<br>DenMark::smGdP-V5}su(Hw)attP5 | Bloomington<br>Drosophila<br>Stock Center | RRID:BDSC_6926 |  |
| Genetic reagent<br>(*D. melanogaster*) | P{TRiP.JF01877}attP2 (RNAi<br>against ChAT) | Bloomington<br>Drosophila<br>Stock Center | RRID:BDSC_25856 | Expresses<br>dsRNA for<br>RNAi of<br>ChAT under<br>UAS control |
| Antibody | Anti-GFP (Chicken polyclonal) | Aves Labs, Inc | GFP-1020 | dilution<br>1:1000 |
| Antibody | anti-bruchpilot (mouse<br>monoclonal) | Developmental<br>Studies<br>Hybridoma<br>Bank | nc82-s | dilution 1:50 |
| Antibody | Anti-DsRed (rabbit polyclonal) | Takara | 632496 | dilution<br>1:250 |
| Antibody | Anti-ChAT (mouse<br>monoclonal) | Developmental<br>Studies<br>Hybridoma<br>Bank | Chat4B1-c | dilution<br>1:100 |
| Antibody | Anti-VGluT (rabbit) | Aaron<br>DiAntonio<br>(Washington<br>University) | (72) | dilution<br>1:500 |
| Antibody | Anti-GABA (rabbit polyclonal) | Sigma-Aldrich | A2052 | dilution<br>1:1000 |
| Antibody | Anti-5-HT (rabbit polyclonal) | Sigma-Aldrich | S5545 | dilution<br>1:500 |
| Antibody | Anti-TH (mouse monoclonal) | Immunostar | 22941 | dilution<br>1:100 |
| Antibody | Anti-Tdc2 (rabbit polyclonal) | Covalab | pab0822-P | dilution<br>1:200 |
| Antibody | Anti-V5 (mouse monoclonal) | Sigma-Aldrich | V8012 | dilution<br>1:400 |
| Antibody | Goat anti-chicken Alexafluor<br>488 | Invitrogen | A32931 | dilution<br>1:800 |
| Antibody | Goat anti-mouse Alexafluor<br>555 | Invitrogen | A32727 | dilution<br>1:400 |
| Antibody | Goat anti-rabbit Alexafluor 555 | Invitrogen | A32732 | dilution 1:400 |
| Antibody | Goat anti-mouse Alexafluor 633 | Invitrogen | A21052 | dilution 1:400 |
| Antibody | Goat anti-rabbit Alexafluor 633 | Invitrogen | A21071 | dilution 1:400 |
| Chemical compound, drug | retinal | Sigma | R2500 | 400µM |
| Chemical compound, drug | Vectashield | Vector Laboratories | H-1000 |  |
| Chemical compound, drug | 20% paraformaldehyde | Electron Microscopy Sciences | 15713 | 4% dilution |
| Media | Drosophila Schneider's Medium | Gibco | 21720024 |  |
| Media | Normal Goat Serum | MP Biomedicals | 191356 | 5% dilution |

### Flies

Flies were cultured at 25°C with 50% humidity under a 12-hour light: 12-hour dark cycle. Flies were kept on a standard yeast and molasses diet (per 1L: 50g yeast, 15g sucrose, 28g corn syrup, 33.3mL molasses, 9g agar).

### Sleep

Sleep was assessed as previously described (32, 73). 4-10 day old virgin females or males were used as described in figure legends. Briefly, flies were placed into individual 65 mm tubes and all activity was continuously measured through the Trikinetics Drosophila Activity Monitoring System (www.Trikinetics.com, Waltham, Ma). Locomotor activity was measured in 1-minute bins and sleep was defined as periods of quiescence lasting at least 5 minutes.

### Optogenetic activation

For optogenetic activation, flies expressing CsChrimson were loaded into individual 65mm tubes with food supplemented with 400µM all-trans retinal (Sigma, #R2500) and kept in the dark for 48 hours. Sleep was then monitored under low-light conditions for 48 hours of baseline measurements. To activate CSChrimson, flies were put under 627nm LEDs (LuxeonStar LXM2-PD01-0040) set to a pulse cycle of (5ms on, 95ms off) x 20 with a 4 second delay between pulse cycles. The light intensity of this protocol is 0.024mW/mm^2^

### Sleep deprivation

Sleep deprivation was performed as previously described (32, 74). Briefly, flies were placed into individual 65 mm tubes and the sleep-nullifying apparatus (SNAP) was used to sleep deprive flies for 12 hours during the dark phase. Sleep homeostasis was calculated for each individual fly as the ratio of the minutes of sleep gained above baseline during 24h of recovery sleep divided by the total min of sleep lost during 12 h of sleep deprivation.

### Arousal threshold

Retinal or vehicle-fed VNC-SP>UAS-CsChrimson flies were placed into individual 65 mm tubes in Trikinetics monitors. These were then loaded in the SNAP sleep deprivation apparatus. The SNAP tilts asymmetrically from −60° to +60°, such that sleeping flies are disturbed during each downward movement (75). Each individual SNAP movement takes 15 seconds to complete. To assess arousal threshold, we modified the SNAP parameters so that flies were subjected to only 1, 2, or 4 downward movements in the SNAP every hour for 24h and their locomotor responses were analyzed. We then averaged each individual hour response to obtain the percentage of flies awakened by a given strength of perturbation.

### QPCR

QPCR were performed as previously described (32, 76). Briefly, total RNA was isolated from the bodies and eyes of ∼10 flies per replicate with Trizol (Invitrogen, Carlsbad, CA) and DNAse I digested. cDNA synthesis was performed using Superscript IV VILO (Invitrogen) according to manufacturer protocol. cDNA samples were loaded in duplicate and quantitatively analyzed using SYBR Green Master Mix (Applied Biosystems) and gene specific primers using the Applied Biosystems 7500 system. Expression values for RP49 were used to normalize results between groups. The following primer sets were used: rp49: fw aagaagcgcaccaagcacttcatc, rev tctgttgtcgatacccttgggctt and ChAT: fw caccgagcgatacaggatgg, rev ggcaccttgggtagagtgtc.

### Immunocytochemistry

Flies were cold anesthetized, brains and VNC were dissected in ice cold Schneider’s Drosophila Medium (Gibco, 21720024) and blocked in 5% normal goat serum. Following blocking flies were incubated overnight at 4°C in primary antibody, washed in PBST and incubated overnight at 4°C in secondary antibody. Description of antibodies used can be found in the resources table. Brains and VNCs were mounted on polylysine treated slides in Vectashield H-1000 mounting medium. Imaging was performed on a Zeiss 510 meta confocal microscope using a Plan-Apochromat 20x or Plan-Neofluar 40x objective. Z-series images were acquired with a 1 uM slice size using the same settings (laser power, gain, offset) to allow for comparison across genotypes. Images were processed and analyzed using ImageJ.

## Statistics

Statistical analyses were performed with Prism9 software (GraphPad). Normal distribution was assessed with the D’Agostino-Pearson test. Normally distributed data were analyzed with parametric statistics: t-test, one-way analysis of variance or two-way ANOVA followed by the planned pairwise multiple comparisons as described in the legends. For data that significantly differed from the normal distribution, non-parametric statistics were applied, Mann-Whitney U test or Kruskal–Wallis test followed by Dunn’s multiple test. Some non-normally distributed data were subjected to log transformation or Box-Cox transformation before two-way ANOVA followed by planned pairwise multiple comparisons as described in the legends. All statistically different groups are defined as *P < 0.05, **P<0.01, ***P<0.001 and ****P<0.0001.

## Author Contributions

S.D., J.J. and J.M.E. designed the experiments and wrote the paper. J.J., B.H., K.E., A.V., A.V., A.E., J.M.E and S.D. performed the experiments. J.J., J.M.E and S.D. analyzed the data.

## Acknowledgments

We thank Aaron DiAntonio, Gerry Rubin, Paul Shaw and Jeff Price for sharing reagents. We also thank Krishna Melnattur for comments on this manuscript.

## Video Legends

**Video 1: Brain of a 23E10-GAL4>UAS-GFP female.**

**Video 2: VNC of a 23E10-GAL4>UAS-GFP female.**

**Video 3: Brain of a FBS5>UAS-GFP female.**

**Video 4: VNC of a FBS5>UAS-GFP female.**

**Video 5: Brain of a VT013602AD-23E10-DBD (VNC-SP)>UAS-GFP female.**

**Video 6: VNC of a VT013602AD-23E10-DBD (VNC-SP)>UAS-GFP female.**

## Supplemental Figure Legends

**Figure 2-figure supplement 1:**
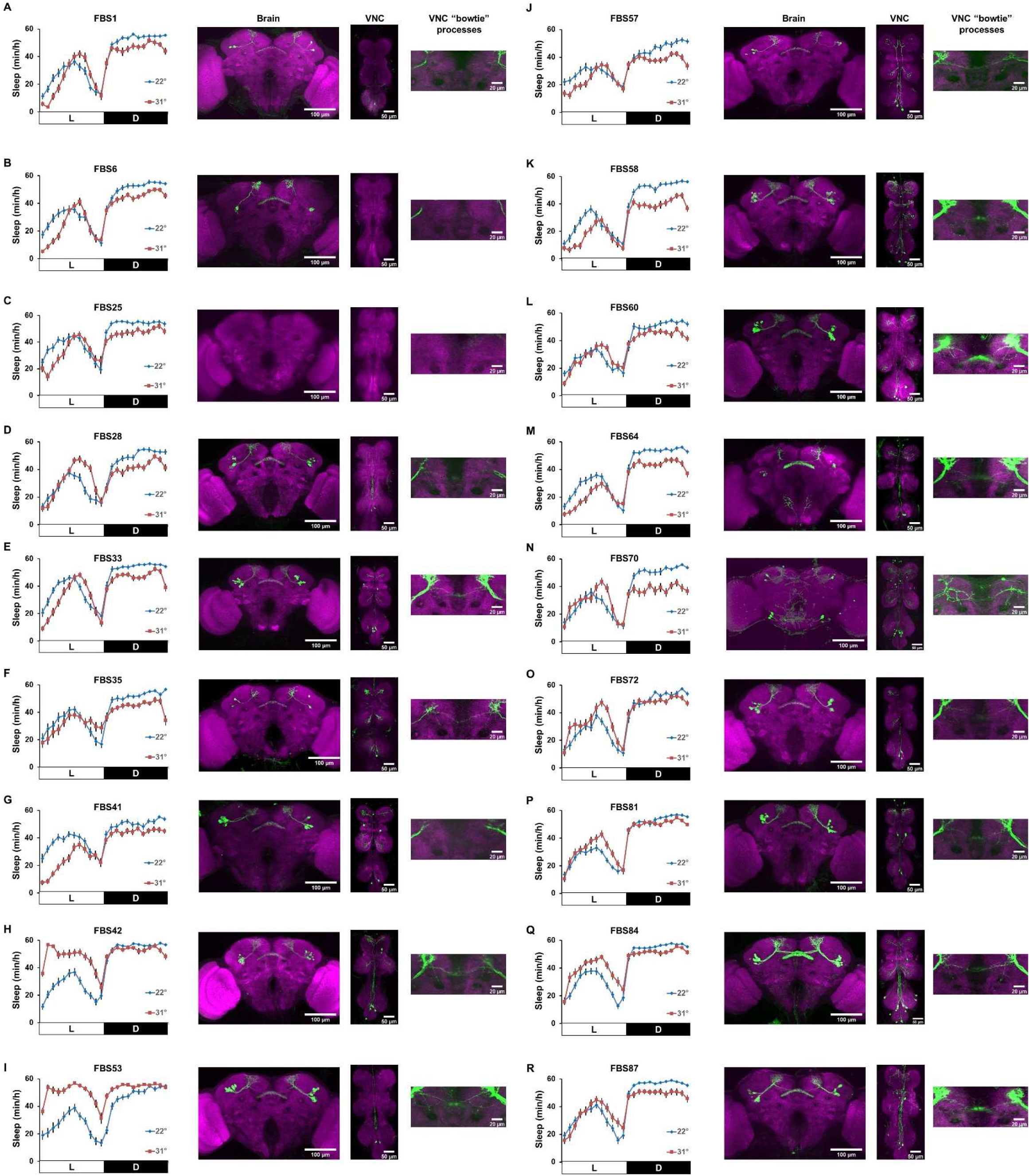
Additional sleep profiles and confocal images of FBS lines. **A-R)** Sleep profile (left) and representative confocal stacks (right) for brain and VNC of female *FBS>UAS-TrpA1; UAS-GFP* for each FBS line not presented in Figure 2.

**Figure 2-figure supplement 2:**
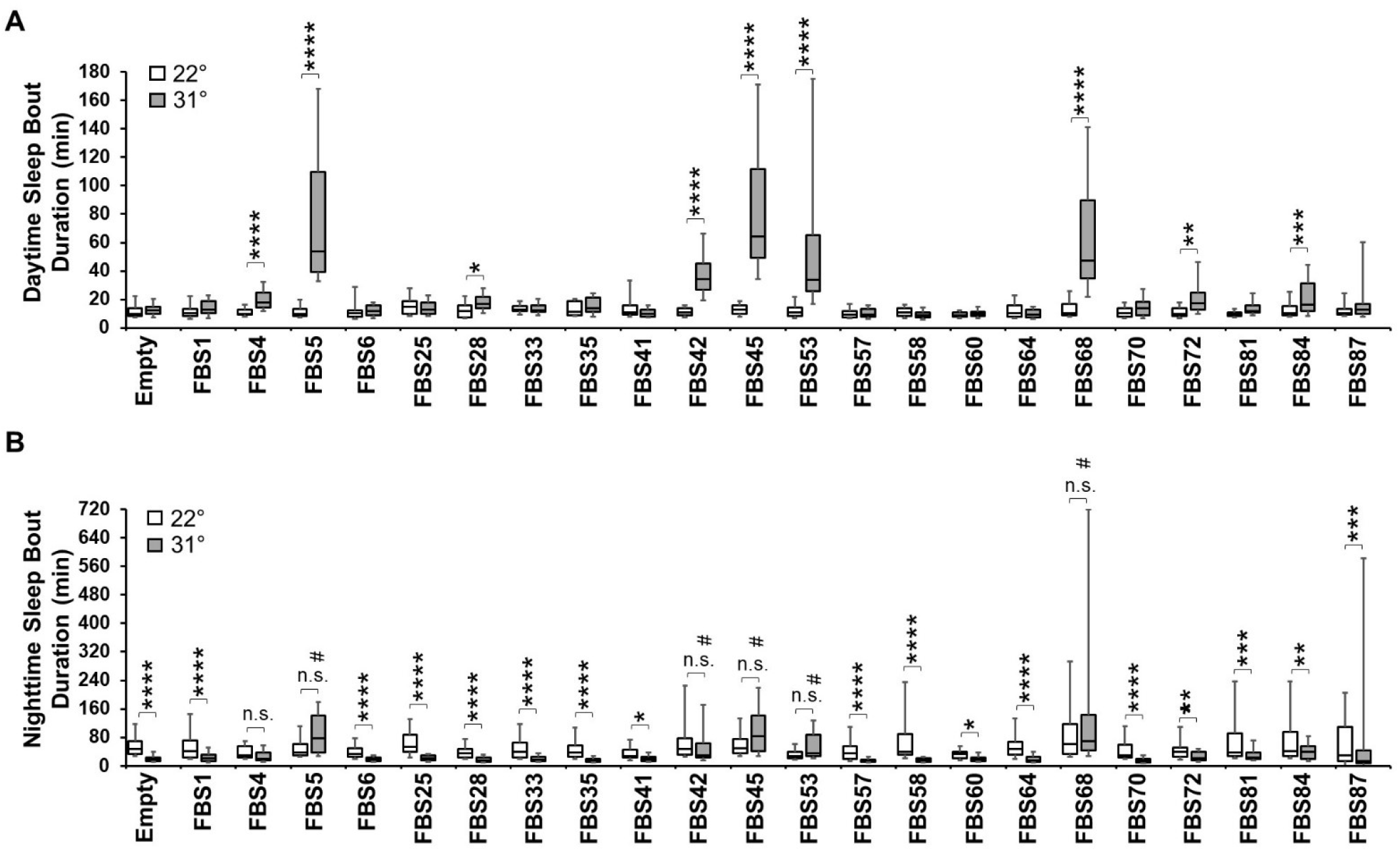
Additional sleep parameters for FBS female flies thermogenetically activated. **A)** Box plots of daytime sleep bout duration in minutes for flies presented in Figure 2B. The bottom and top of each box represents the first and third quartile, and the horizontal line dividing the box is the median. The whiskers represent the 10^th^ and 90^th^ percentiles. Two-way repeated measures ANOVA followed by Sidak’s multiple comparisons test revealed that 9 FBS line show a significant increase in daytime sleep bout duration between 22°C and 31°C. *P<0.05, **P<0.01, ***P<0.001, ****P<0.0001, n=30-51 flies per genotype. **B)** Box plots of nighttime sleep bout duration in minutes for flies presented in Figure 2B. The bottom and top of each box represents the first and third quartile, and the horizontal line dividing the box is the median. The whiskers represent the 10^th^ and 90^th^ percentiles. Two-way repeated measures ANOVA followed by Sidak’s multiple comparisons test revealed that control and most FBS line show a significant decrease in nighttime sleep bout duration between 22°C and 31°C. Only 6 sleep-promoting FBS lines show no difference between 22°C and 31°C. Dunnett’s multiple comparisons reveal that for 5 of them, nighttime sleep bout duration at 31°C is significantly increased compared with Empty control flies (# on figure). *P<0.05, **P<0.01, ***P<0.001, ****P<0.0001, n.s= not significant. n=30-51 flies per genotype.

**Figure 2-figure supplement 3:**
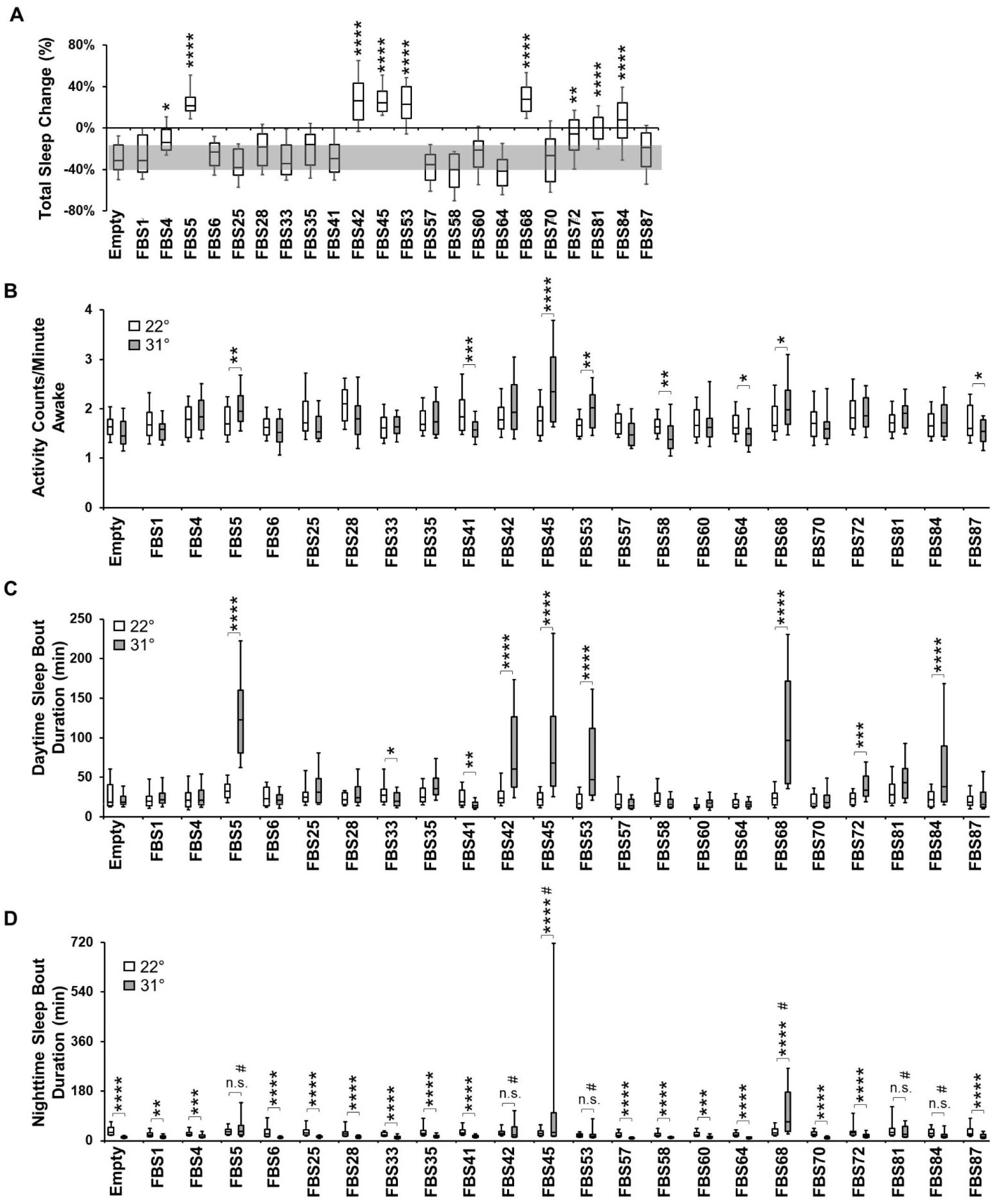
Thermogenetic activation in male flies. **A)** Box plots of total sleep change in % ((total sleep on day3-total sleep on day2/total sleep on day2) X 100) for male control (Empty-AD; 23E10-DBD) and 22 FBS lines expressing UAS-TrpA1; UAS-GFP. The bottom and top of each box represents the first and third quartile, and the horizontal line dividing the box is the median. The whiskers represent the 10^th^ and 90^th^ percentiles. The gray rectangle spanning the horizontal axis indicates the interquartile range of the control. Kruskal-Wallis ANOVA followed by Dunn’s multiple comparisons revealed that 9 FBS lines increase sleep significantly more than control flies when transferred to 31°C. *P<0.05, **P<0.01, ****P<0.0001, n=26-46 flies per genotype. **B)** Box plots of locomotor activity counts per minute awake for flies presented in A. The bottom and top of each box represents the first and third quartile, and the horizontal line dividing the box is the median. The whiskers represent the 10^th^ and 90^th^ percentiles. Two-way repeated measures ANOVA followed by Sidak’s multiple comparisons test found that for 4 sleep-promoting FBS lines (FBS5, FBS45, FBS53 and FBS68) locomotor activity per awake time is increased while no differences are seen for the other 5 sleep-promoting lines between 22°C and 31°C. *P<0.05, **P<0.01, ***P<0.001, ****P<0.0001, n=26-46 flies per genotype. **C)** Box plots of daytime sleep bout duration for flies presented in A. The bottom and top of each box represents the first and third quartile, and the horizontal line dividing the box is the median. The whiskers represent the 10^th^ and 90^th^ percentiles. Two-way repeated measures ANOVA followed by Sidak’s multiple comparisons test found that for 7 out of the 9 sleep-promoting FBS lines, daytime sleep bout duration is significantly increased at 31°C compared with 22°C. *P<0.05, **P<0.01, ***P<0.001, ****P<0.0001, n=26-46 flies per genotype. **D)** Box plots of nighttime sleep bout duration for flies presented in A. The bottom and top of each box represents the first and third quartile, and the horizontal line dividing the box is the median. The whiskers represent the 10^th^ and 90^th^ percentiles. Two-way repeated measures ANOVA followed by Sidak’s multiple comparisons revealed that control and most FBS line show a significant decrease in nighttime sleep bout duration between 22°C and 31°C. 5 sleep-promoting FBS lines show no difference between 22°C and 31°C (FBS5, FBS42, FBS53, FBS81 and FBS84) while FBS45 and FBS68 show an increase in nighttime sleep bout duration at 31°C. Dunnett’s multiple comparisons reveal that for FBS5, FBS42, FBS45, FBS53, FBS68,FBS81 and FBS84, nighttime sleep bout duration at 31°C is significantly increased compared with Empty control flies (# on figure). **P<0.01, ***P<0.001, ****P<0.0001, n=26-46 flies per genotype.

**Figure 3-figure supplement 1:**
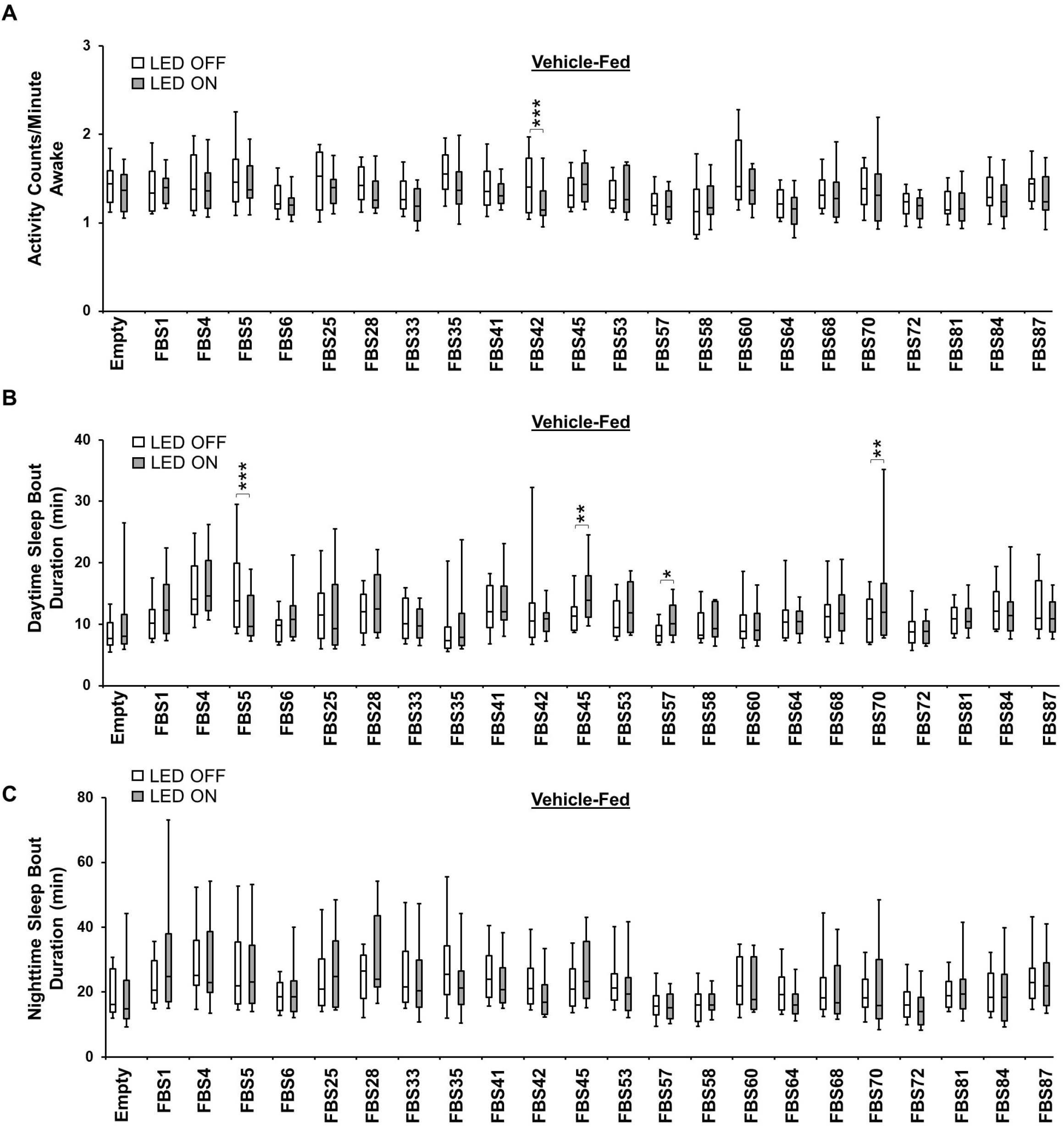
Vehicle-fed sleep data for females in optogenetic experiments. **A)** Box plots of locomotor activity counts per minute awake for vehicle-fed flies presented in Figure 3B. The bottom and top of each box represents the first and third quartile, and the horizontal line dividing the box is the median. The whiskers represent the 10^th^ and 90^th^ percentiles. Two-way repeated measures ANOVA followed by Sidak’s multiple comparisons test found that all lines except FBS42 show no difference in locomotor activity per awake time when the flies are stimulated with 627nm LEDs. ***P<0.001. n=20-39 flies per genotype. **B)** Box plots of daytime sleep bout duration for vehicle-fed flies presented in Figure 3B. The bottom and top of each box represents the first and third quartile, and the horizontal line dividing the box is the median. The whiskers represent the 10^th^ and 90^th^ percentiles. Two-way repeated measures ANOVA followed by Sidak’s multiple comparisons test found that most vehicle-fed sleep-promoting lines show no difference in daytime sleep bout duration when the flies are stimulated with 627nm LEDs. *P<0.05, **P<0.01, ***P<0.001. n=20-39 flies per genotype. **C)** Box plots of nighttime sleep bout duration for vehicle-fed flies presented in Figure 3B. The bottom and top of each box represents the first and third quartile, and the horizontal line dividing the box is the median. The whiskers represent the 10^th^ and 90^th^ percentiles. Two-way repeated measures ANOVA followed by Sidak’s multiple comparisons test show no difference in nighttime sleep bout duration when vehicle-fed flies are stimulated with 627nm LEDs. n=20-39 flies per genotype.

**Figure 3-figure supplement 2:**
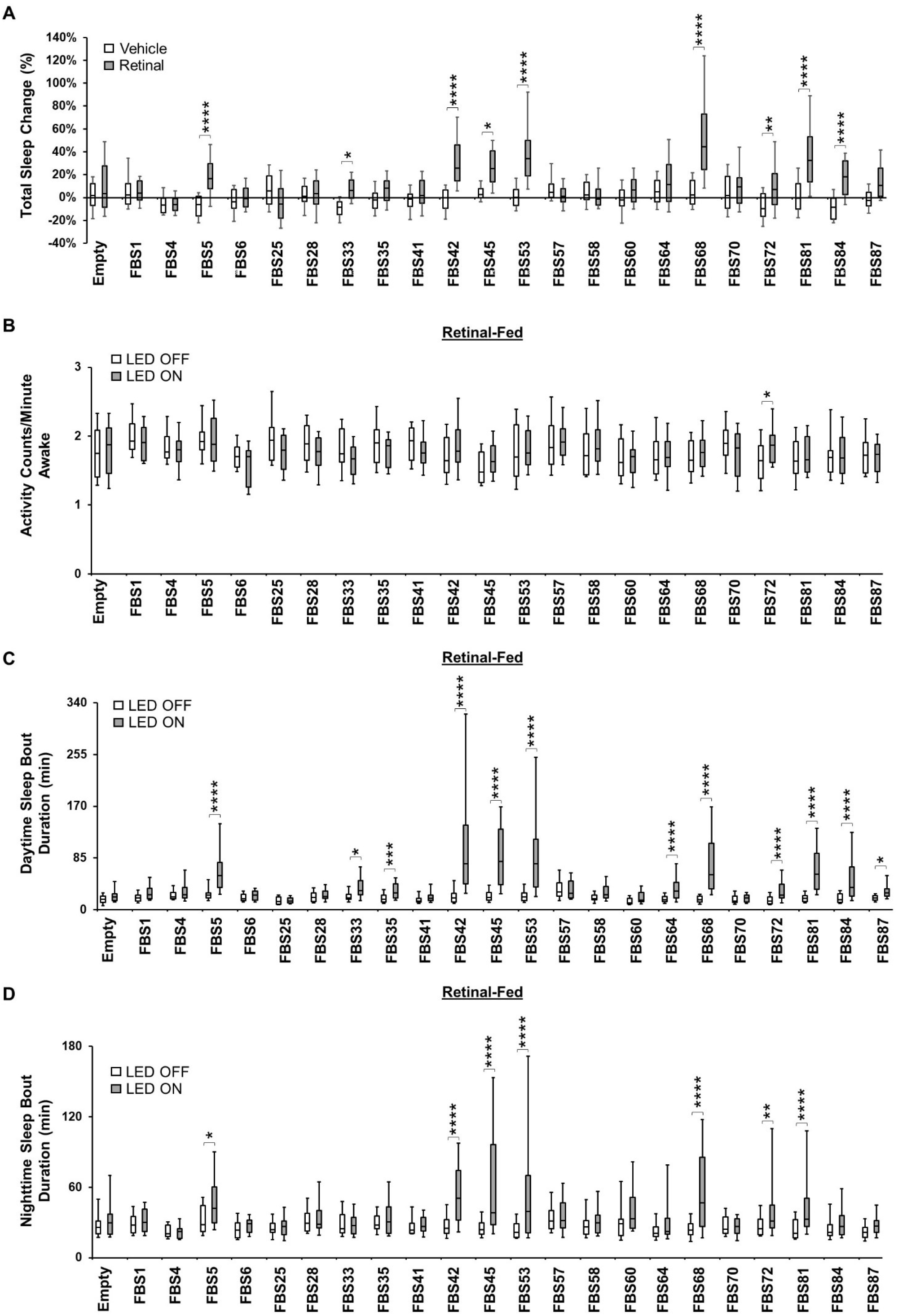
Optogenetic activation in males. **A)** Box plots of total sleep change in % ((total sleep on day3-total sleep on day2/total sleep on day2) X 100) for control (Empty) and 22 experimental vehicle-fed and retinal-fed male flies expressing CsChrimson upon 627nm LED stimulation. The bottom and top of each box represents the first and third quartile, and the horizontal line dividing the box is the median. The whiskers represent the 10^th^ and 90^th^ percentiles. Two-way ANOVA followed by Sidak’s multiple comparisons revealed that 9 retinal-fed FBS lines increase sleep significantly when stimulated with 627nm LEDs when compared with vehicle-fed flies. *P<0.05, **P<0.01, ****P<0.0001. n=20-44 flies per genotype and condition. **B)** Box plots of locomotor activity counts per minute awake for retinal-fed flies presented in A. The bottom and top of each box represents the first and third quartile, and the horizontal line dividing the box is the median. The whiskers represent the 10^th^ and 90^th^ percentiles. Two-way repeated measures ANOVA followed by Sidak’s multiple comparisons test found that for most sleep-promoting FBS lines, locomotor activity per awake time is not affected when the flies are stimulated with 627nm LEDs while it is increased in *FBS72>UAS-CsChrimson* flies. *P<0.05, n=24-44 flies per genotype. **C)** Box plots of daytime sleep bout duration (in minutes) for retinal-fed flies presented in A. The bottom and top of each box represents the first and third quartile, and the horizontal line dividing the box is the median. The whiskers represent the 10^th^ and 90^th^ percentiles. Two-way repeated measures ANOVA followed by Sidak’s multiple comparisons indicate that daytime sleep bout duration is increased in 12 FBS lines expressing CsChrimson when stimulated with 627nm LEDs. *P<0.05, ***P<0.001, ****P<0.0001, n=24-44 flies per genotype. **D)** Box plots of nighttime sleep bout duration (in minutes) for retinal-fed flies presented in A. The bottom and top of each box represents the first and third quartile, and the horizontal line dividing the box is the median. The whiskers represent the 10^th^ and 90^th^ percentiles. Two-way repeated measures ANOVA followed by Sidak’s multiple comparisons indicate that nighttime sleep bout duration is increased in 7 FBS lines expressing CsChrimson when stimulated with 627nm LEDs. *P<0.05, **P<0.01, ****P<0.0001, n=24-44 flies per genotype.

**Figure 3-figure supplement 3:**
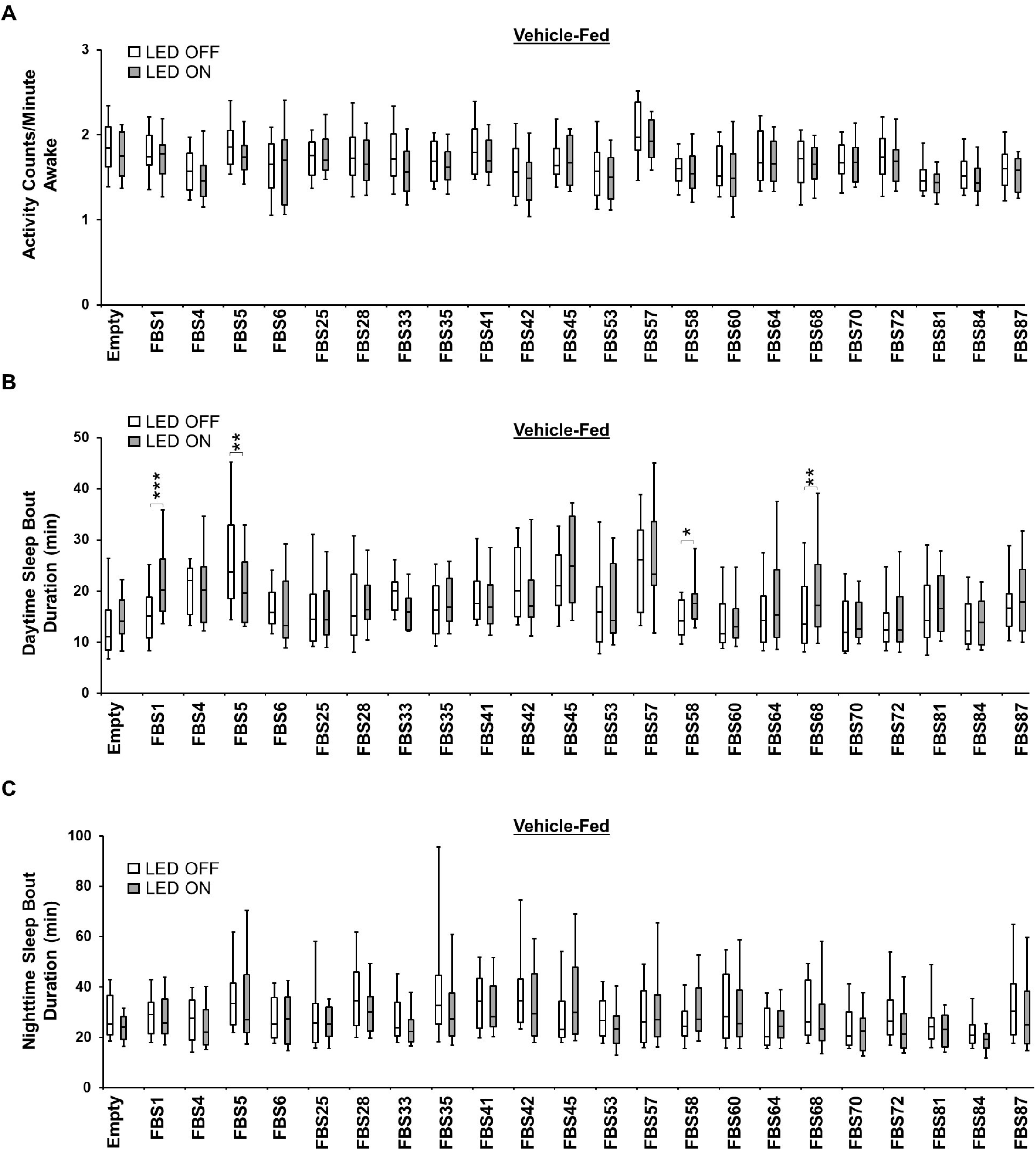
Vehicle-fed sleep data for males in optogenetic experiments. **A)** Box plots of locomotor activity counts per minute awake for vehicle-fed flies presented in Figure 3-figure supplement 2A. The bottom and top of each box represents the first and third quartile, and the horizontal line dividing the box is the median. The whiskers represent the 10^th^ and 90^th^ percentiles. Two-way repeated measures ANOVA followed by Sidak’s multiple comparisons test found no difference in locomotor activity per awake time when the flies are stimulated with 627nm LEDs. n=20-40 flies per genotype. **B)** Box plots of daytime sleep bout duration for vehicle-fed flies presented in Figure 3-figure supplement 2A. The bottom and top of each box represents the first and third quartile, and the horizontal line dividing the box is the median. The whiskers represent the 10^th^ and 90^th^ percentiles. Two-way repeated measures ANOVA followed by Sidak’s multiple comparisons test found that most vehicle-fed sleep-promoting lines show no difference in daytime sleep bout duration when the flies are stimulated with 627nm LEDs. *P<0.05, **P<0.01, ***P<0.001. n=20-40 flies per genotype. **C)** Box plots of nighttime sleep bout duration for vehicle-fed flies presented in Figure 3-figure supplement 2A. The bottom and top of each box represents the first and third quartile, and the horizontal line dividing the box is the median. The whiskers represent the 10^th^ and 90^th^ percentiles. Two-way repeated measures ANOVA followed by Sidak’s multiple comparisons test show no difference in nighttime sleep bout duration when vehicle-fed flies are stimulated with 627nm LEDs. n=20-40 flies per genotype.

**Figure 4-figure supplement 1:**
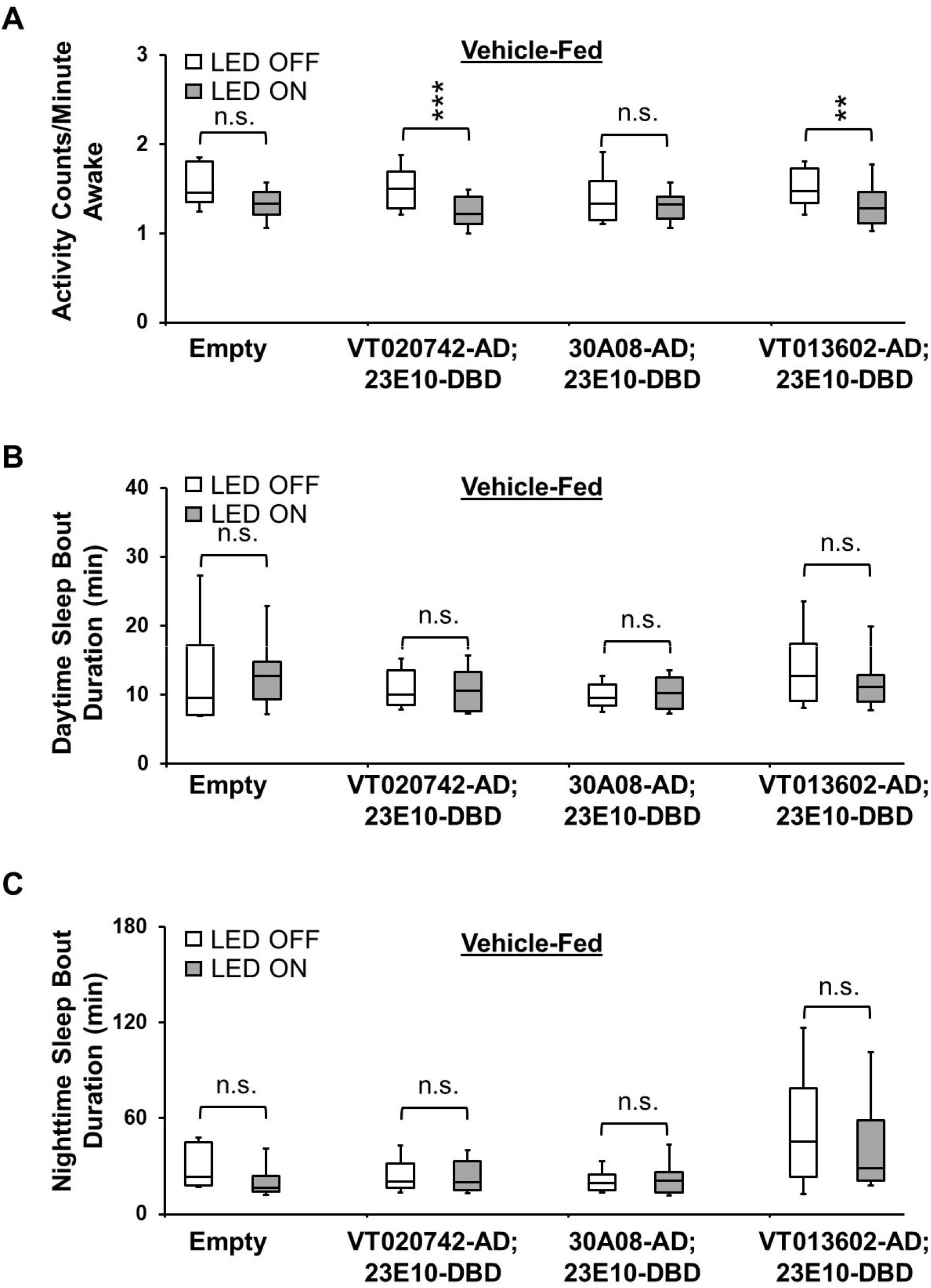
Vehicle-fed sleep data for females in optogenetic activation of lines containing only VNC neurons. **A)** Box plots of locomotor activity counts per minute awake for vehicle-fed flies presented in Figure 4K. The bottom and top of each box represents the first and third quartile, and the horizontal line dividing the box is the median. The whiskers represent the 10^th^ and 90^th^ percentiles. Two-way repeated measures ANOVA followed by Sidak’s multiple comparisons test found that locomotor activity per awake time is reduced in vehicle-fed *VT020742-AD; 23E10-DBD>UAS-CsChrimson* and *VT013602-AD; 23E10-DBD>UAS-CsChrimson* flies that are stimulated with 627nm LEDs. **P<0.01, ***P<0.001. n.s.= not significant. n=13-32 flies per genotype. **B)** Box plots of daytime sleep bout duration for vehicle-fed flies presented in Figure 4K. The bottom and top of each box represents the first and third quartile, and the horizontal line dividing the box is the median. The whiskers represent the 10^th^ and 90^th^ percentiles. Two-way repeated measures ANOVA followed by Sidak’s multiple comparisons test found no difference in daytime sleep bout duration when the flies are stimulated with 627nm LEDs. n.s= not significant. n=13-32 flies per genotype. **C)** Box plots of nighttime sleep bout duration for vehicle-fed flies presented in Figure 4K. The bottom and top of each box represents the first and third quartile, and the horizontal line dividing the box is the median. The whiskers represent the 10^th^ and 90^th^ percentiles. Two-way repeated measures ANOVA followed by Sidak’s multiple comparisons test show that nighttime sleep bout duration is not different when vehicle-fed flies are stimulated with 627nm LEDs. n.s.= not significant. n=13-32 flies per genotype.

**Figure 4-figure supplement 2:**
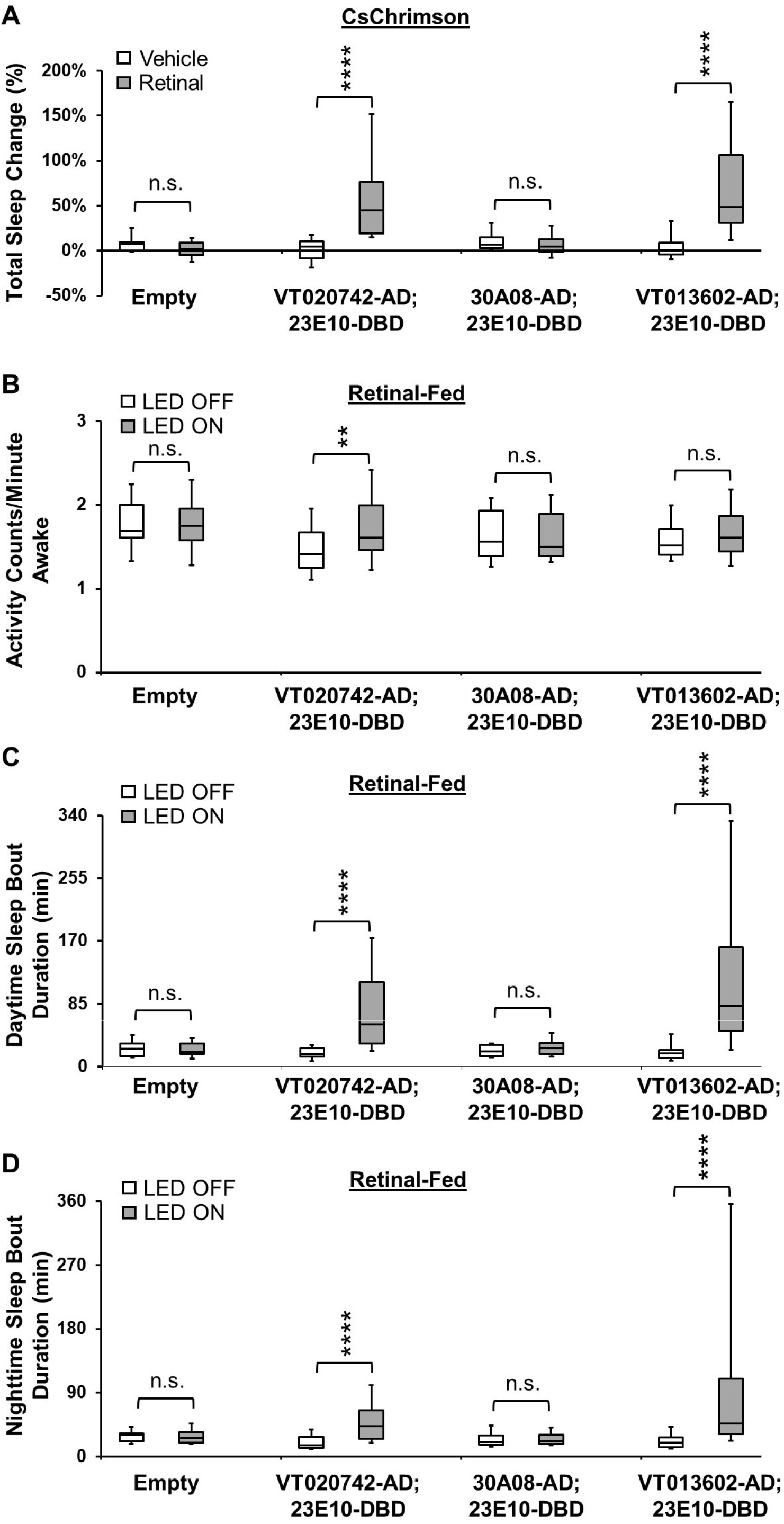
Optogenetic activation of lines containing only VNC neurons. Male data. **A)** Box plots of total sleep change in % for control (*Empty-AD; 23E10-DBD>UAS-CsChrimson*), *VT020742-AD; 23E10-DBD>UAS-CsChrimson*, *30A08-AD; 23E10-DBD>UAS-CsChrimson* and *VT013602-AD; 23E10-DBD>UAS-CsChrimson* vehicle-fed and retinal-fed male flies upon 627nm LED stimulation. The bottom and top of each box represents the first and third quartile, and the horizontal line dividing the box is the median. The whiskers represent the 10^th^ and 90^th^ percentiles. Two-way ANOVA followed by Sidak’s multiple comparisons revealed that retinal-fed *VT020742-AD; 23E10-DBD>UAS-CsChrimson* and *VT013602-AD; 23E10-DBD>UAS-CsChrimson* flies increase sleep significantly when stimulated with 627nm LEDs when compared with vehicle-fed flies. ****P<0.0001, n.s.= not significant. n=18-40 flies per genotype and condition. **B)** Box plots of locomotor activity counts per minute awake for retinal-fed flies presented in A. The bottom and top of each box represents the first and third quartile, and the horizontal line dividing the box is the median. The whiskers represent the 10^th^ and 90^th^ percentiles. Two-way repeated measures ANOVA followed by Sidak’s multiple comparisons test show that locomotor activity per awake time is increased in *VT020742-AD; 23E10-DBD>UAS-CsChrimson* flies and is not affected in the other genotypes when the flies are stimulated with 627nm LEDs. **P<0.05, n.s.= not significant. n=21-40 flies per genotype. **C)** Box plots of daytime sleep bout duration (in minutes) for retinal-fed flies presented in A. The bottom and top of each box represents the first and third quartile, and the horizontal line dividing the box is the median. The whiskers represent the 10^th^ and 90^th^ percentiles. Two-way repeated measures ANOVA followed by Sidak’s multiple comparisons indicates that daytime sleep bout duration is increased in retinal-fed *VT020742-AD; 23E10-DBD>UAS-CsChrimson* and *VT013602-AD; 23E10-DBD>UAS-CsChrimson* flies when stimulated with 627nm LEDs. ****P<0.0001, n.s.= not significant. n=21-40 flies per genotype. **D)** Box plots of nighttime sleep bout duration (in minutes) for retinal-fed flies presented in A. The bottom and top of each box represents the first and third quartile, and the horizontal line dividing the box is the median. The whiskers represent the 10^th^ and 90^th^ percentiles. Two-way repeated measures ANOVA followed by Sidak’s multiple comparisons indicates that nighttime sleep bout duration is increased in retinal-fed *VT020742-AD; 23E10-DBD>UAS-CsChrimson* and *VT013602-AD; 23E10-DBD>UAS-CsChrimson* flies when stimulated with 627nm LEDs. ****P<0.0001, n.s.= not significant. n=21-40 flies per genotype.

**Figure 4-figure supplement 3:**
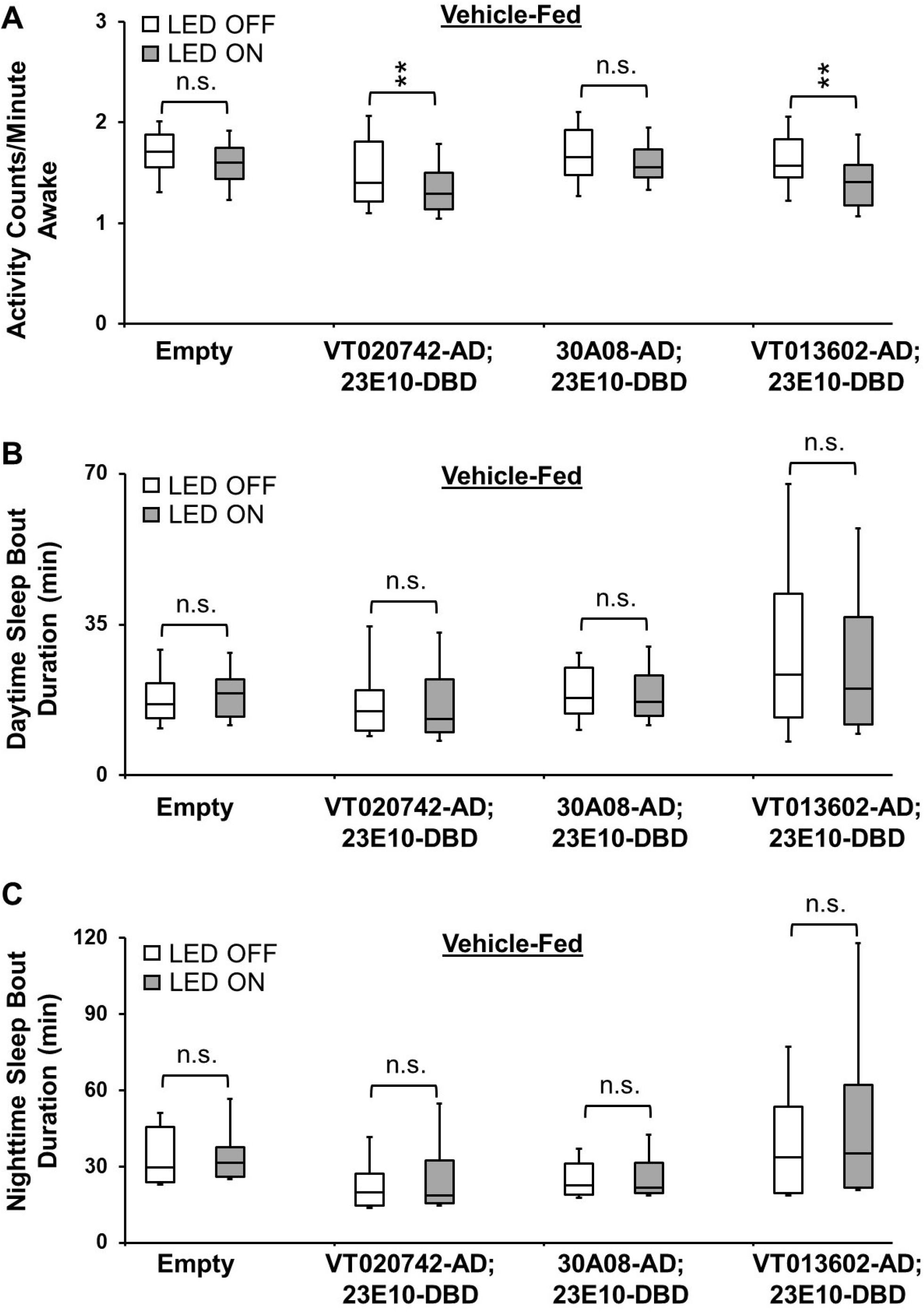
Vehicle-fed sleep data for males in optogenetic activation of lines containing only VNC neurons. **A)** Box plots of locomotor activity counts per minute awake for vehicle-fed flies presented in Figure 4-figure supplement 2A. The bottom and top of each box represents the first and third quartile, and the horizontal line dividing the box is the median. The whiskers represent the 10^th^ and 90^th^ percentiles. Two-way repeated measures ANOVA followed by Sidak’s multiple comparisons test found that locomotor activity per awake time is reduced in vehicle-fed *VT020742-AD; 23E10-DBD>UAS-CsChrimson* and *VT013602-AD; 23E10-DBD>UAS-CsChrimson* flies that are stimulated with 627nm LEDs. **P<0.01, n.s.= not significant. n=18-32 flies per genotype. **B)** Box plots of daytime sleep bout duration for vehicle-fed flies presented in Figure 4-figure supplement 2A. The bottom and top of each box represents the first and third quartile, and the horizontal line dividing the box is the median. The whiskers represent the 10^th^ and 90^th^ percentiles. Two-way repeated measures ANOVA followed by Sidak’s multiple comparisons test found no difference in daytime sleep bout duration when the flies are stimulated with 627nm LEDs. n.s= not significant. n=18-32 flies per genotype. **C)** Box plots of nighttime sleep bout duration for vehicle-fed flies presented in Figure 4-figure supplement 2A. The bottom and top of each box represents the first and third quartile, and the horizontal line dividing the box is the median. The whiskers represent the 10^th^ and 90^th^ percentiles. Two-way repeated measures ANOVA followed by Sidak’s multiple comparisons test show that nighttime sleep bout duration is not increased when vehicle-fed flies are stimulated with 627nm LEDs. n.s= not significant. n=18-32 flies per genotype.

**Figure 4-figure supplement 4:**
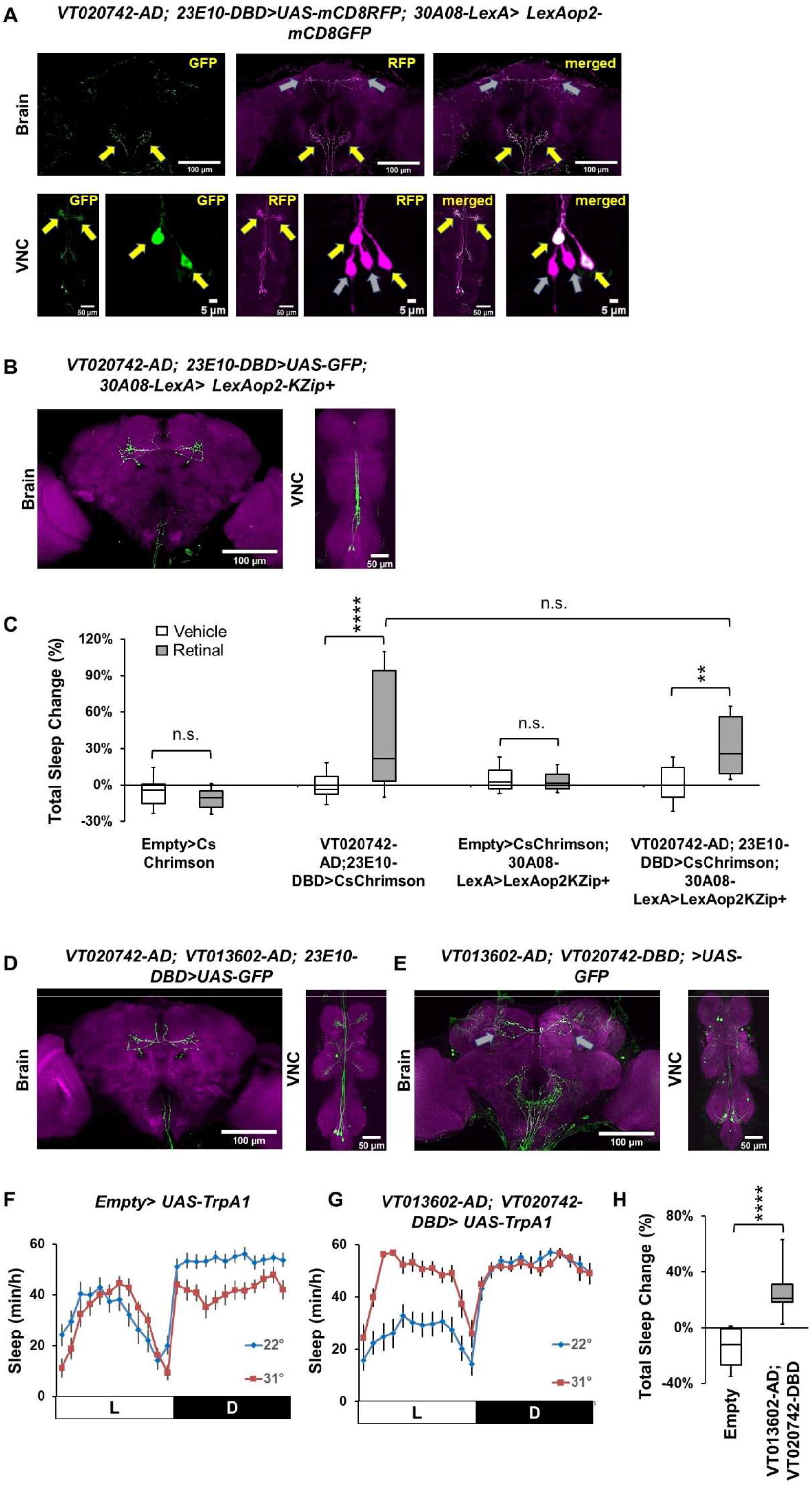
Additional data for TPN1 and VNC-SP neurons. **A)** Representative confocal stacks of a female *30A08-LexA>LexAop2-GFP; VT020742-AD; 23E10-DBD>UAS-RFP* fly showing the brain, the VNC and a magnified view of the cell bodies in the metathoracic ganglion. Yellow arrows show the TPN1 neurons and their processes in the VNC and in the brain and gray arrows the VNC-SP neurons and their processes. TPN1 neurons are labeled by 30A08-LexA and *VT020742-AD; 23E10-DBD* while VNC-SP neurons are only present in *VT020742-AD; 23E10-DBD*. Green, anti-GFP; magenta, anti-RFP. **B)** Representative confocal stacks of a female *30A08-LexA>LexAop2-KZip^+^; VT020742-AD; 23E10-DBD>UAS-GFP* fly showing the brain and the VNC. The KZip^+^ repressor effectively remove expression in the TPN1 neurons leaving only the VNC-SP neurons. Green, anti-GFP; magenta, anti-nc82 (neuropile marker). **C)** Box plots of total sleep change in % for *Empty-AD; 23E10-DBD>UAS-CsChrimson*, *VT020742-AD; 23E10-DBD>UAS-CsChrimson*, *Empty-AD; 23E10-DBD>UAS-CsChrimson*; *30A08-LexA> LexAop2-KZip^+^* and *VT020742-AD; 23E10-DBD>UAS-CsChrimson*; *30A08-LexA> LexAop2-KZip*^+^ vehicle-fed and retinal-fed female flies upon 627nm LED stimulation. The bottom and top of each box represents the first and third quartile, and the horizontal line dividing the box is the median. The whiskers represent the 10^th^ and 90^th^ percentiles. Two-way ANOVA followed by Sidak’s multiple comparisons revealed that retinal-fed *VT020742-AD; 23E10-DBD>UAS-CsChrimson* and *VT020742-AD; 23E10-DBD>UAS-CsChrimson*; *30A08-LexA> LexAop2-KZip^+^* flies increase sleep significantly when stimulated with 627nm LEDs when compared with vehicle-fed flies. Tukey’s multiple comparisons demonstrate that there is no difference in total sleep change between retinal-fed *VT020742-AD; 23E10-DBD>UAS-CsChrimson* and *VT020742-AD; 23E10-DBD>UAS-CsChrimson*; *30A08-LexA> LexAop2-KZip^+^* flies. **P<0.01, ****P<0.0001, n.s.= not significant. n=17-31 flies per genotype and condition. **D)** Representative confocal stacks of a female *VT020742-AD/ VT013602-AD; 23E10-DBD>UAS-GFP* fly showing the brain and the VNC. This line labels the two TPN1 cells and the two VNC-SP neurons. Green, anti-GFP; magenta, anti-nc82 (neuropile marker). **E)** Representative confocal stacks of a female *VT013602-AD; VT020742-DBD>UAS-GFP* fly showing the brain and the VNC. This line labels the two VNC-SP neurons and their processes in the brain (gray arrows). Green, anti-GFP; magenta, anti-nc82 (neuropile marker). **F)** Sleep profile of control female (*Empty-AD; Empty-DBD>UAS-TrpA1*) at 22°C (blue line) and 31°C (red line). **G)** Sleep profile of VT013602-AD; VT020742-DBD>UAS-TrpA1 female flies at 22°C (blue line) and 31°C (red line). **H)** Box plots of total sleep change in % for flies presented in F and G. The bottom and top of each box represents the first and third quartile, and the horizontal line dividing the box is the median. The whiskers represent the 10^th^ and 90^th^ percentiles. A two-tailed unpaired t test revealed that activating VT013602-AD; VT020742-DBD neurons significantly increases sleep compared with controls. ****P<0.0001. n=13-14 flies per genotype.

**Figure 4-figure supplement 5:**
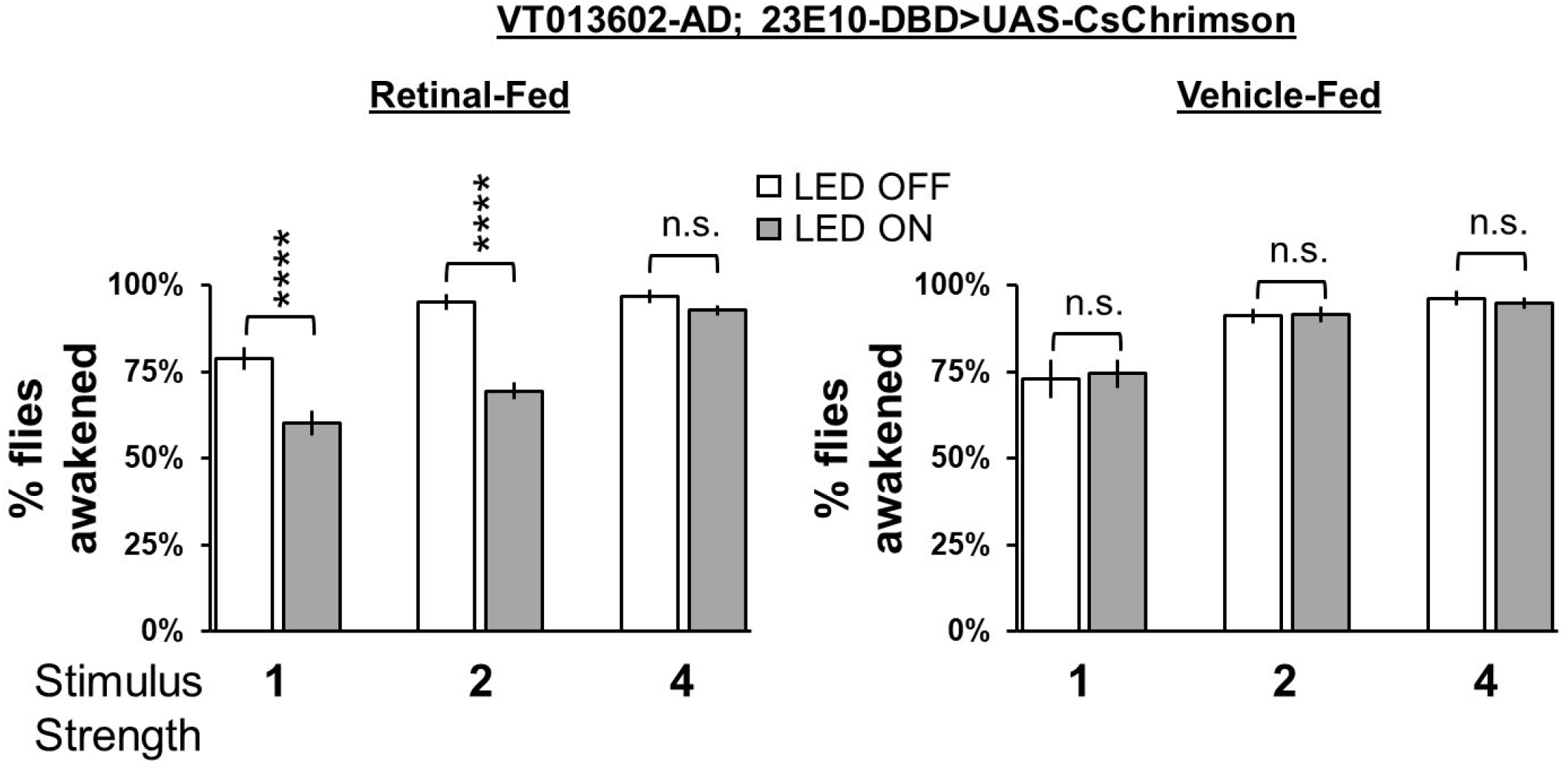
Arousal threshold for VNC-SP>UAS-CsChrimson flies Arousal threshold in vehicle-fed and retinal-fed *VT013602-AD; 23E10-DBD>UAS-CsChrimson* female flies. Percentage of flies awakened by a stimulus of increasing strength (1, 2 and 4 downward movements in the SNAP apparatus) with and without 627nm LEDs stimulation. Two-way ANOVA followed by Sidak’s multiple comparisons indicates that in retinal-fed flies activation of VT013602-AD; 23E10-DBD neurons reduce the responsiveness to the 1 and 2 stimulus strength when compared with non-activated flies. No difference in responsiveness is seen at the strongest stimulus (4). Two-way ANOVA followed by Sidak’s multiple comparisons indicates that in vehicle-fed flies no difference in responsiveness is seen between LED stimulated and non-stimulated flies. ****P<0.0001, n.s.= not significant. n=16 flies per genotype and condition.

**Figure 6-figure supplement 1:**
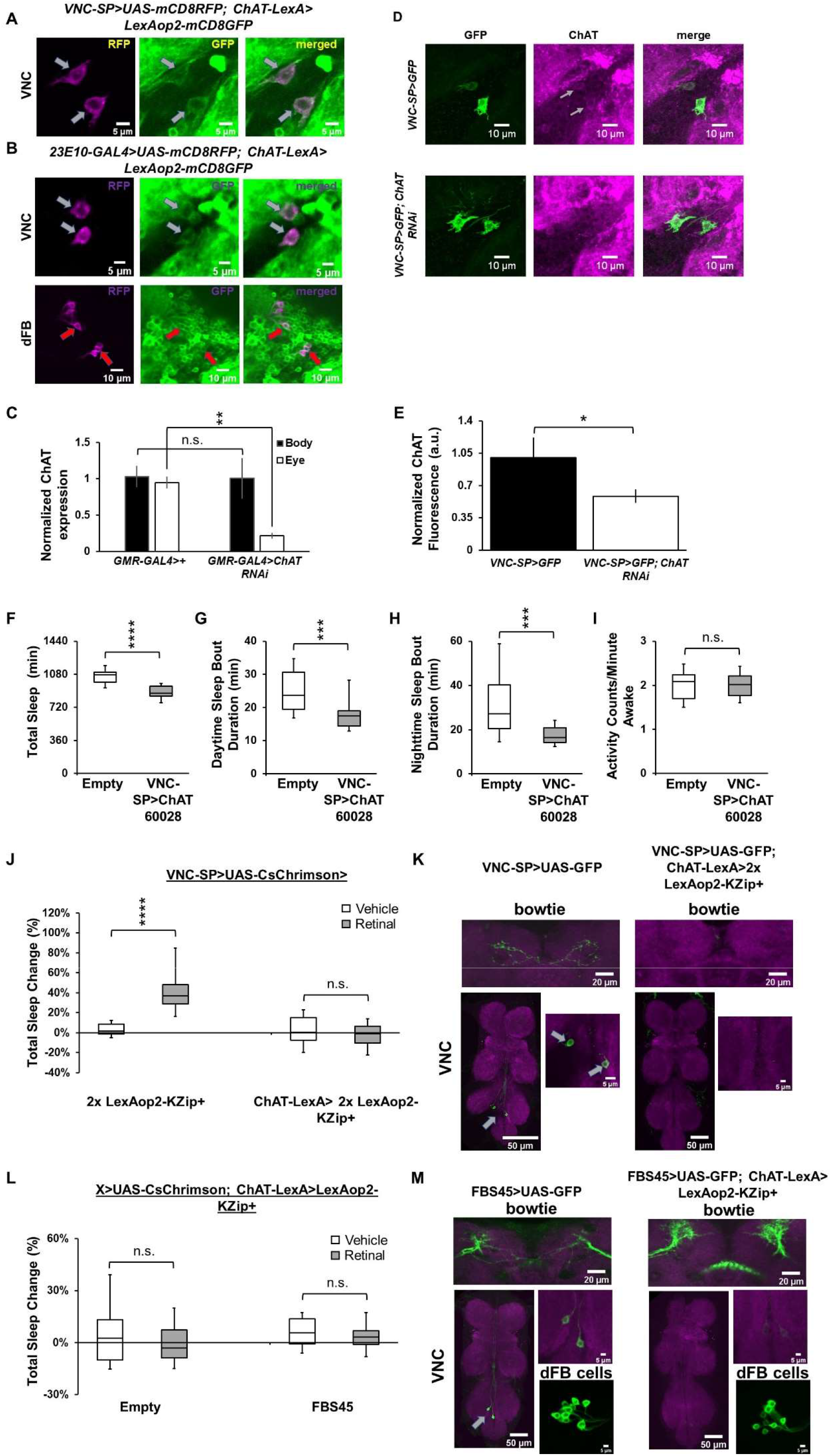
Additional data demonstrating that VNC-SP neurons are cholinergic. **A)** Representative confocal stacks of a female *ChAT-LexA>LexAop2-GFP; VT013602-AD; 23E10-DBD (VNC-SP)>UAS-RFP* focusing on the cell bodies in the metathoracic ganglion of the VNC. Gray arrows show the VNC-SP neurons that are co-labelled by the ChAT-LexA driver. Green, anti-GFP; magenta, anti-RFP. **B)** Representative confocal stacks of a female *ChAT-LexA>LexAop2-GFP; 23E10-GAL4>UAS-RFP*. Top panels, focusing on the cell bodies in the metathoracic ganglion of the VNC. Gray arrows show the VNC-SP neurons that are co-labelled by the ChAT-LexA driver. Bottom panels, focusing on dFB neurons. Red arrows show 23E10-GAL4 dFB neurons that are also labeled by ChAT-LexA. Green, anti-GFP; magenta, anti-RFP. **C)** Quantification of ChAT knockdown efficiency by RNAi. qPCR was performed on the body and eyes of control flies (*GMR-GAL4>+*) and flies expressing ChAT RNAi (line 60028) in the eyes driven by GMR-GAL4 (*GMR-GAL4>ChAT RNAi*). Expression levels were normalized to control levels. Two-way ANOVA followed by Sidak’s multiple comparisons revealed that in *GMR-GAL4>ChAT RNAi* flies ChAT levels are significantly reduced in the eyes but not in the body. **P<0.01. n.s.= not significant. n=3-4 replicates per genotype. **D)** ChAT immunostaining in *VNC-SP>UAS-GFP* (top panels) and *VNC-SP>UAS-GFP; UAS-ChAT RNAi* flies (bottom panels). Gray arrows point to neurons positive for GFP and ChAT. Green, anti-GFP; magenta, anti-ChAT. **E)** Quantification of data presented in D. A one-tailed unpaired t test revealed that expressing ChAT RNAi in VNC-SP neurons significantly reduce ChAT levels as measured with ChAT antibody staining. *P<0.05. n=14-17 VNC analyzed per genotype. **F)** Box plots of total sleep time (in minutes) for control (*Empty-AD; 23E10-DBD>ChAT 60028*) and *VNC-SP>ChAT 60028* male flies. The bottom and top of each box represents the first and third quartile, and the horizontal line dividing the box is the median. The whiskers represent the 10^th^ and 90^th^ percentiles. A two-tailed Mann-Whitney U test revealed that *VNC-SP> ChAT^RNAi^* male flies sleep significantly less than controls. ****P<0.0001. n=24-30 flies per genotype. **G)** Box plots of daytime sleep bout duration (in minutes) for control *(Empty-AD; 23E10-DBD>ChAT 60028)* and *VNC-SP>ChAT 60028* male flies. The bottom and top of each box represents the first and third quartile, and the horizontal line dividing the box is the median. The whiskers represent the 10^th^ and 90^th^ percentiles. A two-tailed Mann-Whitney U test revealed that *VNC-SP> ChAT^RNAi^* male flies daytime sleep bout duration is significantly reduced compared with controls. ***P<0.001. n=24-30 flies per genotype. **H)** Box plots of nighttime sleep bout duration (in minutes) for control *(Empty-AD; 23E10-DBD>ChAT 60028)* and *VNC-SP>ChAT 60028* male flies. The bottom and top of each box represents the first and third quartile, and the horizontal line dividing the box is the median. The whiskers represent the 10^th^ and 90^th^ percentiles. A two-tailed Mann-Whitney U test revealed that *VNC-SP> ChAT^RNAi^* male flies nighttime sleep bout duration is significantly reduced compared with controls. ***P<0.001. n=24-30 flies per genotype. **I)** Box plots of locomotor activity counts per minute awake for control *(Empty-AD; 23E10-DBD>ChAT 60028)* and *VNC-SP>ChAT 60028* male flies. The bottom and top of each box represents the first and third quartile, and the horizontal line dividing the box is the median. The whiskers represent the 10^th^ and 90^th^ percentiles. A two-tailed unpaired t test revealed that there is no difference in waking activity between *VNC-SP> ChAT^RNAi^* male flies and controls. n.s= not significant. n=24-30 flies per genotype. **J)** Box plots of total sleep change in % for *VT013602-AD; 23E10-DBD>UAS-CsChrimson*; 2x LexAop2KZip^+^ (control, no LexA driver) and for *VT013602-AD; 23E10-DBD>UAS-CsChrimson*; *ChAT-LexA>2x LexAop2KZip^+^* vehicle-fed and retinal-fed female flies upon 627nm LED stimulation. The bottom and top of each box represents the first and third quartile, and the horizontal line dividing the box is the median. The whiskers represent the 10^th^ and 90^th^ percentiles. Two-way ANOVA followed by Sidak’s multiple comparisons revealed that retinal-fed *VT013602-AD; 23E10-DBD>UAS-CsChrimson; ChAT-LexA>2x LexAop2KZip^+^* flies do not increase sleep compared with vehicle-fed flies when stimulated with 627nm LEDs. Control flies increase sleep significantly. ****P<0.0001, n.s.= not significant. n=16-27 flies per genotype and condition. **K)** Representative confocal stacks of *VT013602-AD; 23E10-DBD>UAS-GFP* (left) and *VT013602-AD; 23E10-DBD>UAS-GFP*; *ChAT-LexA>2x LexAop2KZip^+^* (right) female flies. Expression of GFP in VNC-SP neurons and in their bowtie brain processes is completely abolished by the expression of the KZip^+^ repressor. Gray arrows show VNC-SP neurons. Green, anti-GFP; magenta, anti-RFP. **L)** Box plots of total sleep change in % for *Empty-AD; 23E10-DBD>UAS-CsChrimson*; *ChAT-LexA> LexAop2KZip^+^* and for *FBS45>UAS-CsChrimson; ChAT-LexA> LexAop2KZip^+^* vehicle-fed and retinal-fed female flies upon 627nm LED stimulation. The bottom and top of each box represents the first and third quartile, and the horizontal line dividing the box is the median. The whiskers represent the 10^th^ and 90^th^ percentiles. Two-way ANOVA followed by Sidak’s multiple comparisons revealed no differences in sleep when flies are stimulated with 627nm LEDs. n.s.= not significant. n=22-29 flies per genotype and condition. **M)** Representative confocal stacks of *FBS45>UAS-GFP* (left) and *FBS45>UAS-GFP; ChAT-LexA>LexAop2KZip^+^* (right) female flies. Expression of GFP in VNC-SP neurons and in their bowtie brain processes is completely abolished by the expression of the KZip^+^ repressor. Gray arrows show VNC-SP neurons. In addition, the KZip^+^ repressor also consistently remove expression in 4-6 dFB neurons. On average 19.33 ± 0.62 dFB neurons are labeled in *FBS45>UAS-GFP; ChAT-LexA>LexAop2KZip^+^* while it is more than 25 without the KZip^+^ repressor (Table 2). Green, anti-GFP; magenta, anti-RFP.

**Figure 6-figure supplement 2:**
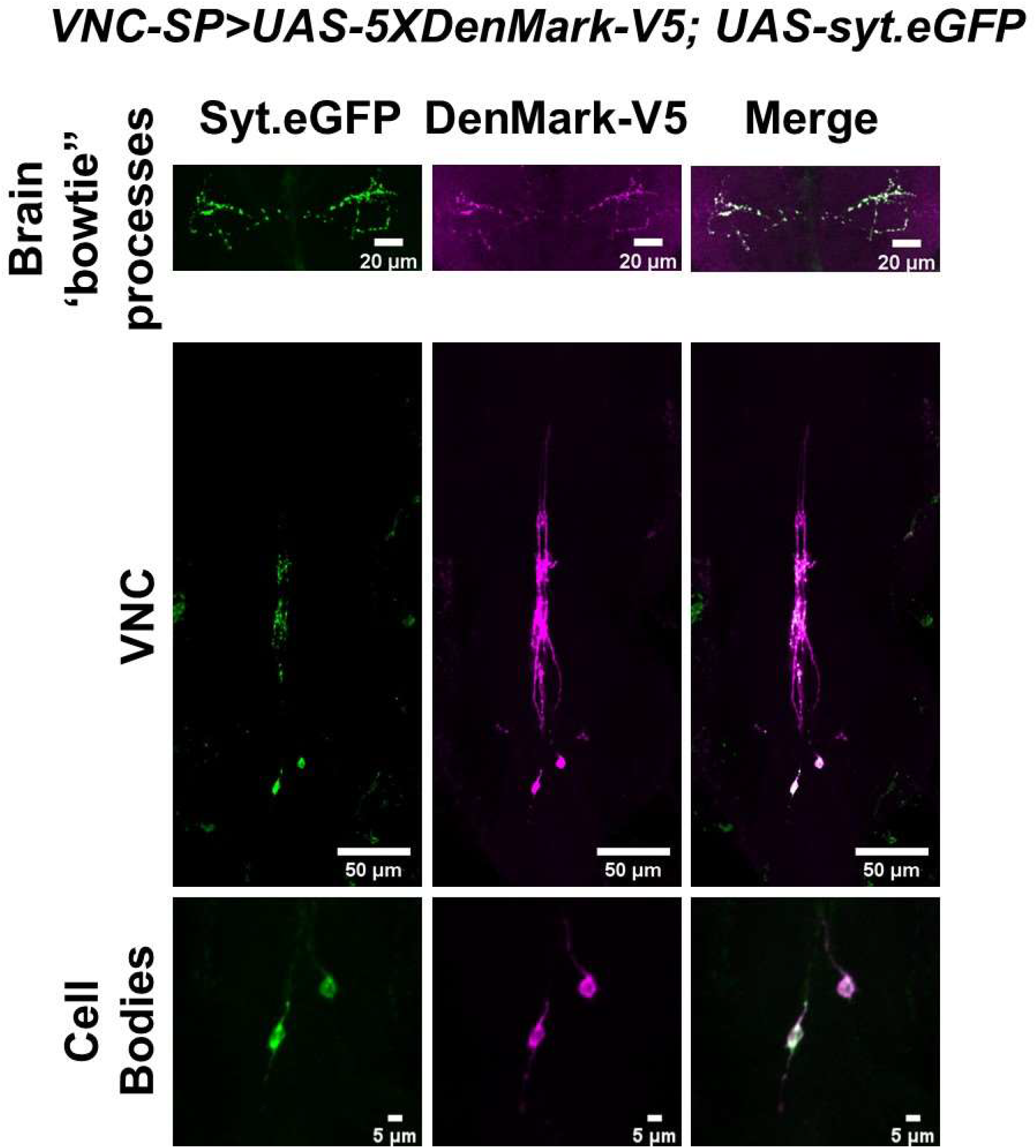
VNC-SP neurons are well positioned to integrate sensory inputs in the VNC and send this information to the brain. Representative confocal stacks of a female *VT013602-AD; 23E10-DBD> UAS-5xDenMark-V5; UAS-syt.eGFP*. Top panels, focusing on the brain “bowtie” processes. Middle panels, the VNC and bottom panels, a magnified view of the cell bodies area. Green, anti-GFP; magenta, anti-V5.

